# Single-cell chiral symmetry breaking under confinement

**DOI:** 10.64898/2025.12.23.696323

**Authors:** Sebastián Echeverría-Alar, Badri Narayanan Narasimhan, Stephanie I Fraley, Wouter-Jan Rappel

## Abstract

Single cells confined by the extracellular matrix can exhibit persistent rotational motion, yet the physical mechanisms underlying this chiral symmetry breaking remain unclear. Here, we address this gap with a cellular phase field model that couples cell deformation, cell polarization governed by stochastic excitable dynamics, and confinement. We identify the confinement strength as a bifurcation parameter determining three regimes: strong confinement prevents rotation through spatial constraints, intermediate confinement induces stochastic transitions between chiral and non-chiral states, and weak confinement allows persistent rotational motion. For the intermediate regime, we develop a semi-Markovian renewal process framework that characterizes the stochastic dynamics through dwell time statistics, transition probabilities and first-passage times. For the weak confinement regime, we reveal that a mechanochemical feedback enables coherent rotations despite internal noise through the reduction of local excitability mediated by mechanical contraction. We formalize this feedback analytically using Kramers escape theory. Experiments on epithelial MCF10A cells in Matrigel validate predictions for the weak confinement regime. Our results establish a theoretical approach for understanding single-cell chiral symmetry breaking under confinement, with implications for controlling single-cell dynamics by tuning extracellular matrix properties.

## I. INTRODUCTION

Chiral symmetry breaking, the spontaneous establishment of a preferred handedness, is a robust emergent phenomenon in macroscopic active systems. Its diverse dynamical consequences have been reported in fish schools [1], ant mills [2], *Dictyostelium discoideum* cell clusters [3], starfish embryos [4], pancreatic spheres [5], mosh pits [6], and in confined multicellular structures [7–9]. In these systems, chirality typically emerges from interactions among many constituents, but also from their respective interactions with the surroundings. In the case of cells in physiologically relevant microenvironments, e.g. the extracellular matrix (ECM), cell-ECM interactions start to play a fundamental role (see [10] and references therein). The ECM locally confines cells, reducing the available space for protrusion growth and migration. This confinement has the ability of forcing multiple cells to self-organize into chiral multi-cellular aggregates [7, 9], and this phenomenon is observed in other types of biological confinement as well, including micropatterned substrates [11–13].

Chirality can also arise at the level of a single constituent, when that unit is an out-of-equilibrium system. Eukaryotic cells provide an excellent example, because they exhibit a wide and diverse range of subcellular processes, including membrane deformations and membranebound biochemical signaling, that break temporal and spatial symmetries [14–16]. Two relevant processes in single-cell chirality are cell polarization [17, 18] and protrusion creation triggered by F-actin polymerization [19– 21]. In both, the appearance of well-defined temporal and spatial scales is governed by excitable wave-like dynamics of membrane-bound proteins at the cell cortex [22–27], which are maintained despite intrinsic noise [25, 28, 29]. Experiments have shown that transient rotational protrusions in single cells arise before eventually transitioning to a fully polarized mesenchymal mode of migration on 2D substrates [30, 31]. Moreover, experiments have demonstrated the possibility of maintaining a coherently rotating cell (MCF10A) in Matrigel [7], yet the physical mechanisms that originate and sustain such behavior have not been fully addressed. Intriguingly, single-cell rotation appears to be variable even within the same cell type under similar confinement conditions: while some MCF10A cells rotate [7], others do not [32]. This variability extends across cell types, as a recent study has reported the absence of rotational motion in single MDCK cells in Matrigel [33], suggesting that the phenomenon may involve intrinsic cellular dynamics with both deterministic and stochastic components.

A simple theoretical framework that integrates cell-ECM interactions, cell mechanics, and signaling dynamics to understand when and why single-cells under confinement rotate is currently unavailable. This framework needs to address key questions: What are the minimal mechanochemical ingredients governing the transition from a non-chiral to chiral cells? What mechanisms maintain rotational coherence against intrinsic noise?

Here we present a 3D phase field model, which couples cell membrane deformation, a polarization mechanism dictated by stochastic excitable dynamics, and homogeneous ECM confinement that robustly show the spontaneous chiral symmetry breaking of single cells. The confinement does not impose the chirality but rather establishes the necessary conditions to observe the spontaneous emergence of cell rotations. We find that the persistence of chiral cells depend on the level of confinement: under strong confinement, cell dynamics is governed by disorganized short-lived protrusions and the cell remains static, while under weak confinement, the cell is able to establish coherent rotations associated with longlived protrusions. The model also predicts an intermediate stochastic regime, governed by intermittent events of rotational coherence.

To describe the stochastic regime, we coarse-grain a two-dimensional representation of the excitable dynamics in our model into a discrete set of states and quantify the dynamics via mean dwell times, transition probabilities, and mean first passage times. We show in this discrete representation that transitions between non-chiral and chiral cells can be captured by a semi-Markov renewal process, providing a compact data-driven approach to analyze stochastic switching in excitable cellular dynamics. For the weak confinement regime, we identify a mechanochemical feedback mechanism: strong contractility induced by large protrusions reduces cortical excitability and, thus, suppresses the noise-triggered nucleation of new protrusions. We formalize this mechanism analytically based on Kramers escape theory: the persistent chiral motion can be understood as a state trapped in a potential well with a contractility-driven barrier.

Finally, we conduct multiple experimental realizations of single epithelial MCF10A cells seeded in Matrigel across a range of concentrations (30%-100%). Comparisons with our theory suggest that MCF10A in Matrigel exhibit dynamics consistent with the weak confinement regime. Therefore, our work provides a theoretical framework for understanding single-cell chiral symmetry breaking under confinement, experimental support for the weak confinement regime, and a foundation for future experimental studies exploring the stochastic and strong confinement regimes.

## II. RESULTS

### A. Rotational dynamics under confinement

The phase field approach is a flexible continuum method that can couple boundary deformations to the spatiotemporal evolution of bulk scalar fields. It has successfully captured cell morphodynamics [34], cell-substrate adhesion [35], cell interactions [12], cell plasticity [20, 36], and collective cell migration [37]. Specifically, we consider a scalar field *φ*_*c*_(**r**, *t*) to represent the cell position: *φ*_*c*_ = 1 inside the cell, *φ*_*c*_ = 0 outside the cell, and *φ*_*c*_ = 1*/*2 corresponding to the interface. A second static phase field, *φ*_*M*_ (**r**), represents the confining ECM (see Fig. S1A). The cell interacts with the ECM through a repulsive force 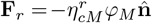, where 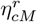 is a parameter that quantifies confinement and 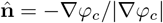 is the outward unit normal on the cell’s interface. An additional friction force mediates the interaction between the cell and the ECM, **F**_*f*_ = −*ξφ*_*M*_ **v**, where *ξ* is a friction coefficient and **v** is the velocity of the cell interface. This friction term phenomenologically encompasses hydrodynamic drag, viscous dissipation from ECM deformation, and dissipation from integrin-based adhesive interactions. Cell size, *V*_*c*_, is maintained by a size-restoring pressure force: 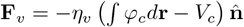, where *η*_*v*_ controls the strength of this constraint. This force produces nonlocal mechanical coupling: local deformations induce compensating membrane changes throughout the cell. Membrane tension, **F**_*t*_, is modeled through an interfacial energy with surface surface tension coefficient *γ*, which resists membrane deformations (see Supplemental Material and Fig. S1B for details [38]). The cell is driven out-of-equilibrium with a protrusion force 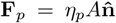 [12, 34], parameterized by *η*_*p*_, where the activator *A* is the mean-field representation of a membrane-bound protein that controls protrusion growth. The activator is coupled to the evolution of a slow membrane-bound inhibitor *R*, and the two-variable system obeys stochastic excitable dynamics:

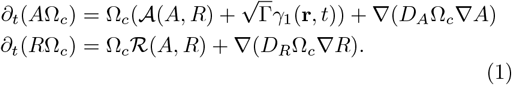

The functionals 𝒜 and ℛ include commonly used positive and negative feedbacks (detailed in the Supplemental Material [38]) between the activator and inhibitor species [20, 25, 36]. The excitable dynamics follows from a single intersection in the phase space, defined by the homogeneous solution {*A*_*o*_, *R*_*o*_}, between the linear and cubic nullcline of the activator-inhibitor system (Fig. S1C). The functional Ω_*c*_ = Ω_*c*_(*φ*_*c*_) is an indicator of the cell position (see Supplemental Material for details [38]), and it couples cell membrane deformations to the signaling dynamics. Both species diffuse with diffusivities *D*_*A*_ and *D*_*R*_, respectively. Stochasticity is introduced as Gaussian noise *γ*_1_(**r**, *t*) with amplitude Γ, a common approximation for systems where fluctuations emerge from many independent subcellular processes [25, 28, 29, 39, 40]. Cell motion is governed by the advection equation

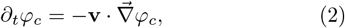

where the velocity is obtained from a force balance at the cell interface that includes all forces described above and is evaluated in an overdamped limit. Consistent with this approximation, we assume that translational cell dynamics being much slower than cell membrane relaxation [41].

We numerically integrate the phase-field model (Eq. 2) from a spherical initial condition corresponding to the resting state {*A*_*s*_, *R*_*s*_} (parameters in Table S1; numerical details in the Supplemental Material [38]). Varying the confinement parameter 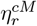 produces three qualitatively distinct single-cell dynamical regimes (Videos S1–S3). Parameter values are presented unitless in the main text for readability; the associated physical units are provided in Table S1 (Supplemental Material [38]). Under strong confinement 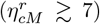, the cell is jammed with an approximately spherical shape, a noisy activator background, and a centroid that remains stationary on average (Fig. 1Ai-ii). To visualize activator dynamics, we analyze its behavior on an arbitrary plane 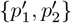, which passes through the centroid. This shows that short-lived and small-amplitude protrusions appear, but are not able to produce organized dynamics (Video S4 and Fig. 1Aiii).

**FIG. 1.**
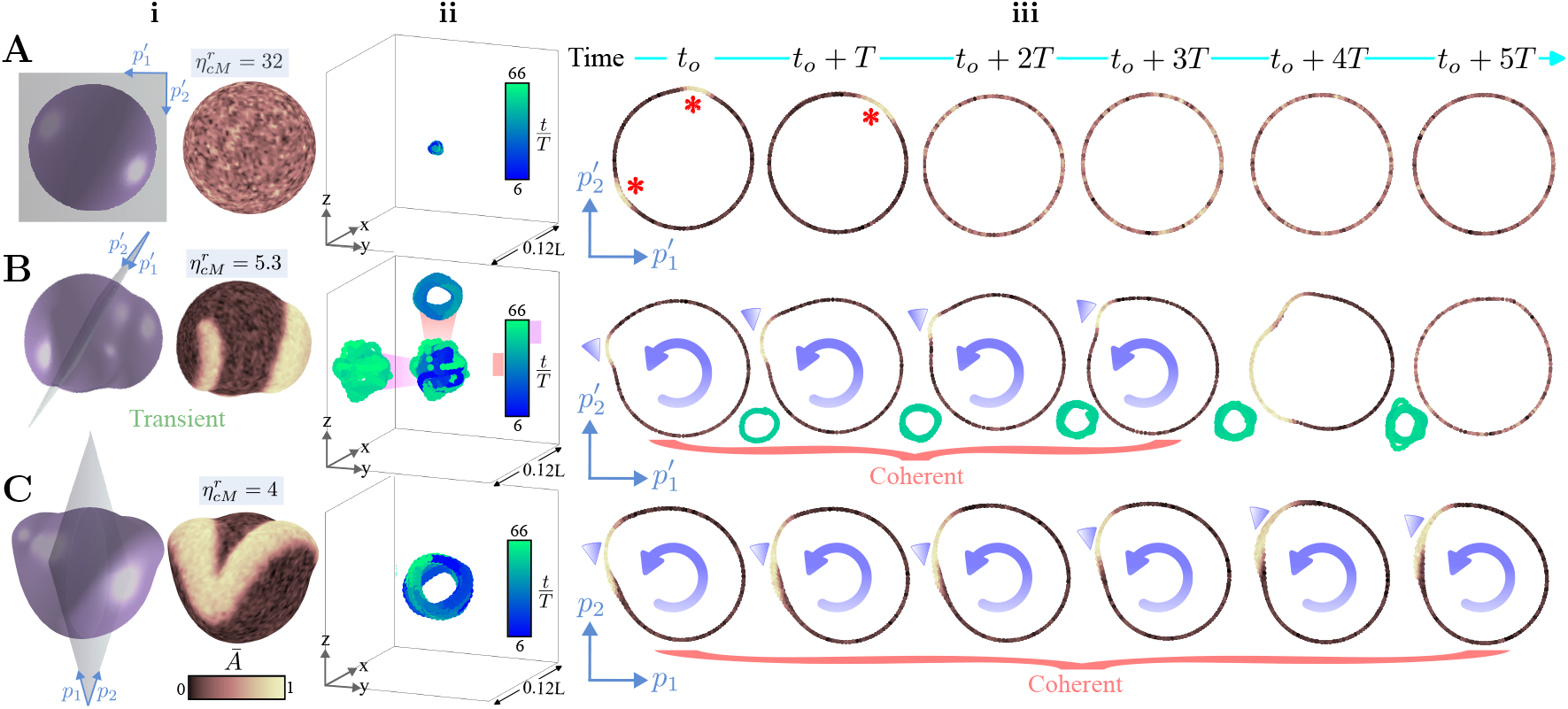
Chiral symmetry breaking under confinement in the 3D phase field model. (A) Cell under high confinement conditions 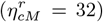, (B) cell in the stochastic regime with 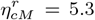, and (C) deterministic rotating cell 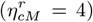. For all the dynamical regimes, the panels show: (i) cell shape (*φ*_*c*_ = 1*/*2) and normalized activator field *Ā* = *A/* max(*A*), (ii) trajectories of the centroid of the cells over 60*T*, and (iii) activator distributions on planes 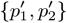 (A and B) or {*p*_1_, *p*_2_} (C). The red (*) symbols in (Aiii) indicate short-lived protrusions. In (Biii) and (Ciii), the blue arrows highlight the coherent rotational motion and the blue triangles indicate the position of the rotating protrusions. The trajectories in (Biii) illustrate the connection between cell centroid motion and cell shape deformation during the loss of rotational coherence. The time variables *T* (ii-iii) and *t*_*o*_ (iii) correspond to the period of the rotational motion at 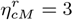 and an arbitrary initial time, respectively.

At intermediate confinement 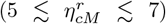, the single-cell dynamics becomes complex: sufficiently large protrusions form and trigger rich spatiotemporal activator patterns, including spiral waves (Video S2). Fig. 1Bi illustrates the case 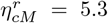, where a transient cell shape and activator field are depicted. The activator shows a spiral wave fragment (left) and the early stage of a protrusion (right). The protrusion, correlated with a local accumulation of the activator, interacts with the ECM and is propagated as a target wave along the surface. Stochastic fluctuations or heterogeneities in refractoriness can break this pulse (see Figs. S2 and S3 in the Supplemental Material [38]), leading to the formation of spiral waves. The resulting centroid trajectory contains localized periods of coherent rotations (shaded red) with the motion governed by two spiral wave fragments synchronously rotating along the cell surface (Video S2 and Fig. 1Bii). The rotational coherence can be lost after some time (shaded pink). Choosing a visualization plane that encompasses the centroid positions during the coherent rotation reveals a single “pulse” that circulates along the periphery until its disappearance (Video S5 and Fig. 1Biii). Eventually, the centroid trajectory exhibits coherent rotations again, but on a different plane (Videos S2 and S5).

For weak confinement 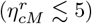, the cell enters a permanently rotating state after an initial transient (Fig.1C and Video S3). The shape adopts a symmetric two-lobe geometry correlated with a frustrated “figure of eight” activator pattern (Fig.1Ci and Fig. S4A in the Supplemental Material [38]). The centroid trajectory is almost fully bounded to the symmetry plane defined by the two-lobe geometry, {*p*_1_, *p*_2_}, where a permanently rotating pulse is observed (Video S6 and Fig. 1Cii-iii). Since the rotational period of the pulse is established by the activator-inhibitor dynamics (Eq. 1), we can define a unique time scale, chosen here to be the period of the rotating state for *η*^*r*^ = 3: *T*. We note that this rotational period can be redefined by rescaling time and the right-hand side in Eqs. (1) and (2). Additional centroid trajectories in the confinement range 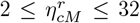 are shown in Fig. S5 (Supplemental Material [38]).

Finally, for confinement values below 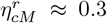, a single protrusion quickly polarizes the cell, inducing a permanent unidirectional motion rather than rotational dynamics. This unconfined regime is characterized by effective cell penetration into the ECM, where confinement no longer governs the dynamics, and thus, we restrict our study to 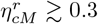. Together, these results demonstrate that signatures of rotation are observed within a finite confinement window and the maintenance of such dynamics is stochastic or purely deterministic depending on the value of 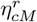.

### B. Discrete state representation captures stochastic transitions

To systematically investigate the emergence and maintenance of rotating dynamics, and motivated by the simpler picture of pulse-dynamics in a plane (Fig. 1), we next reduce our 3D model to a 2D phase field model (see Section V in the Supplemental Material for details [38]). Importantly, in this reduced description, coherent and non-coherent rotations are still observed. We note that while the three confinement regimes persist in 2D, the range of the intermediate regime is larger, 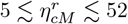, presumably because rotational waves nucleate more readily in the 2D case, where waves propagate along a 1D boundary rather than a 2D surface. Our simulations reveal ten possible 2D activator patterns across confinements, 𝒮 = {𝒮_*i*_} _*i*=1,..,10_, shown in Fig. 2A (see also Video S7). We take advantage of the small discrete number of possible *states*, and decide to study the continuous 2D dynamics from a simpler, discrete perspective. This discrete description allows us to use established concepts in stochastic processes and statistical physics (see below), captures the switch between chiral and non-chiral behaviors while removing irrelevant microscopic details, and places confined cell dynamics in a broader theoretical framework. In this framework, the dynamics is governed by quasi-deterministic and stochastic transitions between the states, and their respective waiting times before transitioning (dwell times) 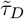, where the superscript indicates normalization by the rotational period *T*.

**FIG. 2.**
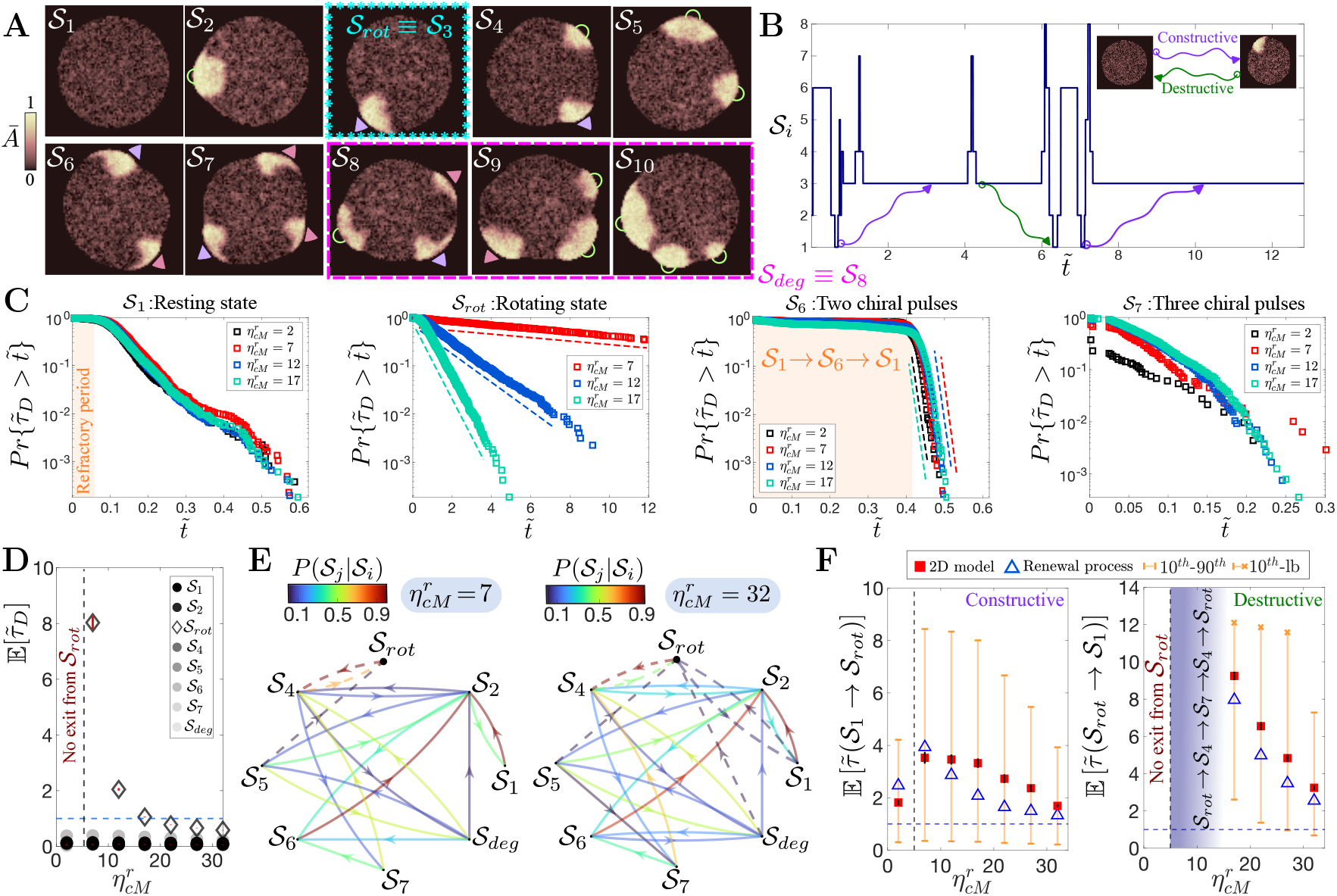
Emergence and maintenance of rotating states in-space. (A) Normalized activator fields from 2D simulations of the phase field model illustrating the ten states 𝒮_*i*_ considered in the analysis. The triangles indicate rotating protrusions while the half ring illustrate non-rotating protrusions. (B) Example of a trajectory of the low-dimensional model highlighting the birth and death of a rotating state: 𝒮_*rot*_. The time has been normalized by the period of the rotating state: 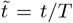. (C) Survival probabilities on a semi-log scale for the resting state (𝒮_1_), rotating state (𝒮_*rot*_), two chiral pulse state (𝒮_6_) and three chiral pulse state (𝒮_7_) for different degrees of confinement. The dashed lines indicate exponential decay for the corresponding cases. (D) Mean dwell times of the discrete states for different confinement values. The horizontal dashed line highlights one rotation period, and the vertical dashed line is 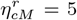. Bootstrap standard errors (500 trajectory resamples) of 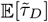 are indicated by solid red lines. (E) Transition probabilities for two different confinement values. The transitions involving 𝒮_*rot*_ are dashed. (F) Mean first passage time from the resting to the rotating state and vice-versa, computed using the 2D phase field model (squares) and the semi-Markov renewal process (triangles) as a function of 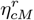.The orange whiskers correspond to the 10^*th*^ and 90^*th*^ percentiles of the first-passage time distributions. The orange (x) symbols indicates the lower bound (maximum first passage time observed) when a 90^*th*^ cannot be estimated (Supplemental Material [38]). Bootstrap standard errors (500 trajectory resamples) of both 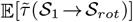 and 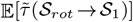 are indicated by solid red lines.

The states are defined as follows: 𝒮_1_ is the resting fixed point of the excitable system (Fig. S1C), _2_ is a non-chiral protrusion, and 𝒮_3_ (or 𝒮_*rot*_) is a rotational protrusion of either handedness, corresponding to the coherent rotational motion in the system. The states 𝒮_4_, 𝒮_5_, 𝒮_6_ are two-protrusion configurations: one chiral and one non-chiral, two non-chiral, and two chiral of opposite handedness, respectively. The last four, 𝒮_7_ − 𝒮_10_, are three-protrusion states: (i) three chiral protrusions, with two sharing the same handedness; (ii) two chiral protrusions of opposite handedness plus one non-chiral; (iii) two non-chiral and one chiral; and (iv) three nonchiral, respectively.

The numerical procedure to coarse-grain the 2D data into 𝒮-space is based on counting activator pulses on the cell periphery, assigning an angular displacement to distinguish between chiral and non-chiral pulses, and defining a time scale separation to account for extreme events of short duration (Figs. S6, S7, and S8 in the Supplemental Material [38]). A key simplification involves col-lapsing the states {𝒮_8_, 𝒮_9_, 𝒮_10_}into a single *degenerated* state, defined as 𝒮_*deg*_ (or the new 𝒮_8_). This is a remedy for the otherwise difficult task of introducing a time scale separation for these three states, since their durations are governed by the same underlying mechanism, a single, non-chiral pulse dynamics. Fig. 2B shows a sample trajectory in *𝒮*-space for 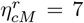, highlighting the creation (𝒮_1_ → 𝒮_*rot*_) and destruction (𝒮_*rot*_ → 𝒮_1_) of a coherent rotation (see exemplary trajectories for other confinements in Fig. S7 in the Supplemental Material [38]). Additionally, this trajectory exemplifies some of the quasi-deterministic transitions: division of a non-chiral protrusion into two counter-propagating pulses (𝒮_2_ → 𝒮_6_), and merging of the two counter-propagating pulses into a non-chiral pulse and its posterior disappearance (𝒮_6_ →𝒮_2_ →𝒮_1_).

We exploit the discrete-state description to extract robust statistics (Fig. S9 in the Supplemental Material [38]) and to quantify the dynamics of each state across different confinements, using trajectories of duration (13*T*). While this duration captures multiple transitions when sufficiently far from the critical confinement 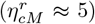, close to it, trajectories can become right-censored. We consider these censored trajectories in our analysis (see Supplemental Material [38]), as they provide valuable information near the critical confinement 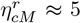, where the dynamics exhibit longer episodes of rotation. A useful statistical measure to distinguish between different states and confinement regimes is the survival probability of a state, 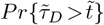 (Fig. 2C). In the weak and intermediate confinement regimes, the survival curves show that the system spends most of the time in the rotating state, while visits to other states are shorter than one rotation (for survival curves of additional states, see Fig. S10 in the Supplemental Material [38]). In the regime of permanent rotations (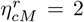 in Fig. 2C), no survival curve is computed for state _*rot*_ as all the trajectories are right-censored, i.e., once the system enters _*rot*_, it never leaves within the trajectory duration. The survival curves for the resting and the two-chiral pulse state exhibit quasi-deterministic waiting times (plateaus) before transitioning. The plateau in 𝒮_1_ persists for all confinements values, pointing to a deterministic time scale imposed by the excitable activator-inhibitor dynamics, the refractory period. The plateau of around half a rotation in 𝒮_6_ corresponds to the creation and annihilation of the two pulses with opposite handedness, i.e., the time spent in 𝒮_6_ within the passage 𝒮_1_ → 𝒮_6_ → 𝒮_1_, which represents the low-dimensional version of target wave propagation in 3D. Noisy activations increasingly perturb this deterministic motion as confinement increases, and the plateaus are replaced by decaying segments (Fig. S10 [38]). The survival curves for the state 𝒮_4_ display short plateaus when 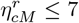 while for 𝒮_5_ there is only a short plateau when 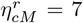 (Fig. S10 [38]). This is likely associated with the timescale of appearance and disappearance of a non-chiral pulse when another pulse is present.

The decay of the survival curves give further insights into the processes underlying the stochastic dynamics. Most states have a complex decay law, which do not follow a single exponential (see also Fig. S10), indicating the presence of multiple timescales rather than simple memory-less dynamics [42]. However, state 𝒮_6_, after its initial plateau, and state 𝒮_*rot*_ display a clear exponential decay across confinements, indicative of memory-less behavior. As confinement weakens toward *η*_*cM*_ ≃ 5, the survival curve of the rotating state exhibits a pronounced slowdown. At *η*_*cM*_ = 7 and *η*_*cM*_ = 12, the survival probability has not decayed to zero within the observation window (Fig. 2C, 𝒮_rot_), corresponding to a considerable fraction of right-censored trajectories (59% and 18%; Fig. S9, respectively).

We quantify state-dependent persistence by estimating the mean dwell time 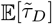 for all states in confinement conditions where it is well-defined (Fig. 2D and Section VI in the Supplemental Material [38]). Notice that because some trajectories are right-censored, 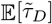 is a lower bound on the true mean. Nevertheless, this measurement clearly illustrates that the residence time in the rotating state increases as confinement is relaxed, and this motion becomes permanent when 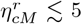 (no exit from 𝒮_*rot*_). We have verified that the persistence in 𝒮_*rot*_ is robust against variations in the amplitude of the noise (Fig. S11 in the Supplemental Material [38]).

The dwell time statistics can be complemented with the calculation of the transition probabilities *P*(𝒮_*j*_ |𝒮_*i*_) between states (details in Section VI in the Supplemental Material [38]). The transition probability graphs in Fig. 2E illustrate several characteristics of the pulse-dynamics. First, not all states are directly connected to each other due to the underlying deterministic dynamics. For example, the transitions 𝒮_2_ ↔ 𝒮_*rot*_ and 𝒮_4_ ↔ 𝒮_6_ are forbidden because of the topological reciprocal transformation between a non-chiral pulse and two pulses of opposite chirality. Second, some direct transitions are suppressed while others become more likely when the confinement is weakened (see also Fig. S12 in the Supplemental Material [38]). Specifically, when the confinement decreases from 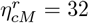 to 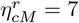, the direct passages 𝒮_*rot*_ → 𝒮_1_and 𝒮_*deg*_ → 𝒮_*rot*_ disappear, and the quasi-deterministic transition probabilities 𝒮_6_ → 𝒮_2_, 𝒮_4_ → 𝒮_3_ become larger.

The transition probabilities and dwell time statistics characterize only the local behavior of the discrete dynamics. To have a global picture of the process, we study first-passage times 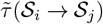, from a starting state _*I*_ to a target state _*j*_, focusing on the constructive and destructive pathways that drive the resting state to rotate (𝒮_1_ → 𝒮_*rot*_) and return to the resting state (𝒮_*rot*_ → 𝒮_1_). The left panel of Fig. 2F shows the mean first-passage time 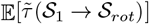 of creation, which is shorter than four rotation periods for all confinement parameters and follows a non-monotonic trend. It also reveals that, in the weak confinement regime 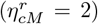, the establishment of permanent rotations is relatively fast. The right panel displays the destructive mean first-passage time 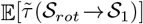, exhibiting an increasing trend as confinement weakens. Notice that for confinements 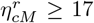, 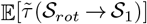 is approximately four times larger than the corresponding mean dwell time in *𝒮*_*rot*_ (Fig. 2D). This observation, combined with short mean dwell times in states *𝒮*_*i?*=*rot*_, suggests that the destruction path is on average an ensemble of multiple visits to *𝒮*_*rot*_. This behavior is determined by *P*(*𝒮*_4_|*𝒮*_*rot*_), the only possible exit from *𝒮*_*rot*_ (aside from a small transition probability to *𝒮*_1_ at 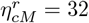), which then triggers the quasi-deterministic sequence: *𝒮*_*rot*_ → *𝒮*_4_ → *𝒮*_7_ → *𝒮*_4_→ *𝒮*_*rot*_. This sequence becomes increasingly deterministic as confinement weakens, and when 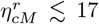, the right-censoring in the distribution of the first passage times 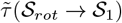 is larger than 77% (Fig. S13 in the Supplemental Material [38]). In these cases, indicated by shaded regions in Fig. 2F and Fig. S14 in the Supplemental Material [38], we are not able to compute the median (50^*th*^ percentile) of the distribution and we do not calculate the corresponding mean first passage times.

To test whether computable mean first-passage times can be inferred from the local dynamics, we adopt a datadriven semi-Markov renewal process (see Section VII in the Supplemental Material [38]). This framework accounts for arbitrary, state-dependent dwell time distributions, as several distributions deviate from exponential behavior (Fig. 2C and Fig. S10 [38]), and is characterized by the following modeled mean first passage times (renewal equation) [43]:

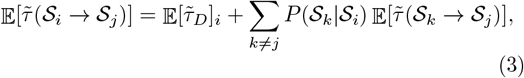

where 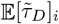 indicates the mean dwell time in state *𝒮*_*i*_. Using Eq. (3), we successfully capture the constructive and destructive paths, as well as the other mean first-passage times from *𝒮*_*rot*_ (Fig. S14 in the Supplemental Material [38]) across almost all confinement values, despite right-censoring in the distribution. The differences between the empirical and modeled mean first-passage times are smaller than 2*T*, except for the cases 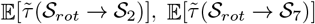 and 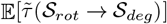 at 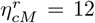, where is around 5*T*. We note that when the right-censoring in the first passage time distributions is less than 25%, the match between the empirical and modeled expectation values is perfect (e.g. *𝒮*_*rot*_ → *𝒮*_4_; Fig. S14 in the Supplemental Material [38]).

### C. Mechanochemical feedback stabilizes persistent rotation

Three features suggest that the stochastic behavior of the excitable activator-inhibitor dynamics is controlled by confinement: 1) the persistent coherent rotations for sufficiently weak confinements 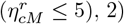 the increase in the probability of quasi-deterministic transitions when weakening the confinement, and 3) the dominance of a single sequence of states close to 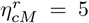. Although stochastic activations can orchestrate the constructive path, they become suppressed once cell rotation is established in weak confinement regimes. This hints to a mechanochemical feedback, which decreases the ability to nucleate new activator pulses (or protrusions) when the rotating pulse is present. To probe this feedback, we focus on the influence of the rotating protrusion on the rest of the cell membrane. Specifically, we concentrate on studying locally the opposite side of the cell, which we term the *back* (Fig. 3A), and define the variable *φ*_*b*_: a zero-dimensional representation of the cell phase field at the back (Section IX in the Supplemental Material [38]). The protrusion deformation induces a back contraction (∂_*t*_*φ*_*b*_ ≈ *v*_*n*_ |∂_*n*_*φ*_*b*_| *<* 0) via membrane tension and the size-restoring force, where the subscript *n* indicates the normal direction at the interface. Thus, larger protrusions, allowed under weak confinement and characterized by a broader activator pattern—quantified by 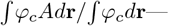trigger stronger back contractions (Fig. 3A). Importantly, these contractions in the weak confinement regime are strong enough to rotate the entire cell and not just the activator pulse (Fig. S15 in the Supplemental Material [38]). The strength of the contractile response can be tuned by the nonlocal mechanical force: reducing *η*_*v*_ decreases the back contraction magnitude across confinements (Fig. S16 in the Supplemental Material [38]). In addition, reductions in *η*_*v*_ affect the coherent rotational motion of the cell, as reflected by shorter mean dwell times in *𝒮*_*rot*_, even when the confinement is weak (Fig. 3B). These observations demonstrate that strong nonlocal mechanical contraction is essential for persistent rotation.

**FIG. 3.**
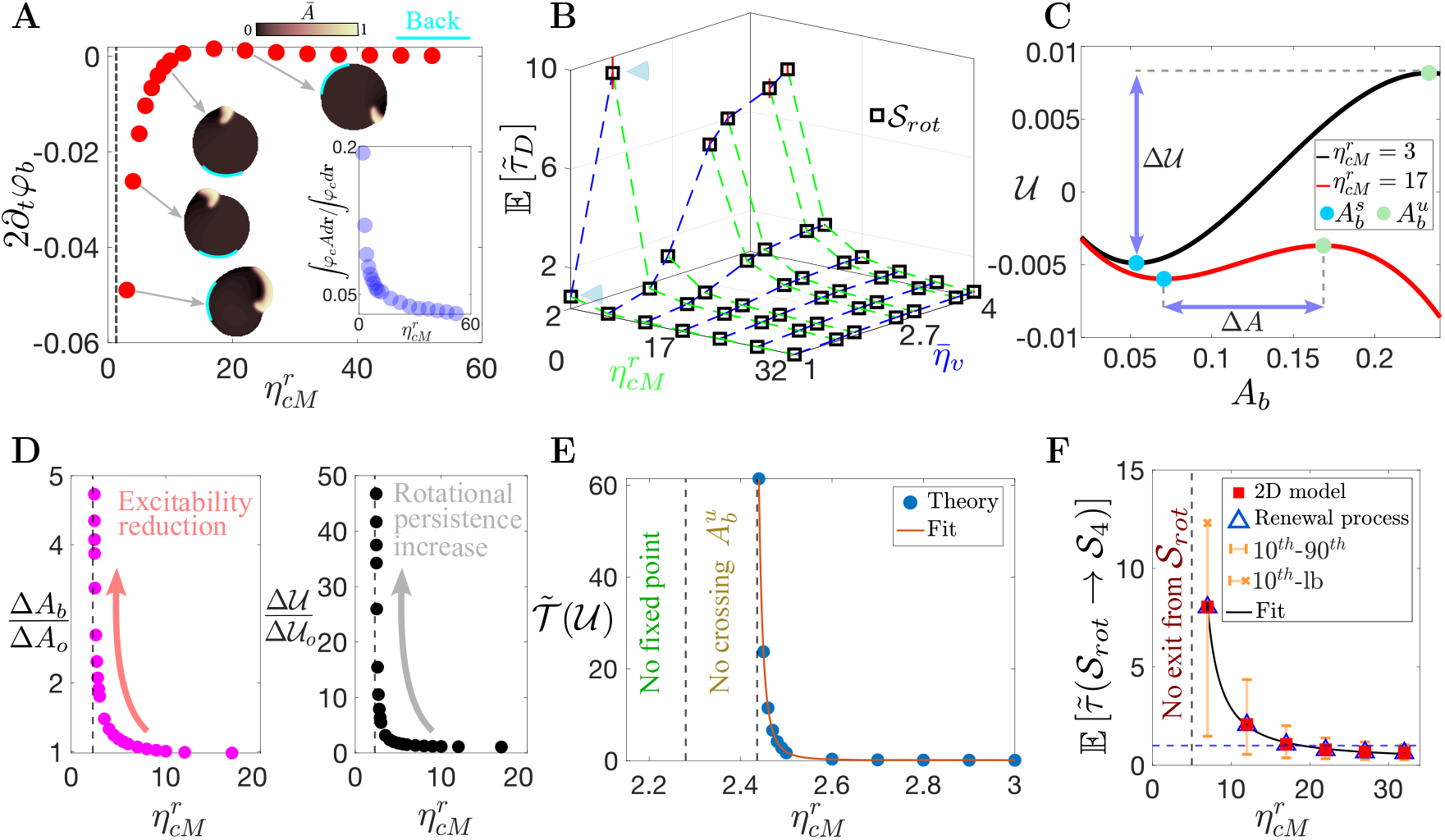
Back contraction regulates rotational persistence via mechanochemical feedback. (A) Confinement dependence of the mechanical feedback term ∂_*t*_*φ*_*b*_, calculated at the back of the cell in noiseless conditions. The insets show some of the spatial patterns of the normalized activator *Ā*, and a plot illustrating protrusion enlargement as confinement is weakened. (B) Phase diagram in 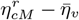 space showing the mean dwell time in the rotating state. For display purposes we have introduced a logarithmic scale, 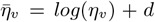 with *d* = 4.6. The blue triangles indicate measurable mean dwell times at 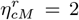 (exit from 𝒮_*rot*_ is possible). (C) Free energy *𝒰* in the quasi-steady approximation for two different confinements. The arrows represent the potential barrier Δ*𝒰* and the potential width Δ*A*. The circles are the relevant minima and maxima of 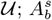 and 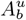, respectively. (D) Variations of the potential width (left panel) and of the potential barrier (right panel) as a function of the confinement. (E) Growth of the mean escape time in the quasi-steady approximation as confinement is weakened. The algebraic fit is 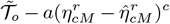 with fitting parameters *a* = 0.001 and *c* = 2.93. At 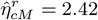 the crossing at 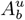 ceases to exist, and 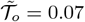 is the normalized mean escape time at at a baseline confinement 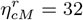. (F) Divergence of 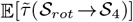 in the 2D model, close to the critical confinement 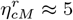. The algebraic fit is 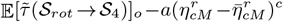 with fitting parameters *a* = 20.93 and *c* = 1.25. At 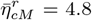, no exits from 𝒮_*rot*_ are observed after 100*T*. The baseline 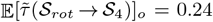 is measured at 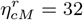. The orange whiskers correspond to the 10^*th*^ and 90^*th*^ percentiles of the first-passage time distributions. The orange (x) symbols indicates the lower bound (maximum first passage time observed) when a 90^*th*^ cannot be estimated (Supplemental Material [38]).

For this back contraction to enhance coherent rotations, it must modulate the activator-inhibitor dynamics. A back contraction of sufficiently large magnitude affects the local activator-inhibitor {*A*_*b*_, *R*_*b*_} excitable dynamics through the coupling with the phase field dynamics (Eq. 1), and thus with cell mechanical deformations. Specifically, the local dynamics of the activator and the inhibitor is modified by the linear terms 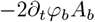 and 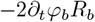 (Section IX in the Supplemental Material [38]), reducing the stochastic excitable dynamics at the back:

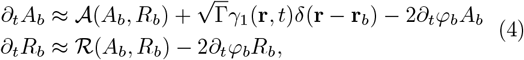

where **r**_*b*_ indicates the position of the back. The mechanical corrections modify the local excitable dynamics by increasing the distance between the fixed point (or resting state) 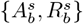 and the minimum of the cubic nullcline, raising the effective excitability threshold (Fig. S17 in the Supplemental Material [38]). This increase of the threshold limits the activation of new pulses, i.e., *P*(*𝒮*_4_ |*𝒮*_*rot*_) → 0, trapping the system in a persistent rotating state. Below a critical confinement, 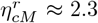, the fixed point at the back disappears and the zero-dimensional picture of the back of the cell being influenced by an isolated rotating pulse is not valid anymore. The slight confinement difference between the disappearance of the fixed point (at the cell back) and the emergence of the persistent coherent rotation 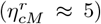 is likely due to finite-size nucleation effects in 2D or 3D geometry.

To formally address the excitability reduction and the transition to persistent coherent rotations, we adopt a quasi-steady approximation of the back activator-inhibitor dynamics: The slow inhibitor is frozen to its fixed point value 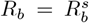, and the fast-scale dynamics for *A*_*b*_ drives the system. In this approximation, the back activator-inhibitor system can be written as an overdamped Langevin equation for *A*_*b*_ under a potential 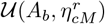 (Sextion X in the Supplemental Material [38] and Fig. 3C):

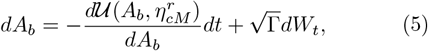

where *W*_*t*_ is a Wiener process [44] and the confinement dependence of the potential comes from the back contraction |∂_*t*_*φ*_*b*_|. As confinement weakens, both the potential well width 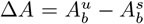 (with 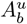 denoting the crossing of 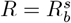 with the unstable branch of the nullcline; Fig. S1C) and the barrier height Δ*𝒰* increase (Fig. 3D). The growth of Δ*A* indicates that the distance from the fixed point to the excitability threshold becomes larger, i.e., the cell becomes less excitable. From a stochastic perspective, the increase of Δ*𝒰* maps to the overdamped Kramers escape problem [45]; a large potential barrier abolishes escapes at finite times.

Transitions out from the rotating state correspond to escapes from the potential well 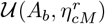, allowing nucleation of new protrusions; *𝒮*_*rot*_ → *𝒮*_4_. Therefore, to bridge the statistical analysis of discrete states with the zero-dimensional approach, we can quantify the stability of the rotating state by estimating the mean escape time from the potential well as a function of confinement, 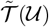. Introducing the corresponding Fokker-Planck equation for Eq. (5) and following standard analytical calculations [44], we obtain (Section X in the Supplemental Material [38]):

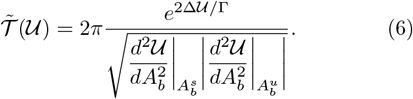

This mean escape time diverges algebraically as confinement decreases (Fig. 3E), indicating that the rotating state becomes increasingly stable. The singularity is marginally shifted from 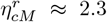, and is related to the extinction of the crossing at the unstable branch 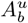. This mean escape time, valid in the zero-dimensional approach, serves as a mechanistic proxy for the empirical mean first passage time 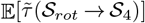, conditioned to the escape being possible. Consistently, 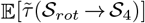 also shows an algebraic diverging behavior near the transition to the regime of permanent rotation (Fig. 3F).

### D. Experiments validate weak confinement regime predictions

Our theory provides a possible resolution to the apparent conflict among experimental observations of single MCF10A cell chirality in Matrigel: sometimes these cells rotate [7], and sometimes they do not [32]. Specifically, our modeling framework suggests that the breaking of chiral symmetry is stochastic and depends on the level of confinement. Therefore, an experiment with sufficient statistics across different confining conditions should reveal the probabilistic behavior of single-cell chiral motion. We test this prediction by seeding multiple MCF10A cells in Matrigel at different concentrations (30%-100%) and tracking their behavior by imaging F-actin (LifeAct-GFP) and the nucleus (H2B-mCherry) over a maximum of 16 h (see Sextion XI in the Supplemental Material [38]), prior to cell division (18-20 h).

In some cells, we observe distinctive F-actin configurations consistent with states *𝒮*_1_, *𝒮*_2_, *𝒮*_*rot*_, *𝒮*_5_ and *𝒮*_6_ (Fig. 4A, and Videos S8-S10 in the Supplemental Material [38]). The most frequently observed experimental state is *𝒮*_1_. A plausible reason for not observing *𝒮*_7_ and *𝒮*_8_ is that resolving three F-actin patches requires a cell size that is large relative to the patch size. While individual F-actin patches can be identified manually, systematic quantification of their dynamics, similar to Fig. 2, is challenging due to difficulties in robustly thresholding the images to distinguish states, as well as the limited observation windows imposed by cell division. We therefore adopt a simpler binary classification, labeling each experimental realization as either exhibiting cell rotation or not (see Section XI in the Supplemental Material [38]).

**FIG. 4.**
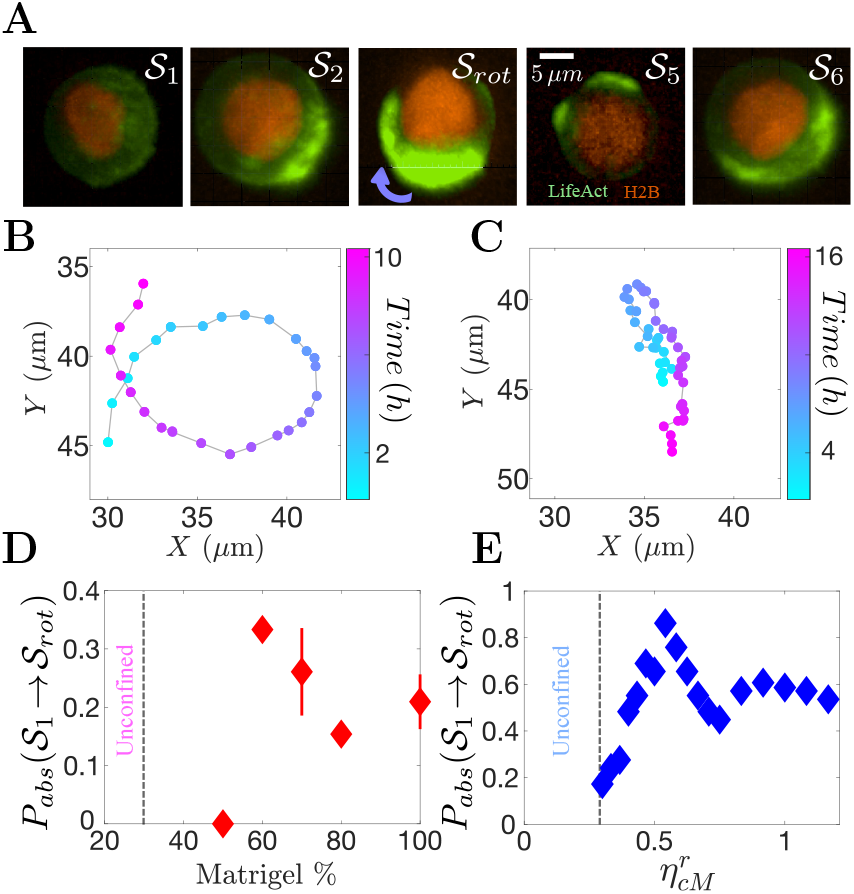
Chiral motion of MCF10A cells in Matrigel. (A) Temporal snapshots of F-actin distributions corresponding to distinct-states observed in 70% and 100% Matrigel. (B) Nucleus trajectory of the rotating cell, 𝒮_*rot*_, shown in (A). (C) Nucleus trajectory of a non-rotating cell in 100% Matrigel. (D-E) Absorption probability of the passage from a non-chiral to chiral state (𝒮_1_ → 𝒮_*rot*_) for different Matrigel concentrations in the experiments and for different 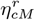 values within the weak confinement regime in the model, respectively. Bars in (D) indicate standard deviation.

Rotating cells exhibited two key features consistent with the weak confinement regime of our model 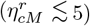. First, the nucleus, a proxy of the cell centroid, translated during the rotational motion (Fig. 4B), demonstrating that the entire cell moves instead of a protrusion orbiting around the surface of an almost static cell. Second, rotating cells displayed a distinct F-actin patch on their surface (Fig. 4A and Video S9), likely corresponding to the large protrusion that drives rotation in the simulations. In contrast, non-rotating cells showed no rotational motion over the entire window of observation, and in several cases cells drift across the ECM (Fig. 4C and Video S11). This translational motion likely arises from cells pushing and deforming the matrix while being still confined, a mechanism not captured by the static ECM in our model.

The analysis of more than 10 cells at each Matrigel concentration revealed that rotation does not emerge deterministically but rather occurs with a probability that depends on the concentration (Fig. 4D). In our framework, this corresponds to the absorption probability of the constructive path *P*_*abs*_(*𝒮*_1_ → *𝒮*_*rot*_): the probability of transitioning from the testing state to the rotating state over a finite observation window. We note that at 30% Matrigel cells sink to the bottom, pointing to a minimal confinement to have chances of observing rotations. While the physical mechanisms differ, this breakdown of confinement parallels the unconfined limit in the model 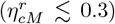. To compare the stochastic nature of the emergence of rotations between experiments and theory, we compute *P*_*abs*_(*𝒮*_1_ → *𝒮*_*rot*_) in the model across weak confinement values 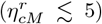. The theory predicts that *P*_*abs*_(*𝒮*_1_ → *𝒮*_*rot*_) = 1 if the observation window accounts for many rotation periods *T* (see the exemplary trajectories in Fig. S7 for 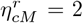). However, in the experiment, the observation window is restricted to approximately 2*T*, with *T* = 9.7 ± 1.2 h. This biological constraint dramatically affects *P*_*abs*_(*𝒮*_1_ →*𝒮*_*rot*_), as cells may not transition to a chiral state before division. Using an observation window of 2*T* to compute *P*_*abs*_(*𝒮*_1_ →*𝒮*_*rot*_), across different weak confinements, yields qualitative agreement with experiments (Fig. 4E). The experimental decay in the probability of observing rotations at the lowest Matrigel concentration where cells are still confined (50%) possibly reflects insufficient confinement, which slows protrusion retraction, and thus the birth of rotational waves.

## III. DISCUSSION

### A. Spontaneous chiral symmetry breaking

One of the key findings of our work is that single-cell chirality arises spontaneously from an intrinsically nonchiral model. The homogeneous confinement does not impose chirality but rather establishes geometric and mechanical conditions (e.g. protrusion size) where spontaneous symmetry breaking becomes possible. While previous experimental works demonstrated cell chirality in single and multiple cells [7, 33, 46], they addressed this emergent phenomenon as a consequence of an explicit symmetry breaking either inside or outside the cell. There is theoretical work approaching spontaneous chiral symmetry breaking in polarized epithelial cells, but only from a multicellular perspective [47]. Our results suggest that there is no necessity of pre-existing chirality at the molecular level for the rotations to emerge, but that nonlinear coupling between excitable dynamics, confinement and cell deformations at the macroscopic level is sufficient. This positions single-cell chirality within a general physics framework of symmetry-breaking in out-of-equilibrium systems [48, 49].

### B. Physical interpretation of cell confinement

In our model, confinement is prescribed by a local isotropic force acting at the cell-ECM interface. This confining effect can be understood through a simplified one-dimensional version of Eq. (2), in which we find that, at first order approximation, the local normal velocity at the cell interface obeys 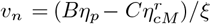, where *B* and *C* are positive constants (Section XII in the Supplemental Material [38]). This simple relationship aligns with the findings presented in the text: Strong confinement can result in negative values for this normal velocity and thus suppresses the growth of local deformations. In other words, large values of 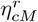 impose fast protrusion retraction that overcomes protrusion extension. The analogy between the degree of confinement and the protrusion retraction timescale suggests that the confinement modeled in this work can be interpreted as reflecting ECM stiffness [50], where the strong (weak) confinement regime could represent a stiff (soft) environment. This interpretation can be directly connected to the results in Figs. 4D and 4E, where variations in Matrigel concentration, with reduced concentrations corresponding to decreased stiffness [51], are mapped to variations in 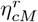. Reductions in Matrigel concentration have also been linked to a decrease in ECM ligand density, which weakens cell-ECM adhesive interactions [52]. This lowers resistance to cell membrane deformation, effectively reducing cell confinement and suggesting that cell-ECM adhesion provides an additional mechanism for controlling it, which could be tested in an extension of our model.

The notion of confinement may also be extended beyond cell-ECM interactions. For example, a cell surrounded by neighboring cells could be sufficient to establish the necessary confinement for cells to display chiral symmetry breaking. In fact, during *Drosophila* oogenesis, there is a stage where nurse cells—surrounded by either other nurse cells, the oocyte, or follicle cells—dump cytoplasm into the oocyte while sustaining rotating cortical myosin waves on their surfaces [53].

In our experimental study, we considered epithelial MCF10A cells because they do not remodel the ECM as much as other cells. However, some cells have the ability to significantly remodel the ECM, enabling them to control their own confinement. This is the case for MDA-MB-231 cells seeded in collagen type I, which release matrix metalloproteinases to degrade their surroundings and make room for cells to come after cell division [9]. From the modeling perspective of a single cell, the degradation could be understood as a reduction in effective confinement due to extra free space (Section XII in the Supplemental Material [38]). Beyond such active remodeling, the effective confinement can also change over time through ECM stress relaxation [54]. Future studies could include matrix degradation, stress relaxation dynamics, and exploring how cell-ECM interactions are modified after ECM remodeling [55, 56].

### C. Physical interpretation of volume/area conservation and nonlocal mechanics

Our phase field model includes a size-restoring force, conserving the volume (in 3D) or the area (in 2D). Although the motivation of this force relies on the argument of the incompressibility of the cytoplasm, volume control in real cells involves multiple processes such as osmotic pressure, ion pumps, and tension generated by the membrane and the actomyosin cortex [57–59]. The nonlocal character of **F**_*v*_, where local deformations (protrusions) induce immediate long-range contractions elsewhere, mirrors the stress transmission effect of the actomyosin cortex [60]. Thus, the simple nonlocal force— widely used in phase field modeling—incorporates not only the phenomenology of volume conservation, but also other processes involving cell contraction. As we showed in Fig.3B, small *η*_*v*_ values do not sustain rotations even for weak confinements. This could be connected to experimental observations where inhibition of actomyosin network regulators (Myosin II, myosin light-chain kinase and Rho-associated kinase) led to loss of rotational coherence [7]. In addition, the relevance of cell contractility for persistent rotational motion (Fig. 3B) may be related to experimental findings in endothelial cell doublets seeded on micropatterns, where the cell with higher contractility imposes the chirality of the doublet [61].

### D. Semi-Markov renewal process for excitable cellular dynamics

The semi-Markovian approach we developed for the stochastic regime represents a broadly applicable technique for studying excitable systems with complex spatiotemporal dynamics [62]. This statistical framework is particularly useful for systems where the states of the system have characteristic timescales (refractory periods, slow wave dynamics) and experience noise-driven transitions. The interplay between deterministic and stochastic elements naturally map into semi-Markov dynamics. Our statistical approach could be extended to other biological systems—governed by excitable responses and membrane deformations—exhibiting stochastic switching between dynamical behaviors. For example, F-actin wave dynamics in Dictyostelium discoideum cells on substrates display birth-death dynamics [63], offering a well-known experimental system to extend the coarse-grained technique. We highlight that our method is data-driven so given the enough spatiotemporal resolution it should be easily applied to experimental data.

### E. Robustness of chiral motion

The model introduced here requires fixing 19 parameters (see Table S1), which control cell mechanics and wave dynamics within the cell. It is thus important to assess how the parameter choices influence the emergence and maintenance of chiral motion in single-cells.

The parameters governing the local activator-inhibitor dynamics establish either an excitable or an oscillatory regime, depending at which point the linear and cubic nullcline intersect (fixed point). Extensive experimental evidence shows that cortical actin waves in migrating and mechanically constrained cells predominantly exhibit excitable properties, including threshold responses and refractory periods [22–24, 26, 64]. Relevant to our case, excitable waves have also been reported under geometric or adhesive confinement [65, 66], suggesting that excitable dynamics is the prevailing behavior of actin waves across diverse biological contexts. For this reason, we developed our theoretical framework considering the excitable regime of Eq. (1), but additional simulations in the oscillatory regime also show the emergence and maintenance of rotating states (see Fig. S18 and Section XIII in the Supplemental Material [38]).

As illustrated in Fig. S17 [38], the level of excitability depends on the distance 𝒟 from the fixed point to the minimum of the cubic nullcline. Therefore, to robustly obtain coherent rotations for which surpassing the excitability threshold is mandatory, the parameters controlling the local dynamics in Eq. (1) must satisfy the necessary condition of a sufficiently small 𝒟 (specifically, 𝒟≲ 0.32), enabling activator nucleations (Fig. S19 in the Supplemental Material [38]).

The possibility of nucleating an activator wave, either in the excitable or in the oscillatory regime, is not only dictated by the local nonlinear dynamics of Eq. (1), but also depends on the length scales of the spatially extended system, which are set by the diffusion coefficients *D*_*A*_ and *D*_*R*_. Larger activator length scales (larger *D*_*A*_), relative to the cell size, make the nucleation of activator waves more difficult, which in turn will decrease the probability of observing rotating waves where multiple nucleations are necessary. Moreover, fluctuations mediate the emergence of rotational motion, and their appearance also depend on the activator length scale being sufficiently small. Therefore, given a cell size—here motivated by MCF10A cells (diameter ~15 *µ*m)—and the parameters in Table S1, the ratio *D*_*A*_*/D*_*R*_ must be smaller than 0.7 to robustly observe cell rotations (Fig. S20 in the Supplemental Material [38]).

The effects of the parameters directly influencing cell mechanics are easier to grasp compared to the parameters involved in the activator-inhibitor dynamics. For example, if the friction force dominates over all others (large *ξ*) rotational coherence is impaired due to the inability of the cell to produce motion (Fig. S21 in the Supplemental Material [38]). Likewise, decreasing the protrusive force relative to the other forces (small *η*_*p*_) leads to the loss of rotations due to the inability of the system to generate a persistently polarized protrusion [20, 35].

While our model qualitatively captures the chiral motion of MCF10A cells within Matrigel (Figs. 4D and 4E), systematic optimization of the model parameters discussed in this subsection could potentially improve the quantitative agreement.

## IV. CONCLUSIONS

In summary, we demonstrate that confinement by the ECM shapes single-cell chiral dynamics through a rich interplay between mechanics, coarse-grained chemical signaling and stochasticity. Our theoretical framework identifies three dynamical regimes controlled by confinement strength. In the intermediate, or stochastic, regime, a data-driven analysis establishes that a semi-Markov renewal process governs the emergence and maintenance of the rotational motion. Under weak confinement, a mechanochemical feedback stabilizes persistent chiral states. This feedback operates in the form of a nonlocal contraction, activated in the presence of the rotating protrusion, lowering the cortical excitability. Experiments on MCF10A cells in Matrigel validate theoretical predictions for the weak confinement regime. Our work opens avenues for understanding and potentially controlling single-cell dynamics under confined conditions, and suggests immediate follow-ups. In particular, promising directions include new experiments that test the theoretically predicted transition from persistent to stochastic behavior, and investigations into the biological role of single-cell rotation and its relationship to multicellular rotating structures formed after cell division.

The authors thank Mahesh Kumar Mulimani for his comments on this manuscript. This work was supported by NSF MCB 2426002 and NSF PHY 2310496 to W.-J.R., and by a Prebys Foundation Research Heroes grant to S.I.F. We would like to thank the UC San Diego School of Medicine Microscopy Core, which is supported by the National Institute of Neurological Disorders and Stroke grant P30NS047101. S.E.-A. acknowledges the financial support of ANID by Beca Chile 74230063.

The data that support the findings of this article are openly available [67].

## Supplemental Material

## I. Phase field model

We model confined single-cell dynamics using a phase field approach. Let us consider a scalar field *φ*_*c*_(**r**, *t*) where **r** = (*x, y, z*), which provides a smooth and continuous representation of the cell position. Specifically, *φ*_*c*_ = 1 inside the cell and *φ*_*c*_ = 0 outside the cell, with the two states connected through an interface of width *ϵ* (Fig. S1A). The spatiotemporal evolution of the cell interface is governed by an advection equation, 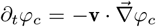, where **v** is the velocity of the cell membrane in the outward normal direction, 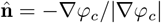. We assume that cell motion occurs in a highly viscous extracellular matrix (ECM); thus, the friction at the cell-ECM interface can be modeled as a simple linear drag, **F**_*f*_ = −*ξφ*_*M*_ **v**, where *ξ* is the friction coefficient and *φ*_*M*_ is the position of the ECM (see below). Then, a force balance at the cell membrane determines the velocity (Fig. S1B):

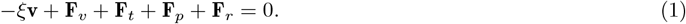

The second term in Eq. (1) represents an isotropic size-restoring force acting in the normal direction, which enforces a prescribed cell size 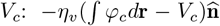, where *η*_*v*_ is a parameter controlling the strength of this force. The third term models the membrane tension of the cell. In equilibrium, this tension is associated with a Ginzburg-Landau type of energy 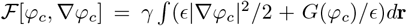, and the corresponding force can be expressed as 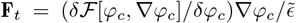 [1], where 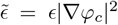 and *γ* is the surface tension coefficient. The function 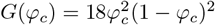 is a double well potential with minima at *φ*_*c*_ = 0 and *φ*_*c*_ = 1.

The fourth term is a non-equilibrium force in the outward normal direction that drives cell motion through protrusions, governed by activator-inhibitor (*A*-*R*) dynamics: 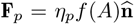. The parameter *η*_*p*_ is chosen such that, in the absence of the ECM, the cell exhibits a persistent unidirectional motion, which can be interpreted as migration on a substrate [2, 3]. The activator *A* represents a mean-field description of a signaling protein that models cell polarization by promoting actin polymerization. Following previous studies, we adopt the simple linear case *f*(*A*) = *A* [1, 4]. However, we have verified that the more complex Hill-type function 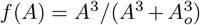, used in Ref. [2], can also generate single-cell rotation. The activator-inhibitor dynamics is restricted to the cell interior through a function Ω_*c*_(*φ*_*c*_), which implicitly imposes non-flux boundary conditions, and is described by the coupled reaction-diffusion equation:

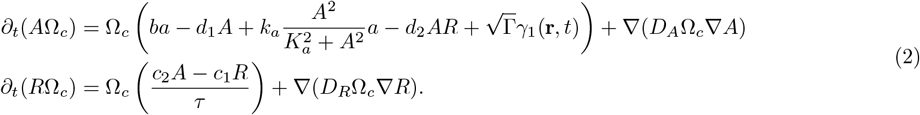

This equation is equivalent to Eq. (1) of the main text with

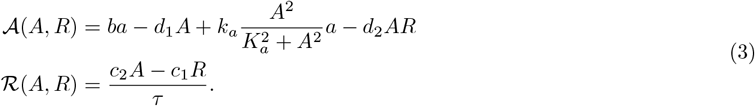

The reaction terms have a similar functional form to those used in previous studies [5], and Eq. (2) has been used in previous models of cell plasticity in both two and three dimensions [2, 3]. The term *γ*_1_(**r**, *t*) denotes delta-correlated Gaussian noise in space and time, with ⟨*γ*_1_(**r**, *t*) ⟩ = 0 and ⟨*γ*_1_(**r**, *t*) ⟩ ⟨*γ*_1_(**r**^*′*^, *t*^*′*^)⟩ = *δ*(**r** − **r**^*′*^)*δ*(*t* − *t*^*′*^), modeling the inherent fluctuations of microscopic protein dynamics. We have verified that including noise in both the activator and inhibitor equations does not affect our results in a qualitative way. The parameters in Eq. (2) are chosen such that the activator-inhibitor system exhibits excitable dynamics (Table S1 and Fig. S1C) and, as discussed above, induces unidirectional migration in the absence of confinement. Up to this point, the modeling framework has considered a three-dimensional cell (Fig. S1A). In this case, Ω_*c*_(*φ*_*c*_) = 2*G*(*φ*_*c*_)/*ϵ*, restricting the activator-inhibitor evolution near the cell surface to minimize numerical costs. Nevertheless, we also employ the same approach to study cell dynamics in a two-dimensional representation, where we define Ω_*c*_(*φ*_*c*_) = *φ*_*c*_(*x, y, t*). Choosing the later for the 3D case does not affect qualitatively our results. Importantly, irrespective of the smooth function Ω_*c*_ used to indicate the cell position in space, the true free boundary problem for the activator-inhibitor system in a moving domain is recovered in the sharp interface limit (*ϵ* → 0) [6].

The last term in Eq. (1) accounts for the confining effect of the ECM. To model this interaction, we introduce a second phase field, *φ*_*M*_ (**r**), which specifies the position of ECM (Fig. S1A). The ECM is static, and its geometry is generated by carving a spherical (circular) hole in a cube (square) of size *L*^3^ (*L*^2^), and subsequently allowing the interface to relax diffusely until a prescribed volume *V*_*M*_ is reached (see Section II). The force exerted by the ECM is given by 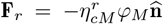, where 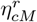 is the parameter controlling the confinement strength. We have further verified that modeling confinement as a repulsive potential interaction between the cell and the ECM, using 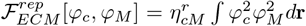, does not alter our conclusions. Finally, substituting the velocity from Eq. (1) into the advection equation for *φ*_*c*_ yields:

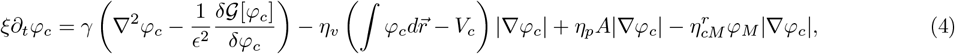

which is valid only when the cell is in contact with the ECM. Note that Eq. (4) implicitly incorporates cell contraction. For example, immediately after a protrusion locally extends the interface of a cell at equilibrium, the cell size increases: 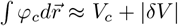. This results in a contraction away from the protrusion with a strength proportional to the size increment |*δV*|. We have verified that explicit incorporation of local cell contraction, for example driven by myosin activity [7], as implemented in previous studies [4, 8], does not qualitatively alter our results. All parameters in Eq. (4) are summarized in Table S1.

**Figure S1:**
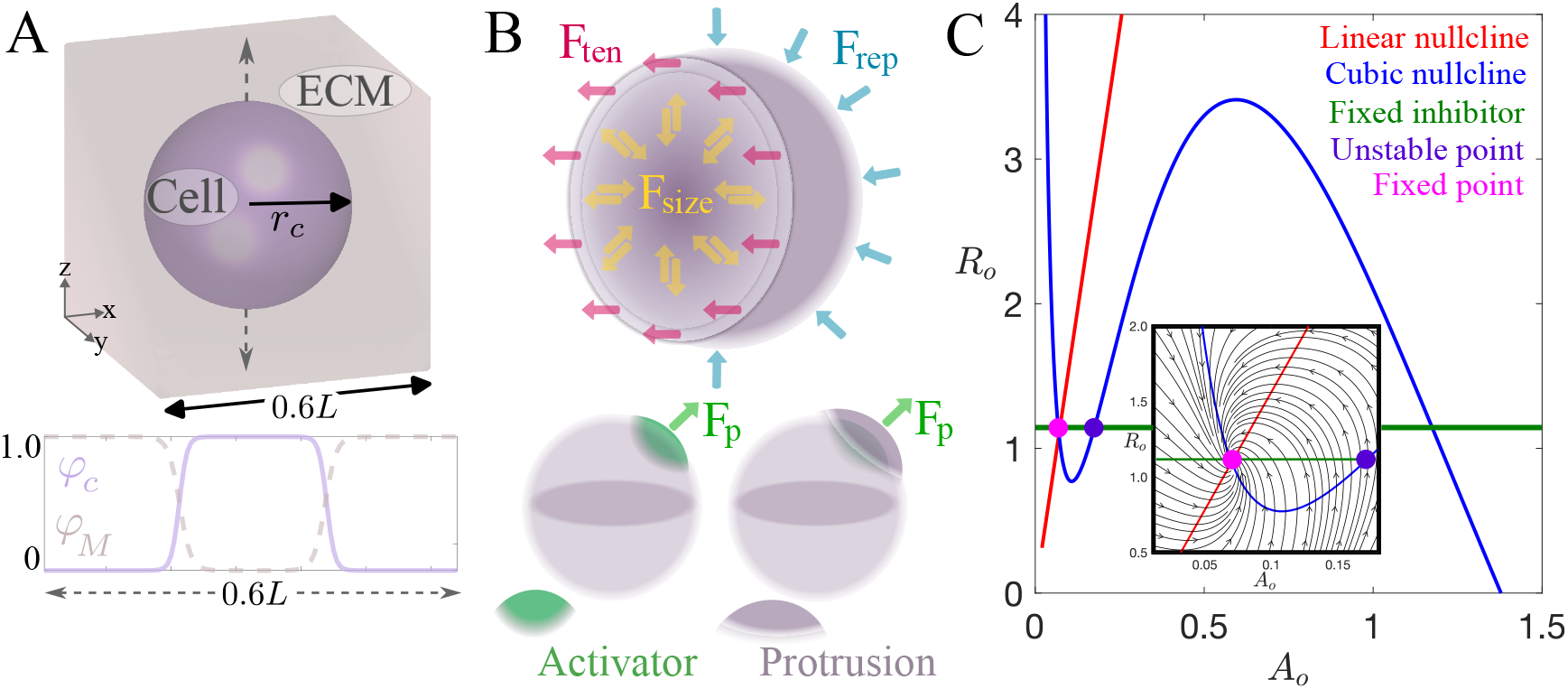
Model set-up. (A) Top panel: Initial three dimensional phase fields. For the cell, the phase field is shown at *φ*_*c*_ = 1/2, while the ECM phase field is shown for *φ*_*M*_ *>* 1/2. Bottom panel: One-dimensional cuts of *φ*_*c*_ and *φ*_*M*_ along the dashed line in the top panel. *r*_*c*_ denotes the initial radius of the spherical cell. (B) Schematic representation of the forces acting on the cell surface, excluding ECM-cell friction. (C) Phase space of the zero-dimensional version of the activator-inhibitor dynamics in Eq. (2), given by {*A*_*o*_, *R*_*o*_}, in the excitable regime. The inset shows the structure of the phase flows around the fixed point (attractor). The unstable point, crossing between the fixed inhibitor and the cubic nullcline, is introduced further in this Supplemental Material.

## II. Numerical methods

The initial condition for the ECM, in both 2D and 3D simulations, is given by the steady state of an equation similar to Eq. (4), but dimensionless:

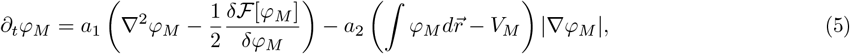

where *a*_1_ = 2/150 and *a*_2_ = 100/150 sets the strength of the diffusive process and the strength of prescribing the ECM size, respectively. This reaction-diffusion equation is discretized using finite differences with anisotropic stencils in the {*x, y, z*} directions in 3D ({*x, y*} directions in 2D) with non-flux boundary conditions and integrated in time with a forward Euler scheme. The integral is discretized using the trapezoidal method. The initial condition for the cell, *φ*_*c*_, is designed to emulate the experimental scenario of a cell seeded into an ECM. Particularly, the phase field *φ*_*c*_ is initiated in the cavity of *φ*_*M*_ as 1 − *φ*_*M*_, and is evolved for a short period with *η*_*v*_ = *η*_*p*_ = 0 and 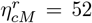 until reaching an equilibrium size determined by the balance between curvature and confinement (Fig. S1A). This equilibrium size is then used as the prescribed cell size *V*_*c*_.

The three-dimensional phase field simulations were implemented in Matlab using GPU acceleration. The equation for *φ*_*c*_ is solved via a pseudospectral integrating factor method with a RK4 time-stepping [9]. In this numerical scheme, the linear part is directly approximated in Fourier space, while the nonlinear terms are discretized using fast Fourier transforms (FFT) and transformed back to real space via inverse FFT. We use periodic boundary conditions for the phase field equations. Once *φ*_*c*_ is obtained at times *t* and *t*+Δ*t*, the activator and inhibitor equations are solved using finite differences with anisotropic stencils (two neighbors per spatial direction). For example, the weighted diffusive term in the x-direction (for fixed *y* and *z*) is discretized as

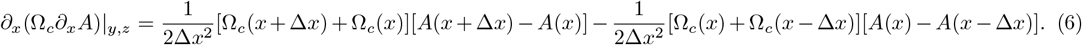

The corresponding terms in *y* and *z* directions of *A* as well as the diffusive terms for *R* are computed analogously. The weighting by the phase field automatically introduces non-flux boundary conditions. The activator and inhibitor are marched in time only inside the cell, 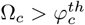, using a forward Euler scheme. For example, the activator is updated as (similarly for the inhibitor):

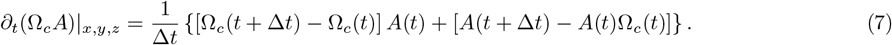

The noise terms *γ*_1_(**r**, *t*) and *γ*_2_(**r**, *t*) are generated using a Box-Mueller transform applied to two uniform random samples, and their discretization follows the Ito representation.

The two-dimensional simulations are performed either using WebGL, with custom Java and GLSL codes based on the *Abubu*.*js* library [10] (see Section III), or using Matlab. Numerically, the main difference from the 3D simulations is that the entire discretization of the phase field equation is carried out in real space using finite differences. Additional details can be found in the repository for this paper [11]. The numerical parameters are summarized in Table S1.

**Table S1:**
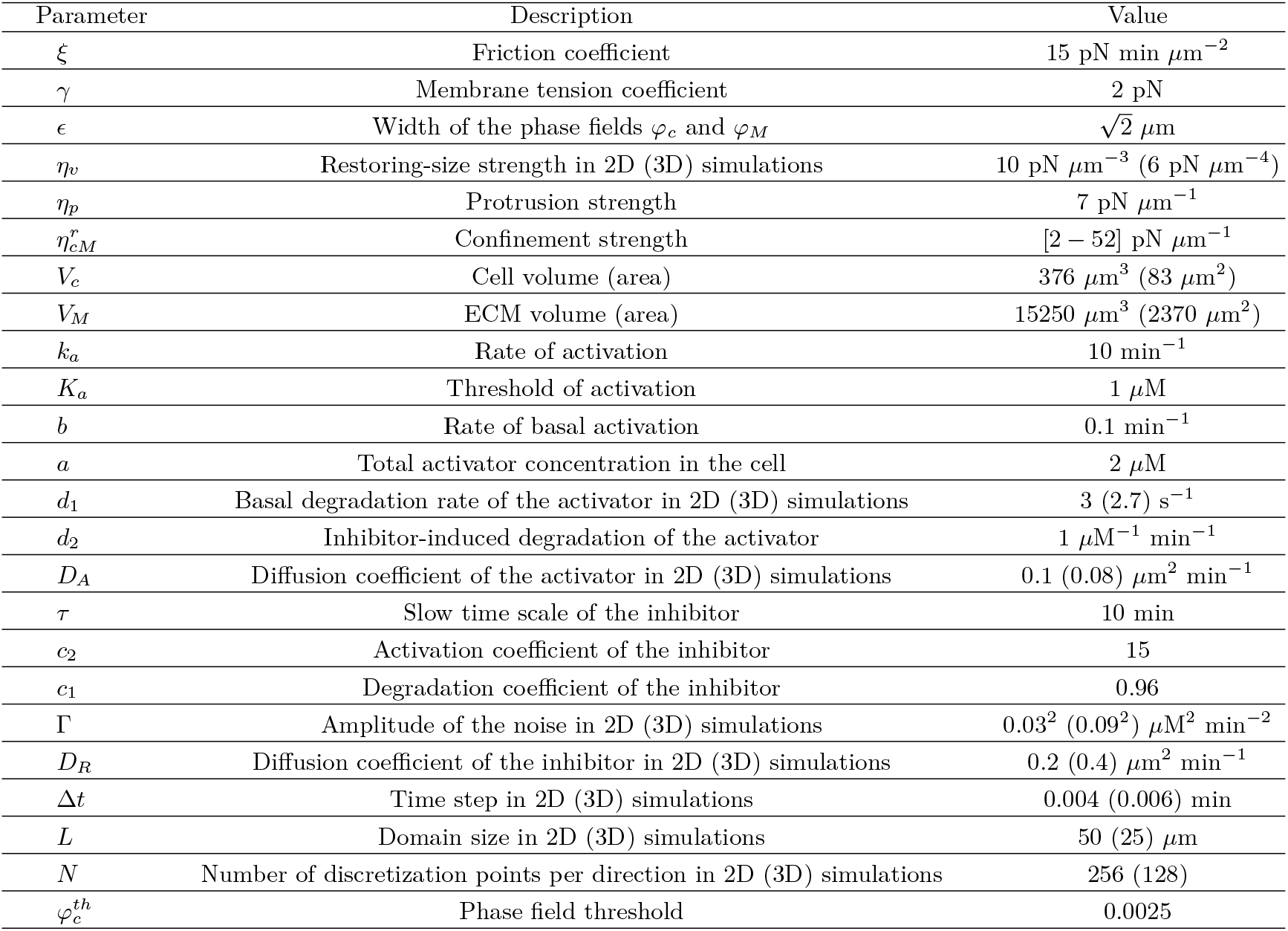
Model and numerical parameters used for all the 2D and 3D computational simulations in this study. Note that Γ and *η*_*v*_ are fixed except in Figs. S11 and S16, respectively.

## III. Numerical implementation of the 2D phase field model in WebGL

We have developed a GPU-accelerated WebGL platform to solve the two-dimensional forms of Eqs (2) and (4) [11], using a specialized library for integrating partial differential equations via fragment shaders [10]. The simulations run directly in a web browser (e.g., Google Chrome, Safari, Firefox), enabling real-time interaction with the dynamics. For example, a click on top of the cell can perturb its activator field *A*, allowing the manual generation of protrusions. Additionally, the code is capable of storing data—simply by taking snapshots of the canvas on the webpage—while bypassing the usual GPU-CPU memory-transfer bottleneck [12]. We have leveraged this storage approach to efficiently explore multiple parameters in parallel (see [11] for details).

## IV. Supplemental figures for Fig. 1: Figs. S2, S3, S4 and S5

The activator-inhibitor dynamics in the weak and intermediate regimes that lead to rotational motion involve the break-up of target waves along the cell surface. This breakage can be triggered either by stochastic fluctuations (Fig. S2), or by heterogeneities in the refractoriness of the excitable system (Fig. S3), which are dictated by the presence of multiple waves interacting along the cell surface.

The rotational motion in the weak confinement regime 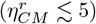 is characterized by a frustrated “figure of eight” activator pattern (Fig. S4), and by a rotational period *T*. This frustration arises because the characteristic wavelength of the excitable waves—imposed by the diffusion coefficients *D*_*A*_ and *D*_*R*_—is relatively large compared with the finite size of the cell. Another signature of the coherent motion is the increase in the amplitude of the centroid trajectory, which reflects protrusion enlargement as confinement weakens (Fig. S5).

**Figure S2:**
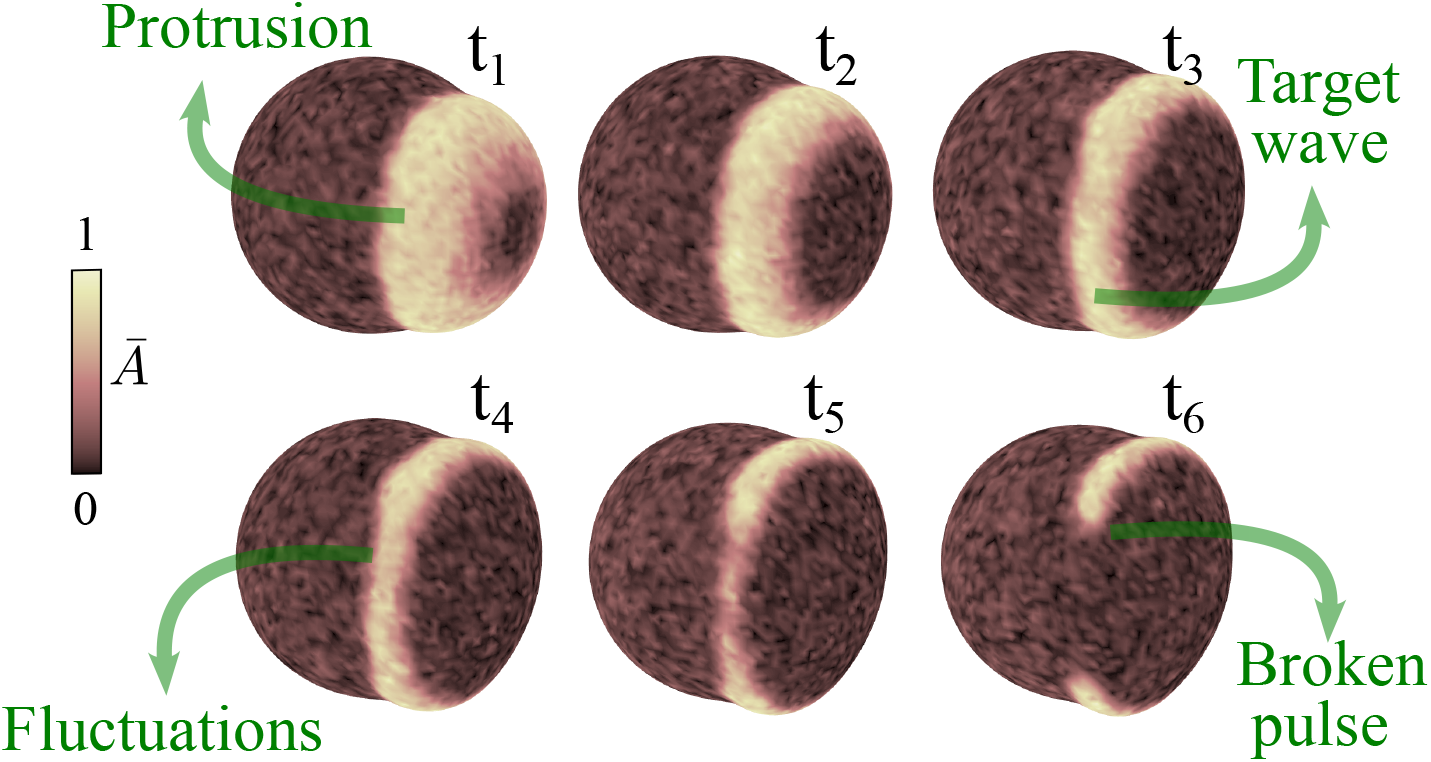
Temporal snapshots (*t*_1_ *< t*_2_ *< t*_3_ *< t*_4_ *< t*_5_ *< t*_6_) of the normalized activator field *Ā* = *A*/ max (*A*), illustrating the break-up of a target wave induced by stochastic fluctuations in the intermediate regime 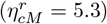.

## V. Coarse-graining into 𝒮-space

To coarse-grain the 2D phase-field data into the discrete space *𝒮*, we begin by processing the numerical results in MATLAB, using output from 2D simulations of the phase-field and activator–inhibitor dynamics using WebGL. We use the activator field *A* to detect pulses (case 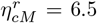 in Fig. S6A). First, the raw RGB image of the activator is smoothed using a Gaussian blur with a 9 × 9 kernel to homogenize the noisy background. Next, a binary image (mask) is created to indicate where the activator exceeds 90% of its maximum value. To avoid holes in the mask, a dilation process with a 9 × 9 kernel is applied. Finally, we use the function *regionprops()* of Matlab to identify connected regions in the binary image and extract their centroids. The number of centroids serves as a proxy for the number of pulses.

Errors in the detection algorithm are expected due to noisy transient pulses that may become frequent in the large dataset under analysis. These transients exhibit a high value of *A* and are large enough to produce a measurable centroid, but they persist only for a short time. Figure S6B shows a raw trajectory of the number of pulses, where many of such short episodes are detected. It is important to remove them because they bias the computation of transition probabilities and first passage times. To address this, we implement a custom code to filter out spurious events defined as having a duration shorter than *T* /45. Briefly, the code detects segments of the trajectory, e.g., 11110001111, and replaces the sandwiched zeros with ones if their duration is smaller than *T* /45. Notice that events containing four pulses are always removed because they last less than the cutoff. The origin of this configuration is the fragmentation of two protrusions (Video S12), and their short duration is governed by the size-restoring force preventing the presence of more than three pulses. Figure S6C illustrates the temporal evolution of the number of pulses after applying the cutoff *T* /45.

**Figure S3:**
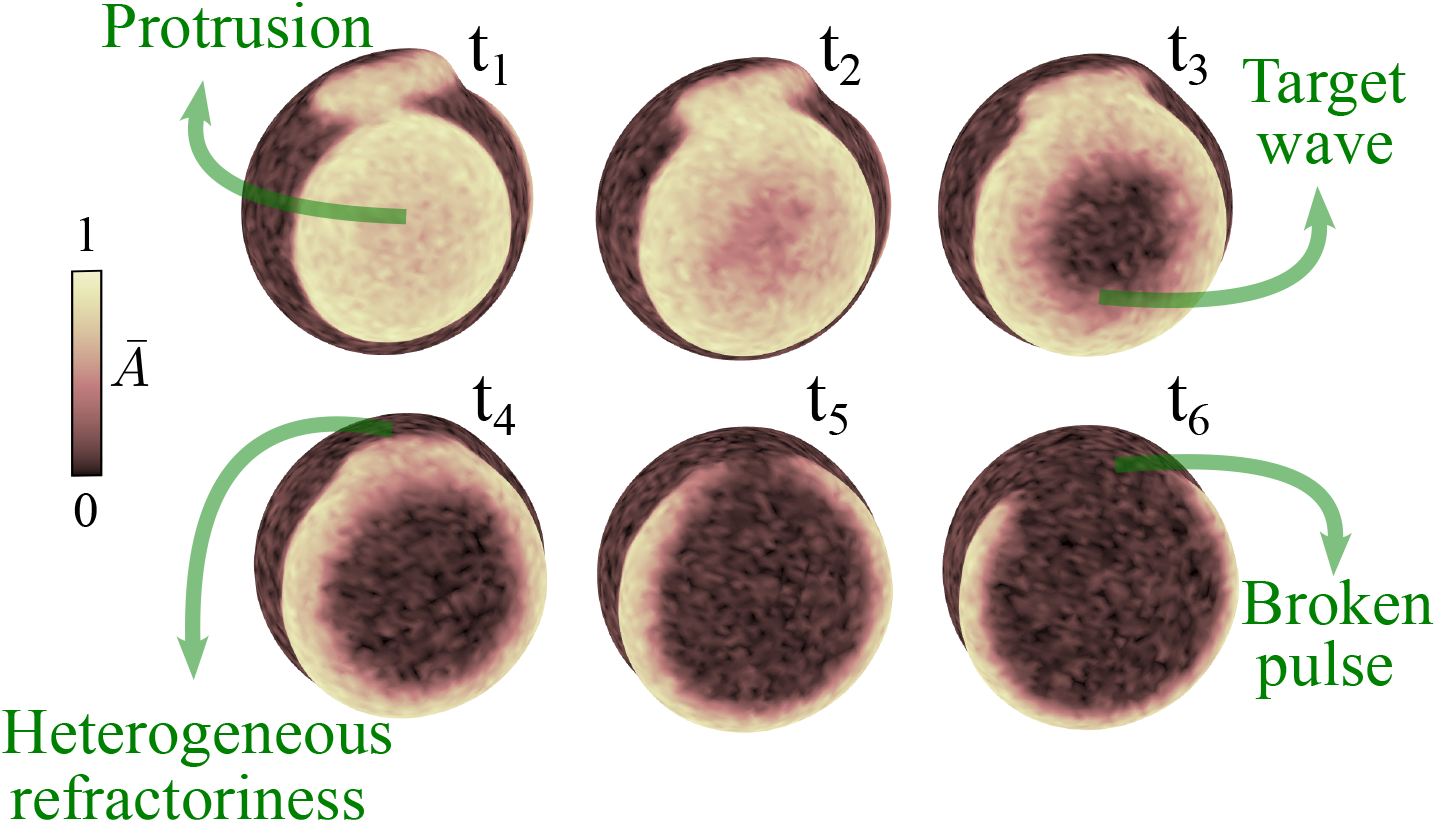
Temporal snapshots (*t*_1_ *< t*_2_ *< t*_3_ *< t*_4_ *< t*_5_ *< t*_6_) of the normalized activator field *Ā* = *A*/ max (*A*), illustrating the break-up of a target wave induced by heterogeneities in refractoriness in the intermediate regime 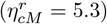.

**Figure S4:**
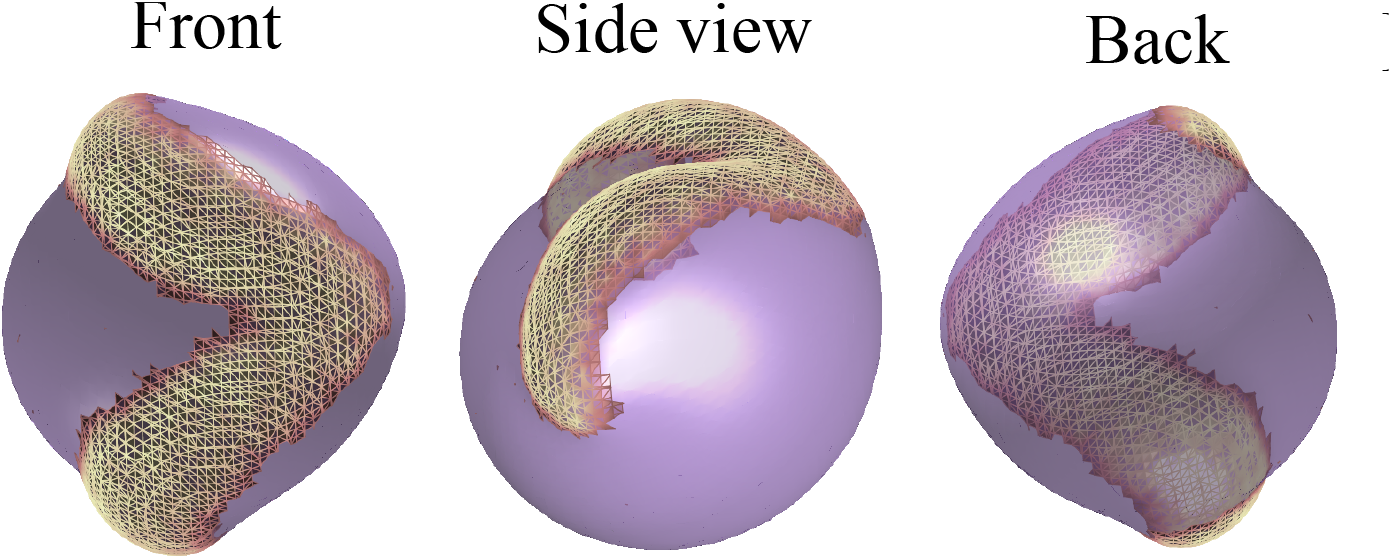
Coherent rotational wave. Different three-dimensional views of the activator spatial structure in the rotating state at 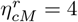. Only the portion *A >* 0.8 of the activator field is shown.

**Figure S5:**
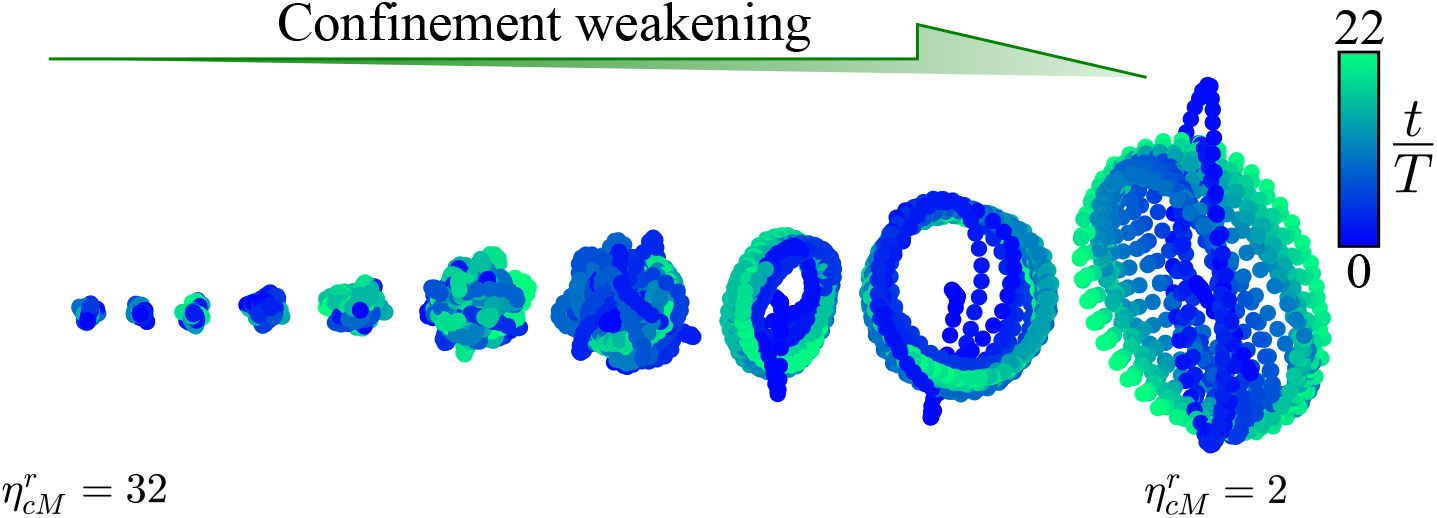
Three dimensional trajectories of the centroid of the cell for the range 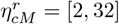 over 22*T*. The trajectories start at the initial condition.

Detecting the number of pulses alone is not sufficient to distinguish between the possible discrete states of the system, as discussed in the main text. Particularly, both a single chiral pulse and a non-chiral pulse yield the same pulse count of one, but their difference is fundamental. To tackle this, we include in our data analysis a routine to determine whether or not pulses rotate. For this, we compute the angle *θ* of the orbit of each activator pulse with respect to the cell center (rightmost panel; Figure S6A). Next, this angle is unwrapped and a linear model is fitted over the duration of the single pulse visit. For each visit, we calculate the coefficient

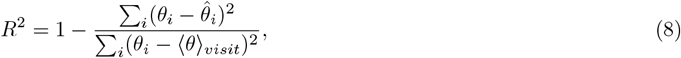

where 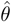 are the angle values predicted by the linear model, and ⟨*θ*⟩_*visit*_ is the average angle within the visit. If 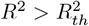, the single pulse event is classified as a rotating pulse. Short duration events may produce false positives by flagging non-chiral pulses as chiral pulses. We correct these cases by considering the quasi-deterministic transitions on the discrete system. For example, the transition out of the resting state typically proceeds through the creation of a non-chiral pulse, which then divides into two chiral pulses (Video S7). If that one-pulse state, which has a short duration (*<* 0.25*T*), is flagged as a chiral pulse, we reclassify it as non-chiral.

The same procedure is applied to differentiate between chiral and non-chiral structures when the pulse count is equal to two, by defining two angles. However, when the pulse count is equal to three, the differentiation by angle is more cumbersome due to the short duration of almost all the episodes with three pulses. As mentioned in the main text, we distinguish only one of the four possible three-pulse configurations. This analysis leads to the coarse-grained state space *𝒮* = {*𝒮*_1_, *𝒮*_2_, *𝒮*_3_, *𝒮*_4_, *𝒮*_5_, *𝒮*_6_, *𝒮*_7_, *𝒮*_8_} (Fig. S6D), where the chiral single pulse corresponds to _3_, which is equivalent to *𝒮*_*rot*_ (see the main text for the other definitions). More trajectories, for different confinement values, are shown in Fig. S7.

We chose 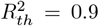 to define coherent rotations. While this value affects the classification quantitatively, the qualitative trends are preserved across different confinement regimes. As a consistency check, Fig. S8 shows how the average time spent at the rotating state *𝒮*_3_ before jumping to another state (average dwell time; see next Section VI) depends on *R*_*th*_ for different values of 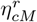.

## VI. Statistics in 𝒮-space

In the framework of the discrete dynamics in, we characterize the system in terms of dwell times 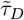 in each _*i*_ and the transition probabilities between states *P*(*𝒮*_*j*_ | *𝒮*_*i*_). Dwell times are recorded from 400 trajectories for each confinement parameter. In each trajectory, every visit to state *𝒮*_*i*_ contributes to the dwell time count, even if the visit is right-censored. We record which dwell times are uncensored versus right-censored and store them as separate distributions (see Fig. S9). From this statistical information, the survival probability 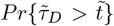 can be obtained with the built-in function of Matlab *ecdf()*, which computes the probability distribution 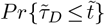 including rightcensored data via the Kaplan-Meier estimator (Fig S10). Moreover, mean dwell times 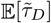 per state can also be extracted from the distributions in Fig. S9. In order not to underestimate this averaged quantity, we approximate the expectation by integrating the corresponding survival probabilities [13]:

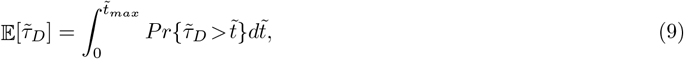

where 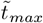is the largest time at which one can estimate 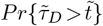. 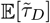 is a compact measurement of persistence in each state of the discrete system, which is particularly useful to study confinement effects on the maintenance of cell rotation (see Main text). In additional simulations, we have checked the influence of noise amplitude (Γ) on mean dwell times at *𝒮*_*rot*_ (Fig. S11).

From the observed uncensored visits per state, the transition probabilities are approximated by

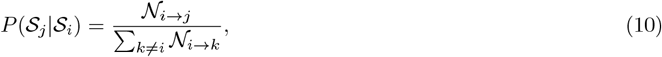

where 𝒩_*i*→*j*_ is the number of times a transition is observed from *𝒮*_*i*_ to *𝒮*_*j*_ and 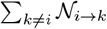 is the total number of exits from *𝒮*_*i*_. Transition probabilities smaller than 2% are set to zero, and the respective state probabilities are renormalized. The dependence of *P*(*𝒮*_*j*_|*𝒮*_*i*_) on the confinement parameter is illustrated in Fig S12.

Additionally, we estimate first passage times 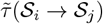 from the rotating state (*𝒮*_*rot*_) to all the other possible states (Fig. S13). The first passage time is numerically defined as the time spent in state *𝒮*_*i*_ before transitioning for the first time to another state *𝒮*_*j*_. Mean first passage times are calculated by building a survival probability for empirical first passage times 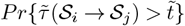, considering uncensored and censored (not reaching the target _*j*_ over the observation window given the start *𝒮*_*i*_) trips. Then, the expectation (mean) value is determined by the integration of this survival probability. In addition to the mean, we also report the 10^**th**^ and 90^**th**^ percentiles of the first-passage times. These are estimated from the probability distribution 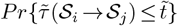, as the smallest 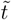 such that 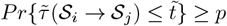, with *p* equal to 0.1 and 0.9, respectively. We do this to show the variability of the data, but also to stress the effects of heavy right-censoring, which sometimes forbids computing the 90^**th**^ percentile. The latter is a signature that expectation values are being underestimated. We also estimate the median (50^**th**^) to determine whether computing mean first passage times is meaningful.

**Figure S6:**
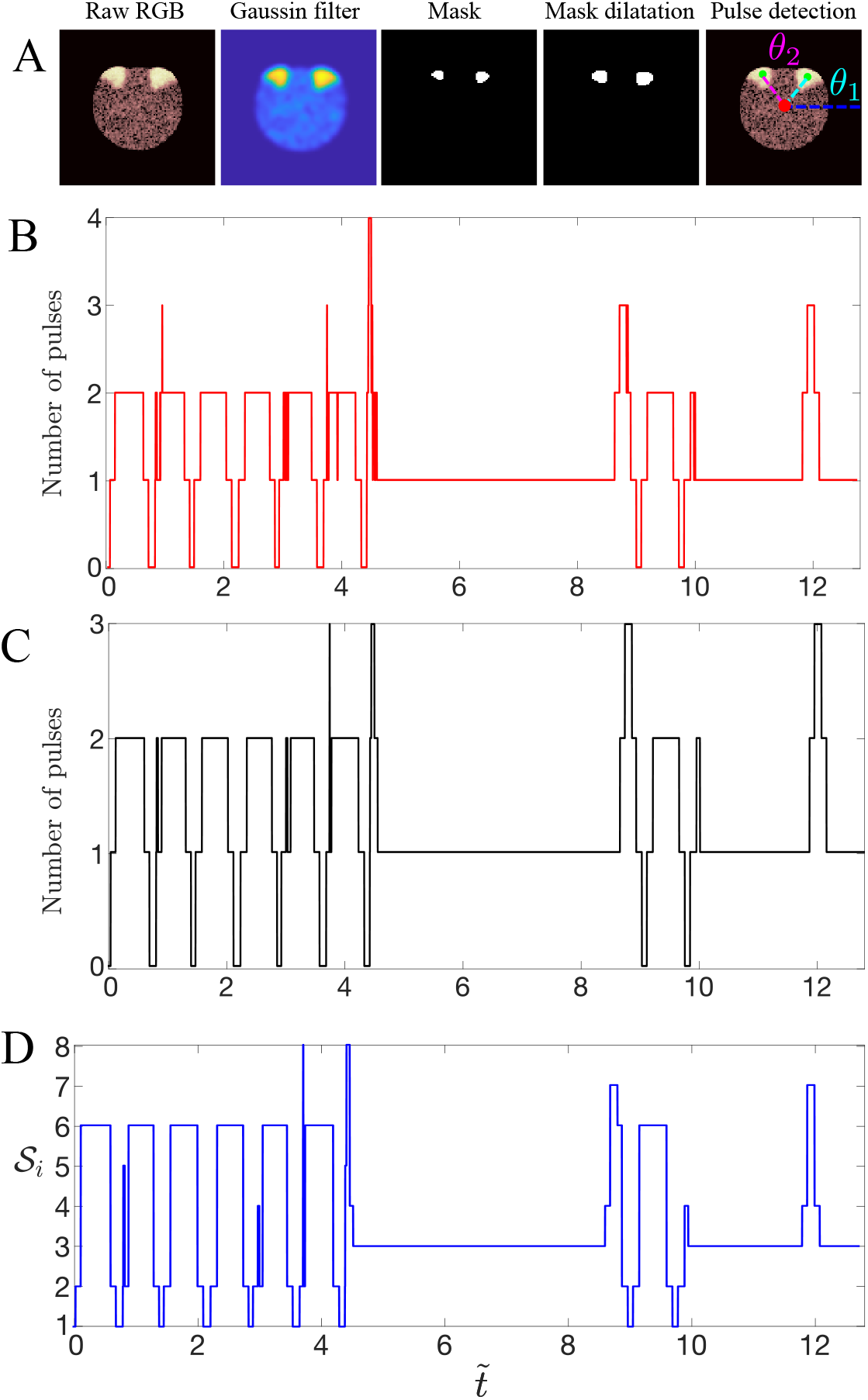
Detection of pulses. (A) Image process to extract the number of pulses (green circles) from the whole activator field. The red circle indicate the centroid of the cell. (B) Raw temporal trajectory of the number of pulses after image processing shown in (A). (C) Temporal trajectory of the number of pulses after removing spurious pulse counts. (D) Trajectory in *𝒮*-space. The confinement value for this particular trajectory is 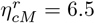. The tilde superscript represents normalization by the rotational period *T* of the rotating state, measured at 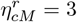.

**Figure S7:**
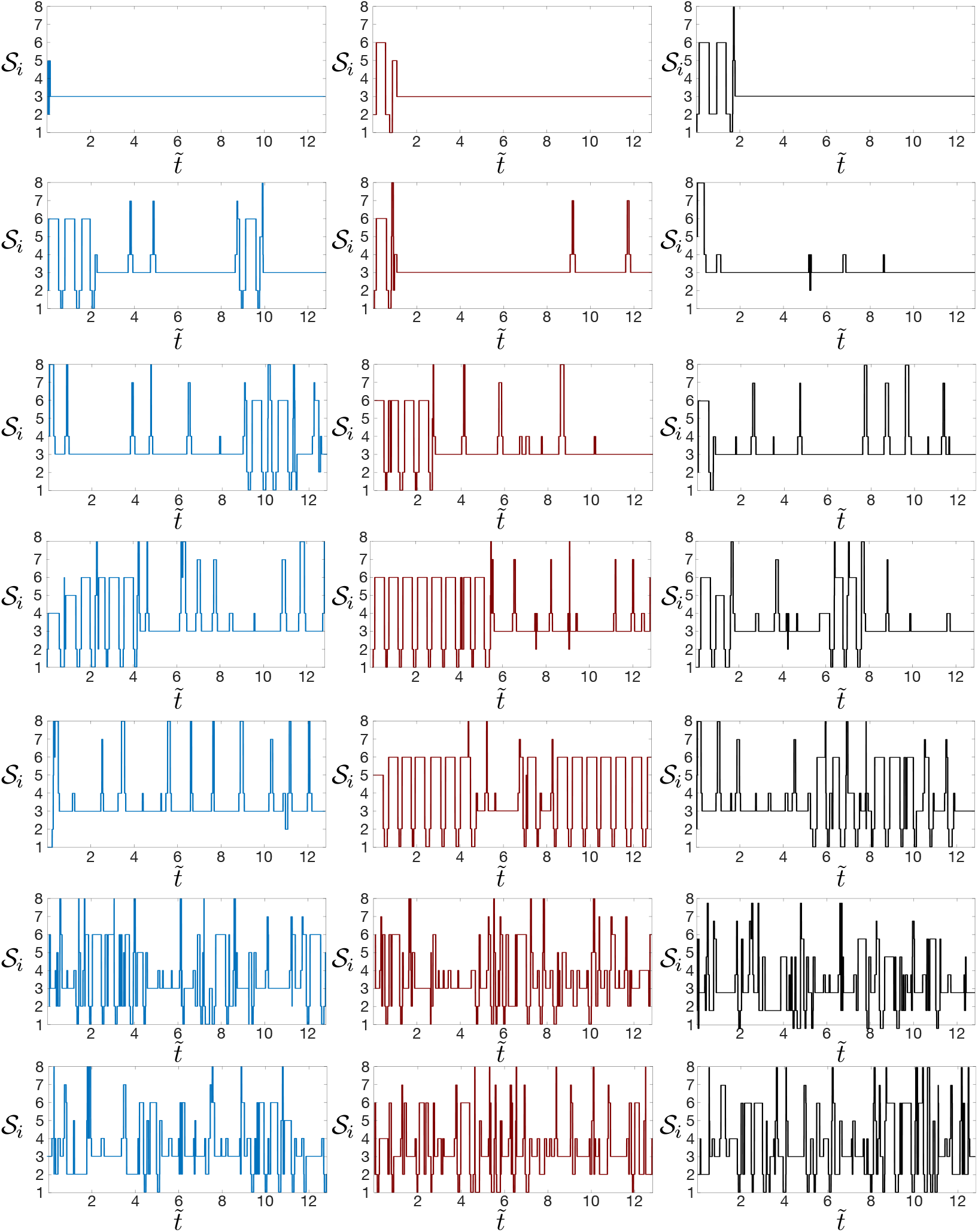
Three exemplary trajectories (columns) for each confinement 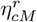 in the range [2, 32]. The tilde superscript represents normalization by the rotational period *T* of the rotating state, measured at 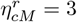

**Figure S8:**
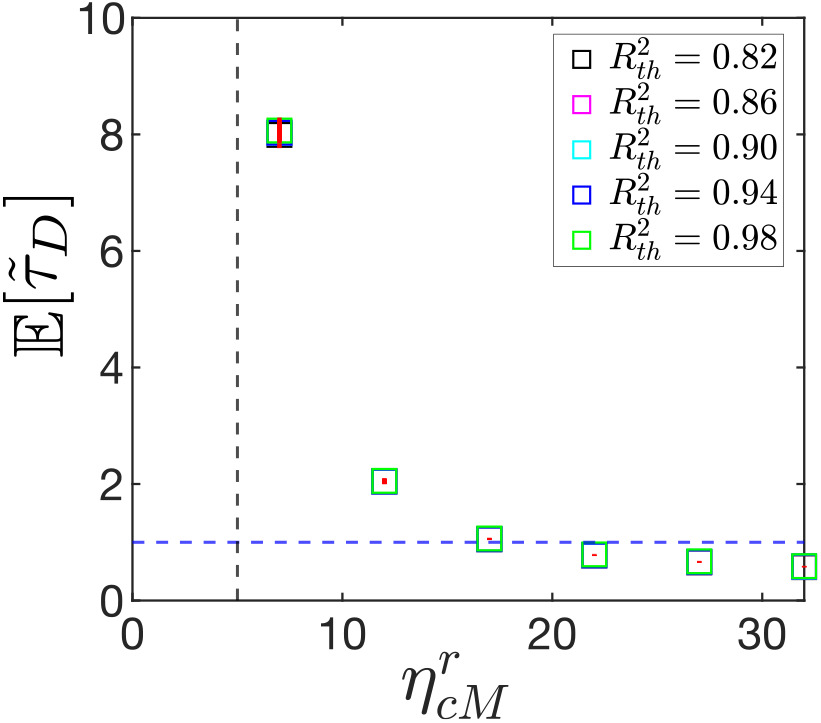
Effect of 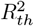 on the average dwell time at state *𝒮*_3_. The tilde represents normalization by the rotational period *T*. The solid red lines indicate the bootstrap standard error (SE) of the average dwell time, estimated from 500 resamples. The maximum SE is less than *T* /5. The blue dashed line labels one rotational period and the black dashed line accounts for the critical confinement 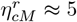.

## VII. Semi-Markov renewal process

Based on the dwell time statistics, we decide to introduce a semi-Markov renewal process as a data-driven model for our 2D system. We assume that an embedded Markov chain governs the transitions between states, i.e., the process is Markovian just before and after the transition occurs. However, the waiting times in each state before jumping to any other state are not exponentially distributed across confinements and across the whole observation time. Therefore, the whole stochastic dynamics is better represented by a semi-Markovian process. The renewal part comes from the assumption that upon arrival at a new state, the system resets and it is governed by the waiting time distributions and transition probabilities of that new state only. In this framework, one can write a simple linear equation for the mean first passage time between states *𝒮*_*i*_ and *𝒮*_*j*_, which compactly combines average dwell times and transition probabilities [14]:

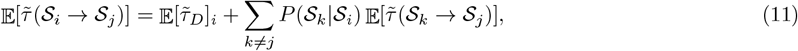

where the first passage times are 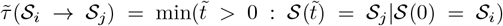. The first term accounts for the average waiting time in state *𝒮*_*i*_, while the second term considers all the possible transient trips *𝒮*_*i*_ → *𝒮*_*k*_ before absorption at *𝒮*_*j*_. The linear system (11) can be solved by fixing the target state *𝒮*_*j*_, extracting the matrix 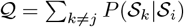, and inverting the matrix *I*−*𝒬*, where *I* is the identity matrix.

To test the predictive power of the renewal model, we compare the empirical and modeled mean first passage times. The semi-Markov renewal description reproduces fairly well 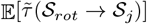 sufficiently far from 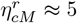 (Fig. S14). The differences are less than two rotational periods except at 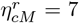. The significant mismatch at 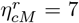 comes from a large number of censored trajectories at *𝒮*_*rot*_ (Fig. S13), which greatly underestimates the empirical mean first passage times. It also reflects that the renewal theory assigns a small probability of transitioning to *𝒮*_1_, and a high probability of remaining in the loop *𝒮*_*rot*_ ↔ *𝒮*_4_ (Fig. S12).

## VI. Supplemental figures for Fig. 3: Fig.S15 and Fig.S16

Upon confinement weakening, the localized protrusion that drives the rotational motion becomes larger, and eventually 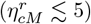, is able to translate the entire cell (Fig. S15). This morphological and dynamical change is associated with an enhancement of cell contraction far from the protrusion, which is sensible to variations in the strength of the size-restoring force (Fig. S16).

**Figure S9:**
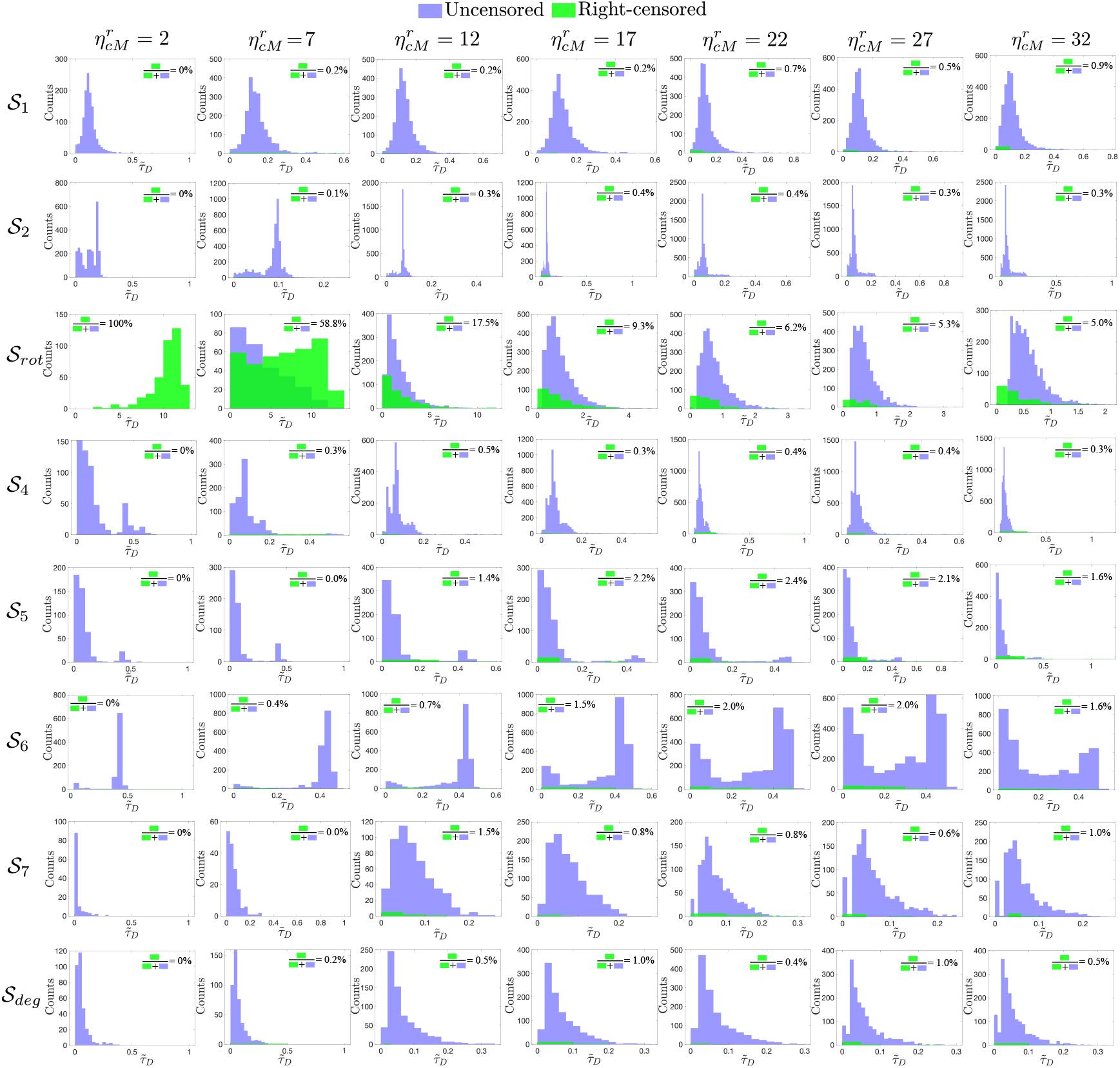
Distribution of dwell times 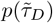 of the eight states of the discrete for different confinement values 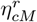.

## IX. Zero-dimensional representation of the back of the cell

In the main text, we investigate the role of the rotational protrusion in the dynamics of the rest of the cell. Particularly, due to trapping in the rotating state under weak confinement, a key question is how the excitable dynamics of the system is affected such that new activations (trips out of *𝒮*_3_) are not observed over finite times. This question is complex to address at the whole cell level, so we adopt a simplified approach. Excitations cannot be triggered at the location of the first pulse, but only at a finite distance from it. Then, a convenient and simple region to analyze the effects of activation suppression is at the back of the cell; farthest from the protrusion.

Activation suppression in the excitable system arises from cell mechanics, as reducing confinement modifies the force balance in Eq. (1). At the back, the cell experiences compression triggered by membrane extension at the protrusion site. This mechanical effect feeds back into the activator-inhibitor dynamics via the phase-field coupling in Eq. (2). In our simplified approach, we collapse Eq. (2) to a single point at the back by fixing *φ*_*c*_ = 1/2, defining local back variables {*A*_*b*_, *R*_*b*_} and neglecting spatial variations:

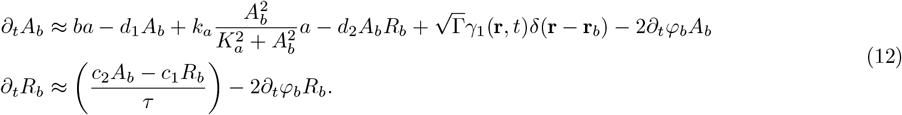

**Figure S10:**
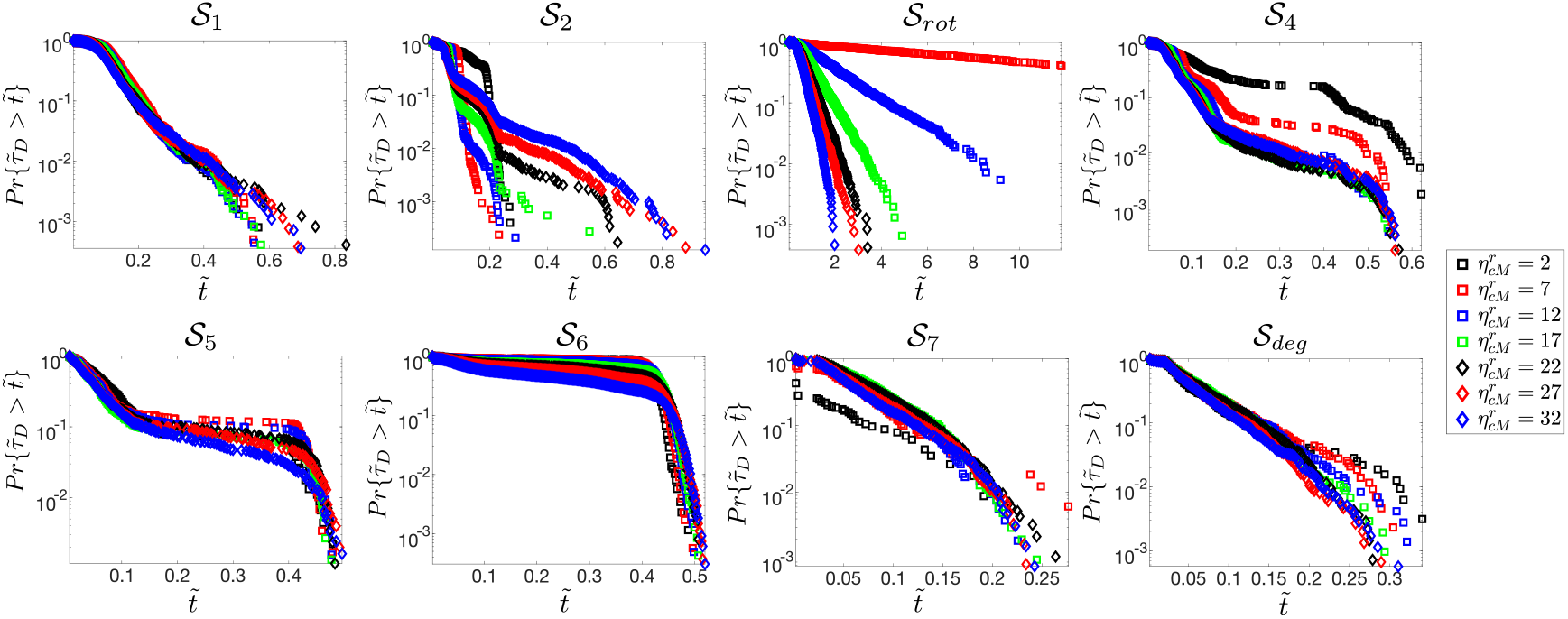
Survival curves 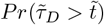 of the eight states of the discrete system for different confinements.

**Figure S11:**
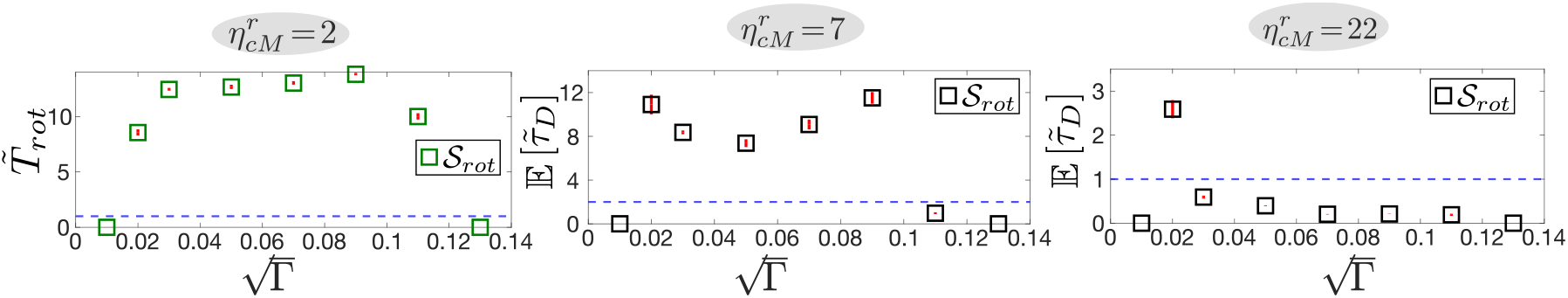
Characterization of noise amplitude variations in 2D simulations. The sweep of 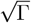 is is in steps of 0.02. There is a minimum noise necessary to trigger transitions out of 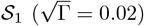 and a maximum amount of noise that does not allow the necessary coherence to create protrusions 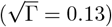. 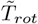 is the average time spent in the rotating state at 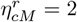, where all the trajectories are right-censored. The large 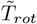 is robust over a range of noise amplitudes. Interestingly, the behavior of 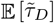 at 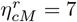 as a function of 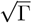 is bimodal. The second peak for relatively high noises is correlated with a high chance of reaching *𝒮*_*rot*_ from *𝒮*_1_, but a low chance of going out. All the results in the main text are reported for 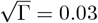.

The corrections arising from the phase field coupling, proportional to 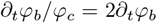, incorporate boundary deformation effects on the excitable dynamics in the form of back contractions: 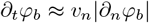 (using Eq. (2) of the main text), where the subscript *n* indicates the normal direction at the interface and *v*_*n*_ *<* 0. Additionally, the term ∂_*t*_*φ*_*b*_ serves as a proxy for the advection correction localized at the boundary in the sharp interface limit [15]. The reported value of ∂_*t*_*ϕ*_*b*_ is a spatiotemporal average over 2*T* rotations and over an arc—centered at the interface point farthest from max(*A*) and spanning approximately one quarter of the cell perimeter 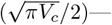obtained from 2D numerical integrations of Eqs. (2) and (4) in the noiseless case (Γ = 0), using a stable rotating state as the initial condition (Fig. 3; main text). The modification of the underlying excitable activator-inhibitor dynamics can be quantified by computing the corrected fixed point of Eq. (12), and a comparison between the corrected and numerically measured fixed points at the back 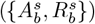 reveals the same decreasing trend with decreasing confinement (Fig. S17A). In phase space, the change upon confinement (or contraction) can be simply characterized by the distance 𝒟 from the fixed point to the minimum of the cubic nullcline (Fig. S17B). The trend in 𝒟 as a function of 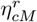 indicates that, when the cell is less confined (and more contracted), the activator-inhibitor dynamics becomes less excitable.

**Figure S12:**
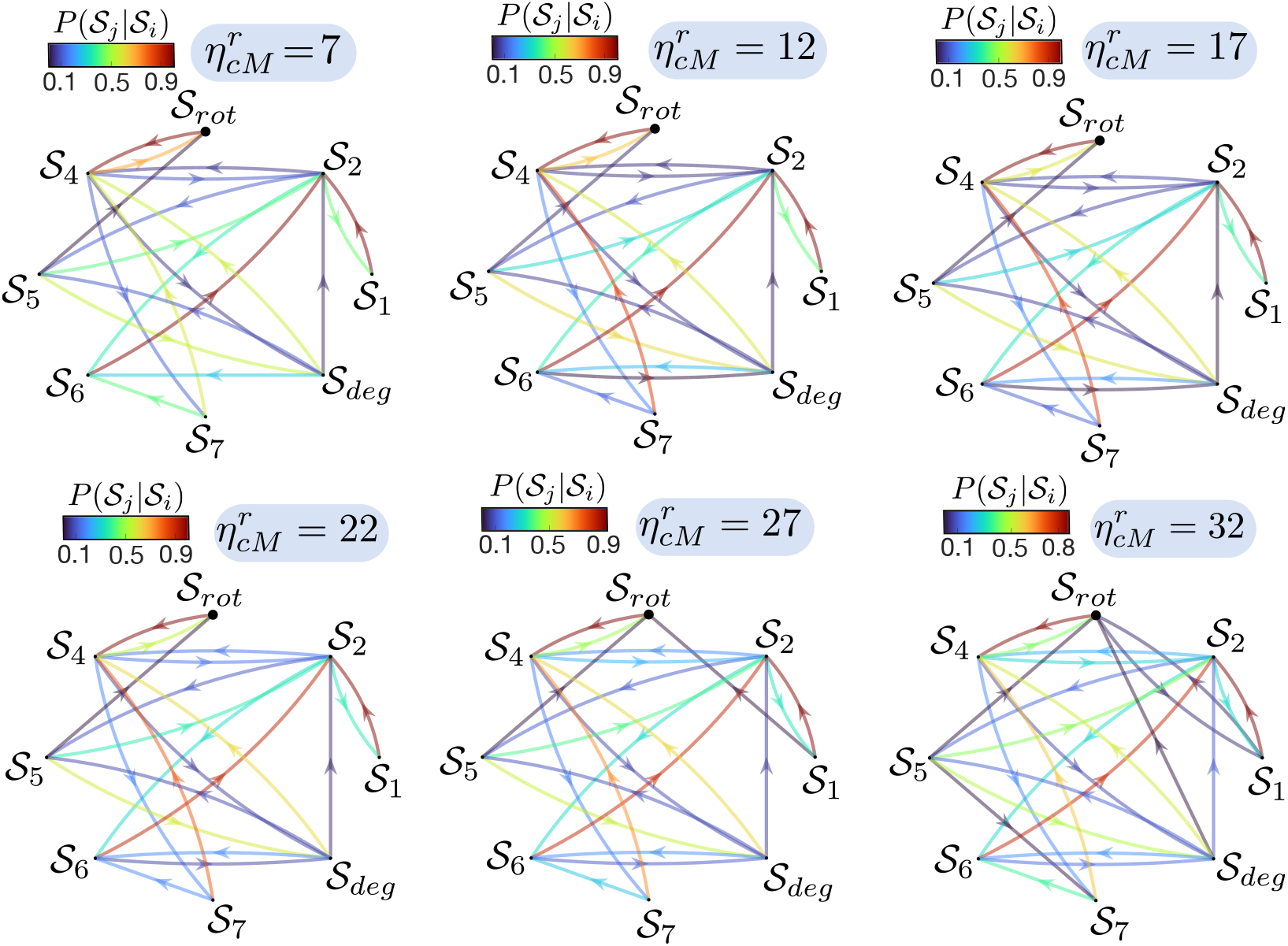
Transition probabilities in the discrete space *𝒮* for different confinement values 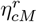.

## X. Quasi-steady approximation and Kramers escape

To connect the statistical analysis performed in this study with the simplified back of the cell approach, we assume a quasi-steady approximation for the activator-inhibitor dynamics and freeze the slow inhibitor dynamics in time, setting 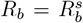, i.e., *τ* ≫ 1 in the dimensionless version of Eq. (12). The mechanical correction in the dynamics of the back inhibitor does not introduce additional time scales (*c*_2_ ~ *τ*| ∂_*t*_*φ*_*b*_|). Under this approximation, the back activator-inhibitor dynamics can be recast into a compact Langevin equation in the Ito representation

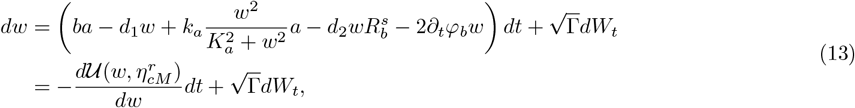

where *W*_*t*_ is a Wiener process [16], and for simplicity *w* = *A*_*b*_. Moreover, this Langevin equation is variational with potential:

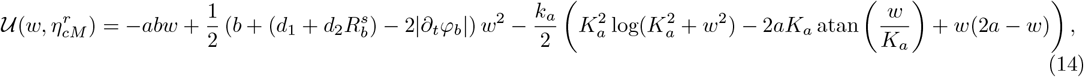

where the confinement dependence comes via ∂_*t*_*φ*_*b*_. The problem is thus reduced to a particle in a potential well that may escape over the potential barrier due to stochastic fluctuations. This is the well-known Kramers escape problem, in which the escape rate ℛ from the potential well—defined as the conditional probability per unit time that a particle escapes, given that it is in the potential well minima—can be computed [16]. For this, we introduce the corresponding Fokker-Planck equation for the probability density *P*(*w, t*):

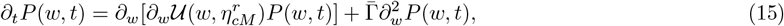

with 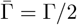. This drift-diffusion equation for the probability density can be written in flux form:

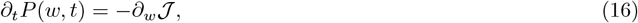

Where

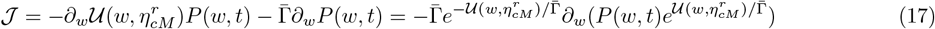

can be interpreted as a probability flux over the potential barrier. Therefore, this flux is related to ℛ by the simple relationship ℛ *p* = 𝒥, where *p* is the probability of being near the potential minimum 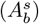:

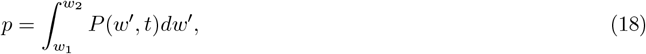

where *w*_1_ and *w*_2_ are arbitrary points inside the potential. If the barrier height, 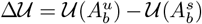, is much larger than 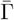, which holds for sufficiently weak confinements and the noise amplitudes considered in our study (Fig. 3C and Table S1), the probability distribution near 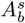 can be approximated by the stationary distribution:

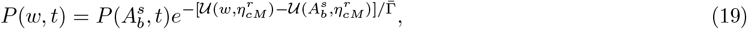

which is obtained from Eq. (17) with 𝒥 = 0 and normalizing by the probability distribution at 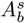. To obtain an expression for 𝒥 and estimate ℛ, we assume that the system is near steady state, and thus, one can consider the limit ∂_*t*_*P*(*w, t*) ≈ 0, which implies that the flux 𝒥 is independent of *w*, and integrate Eq. (17) from 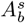 to 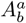, to obtain

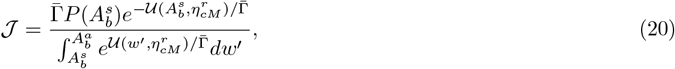

where 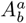 is an arbitrary point outside the potential in which 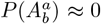 and 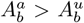. Combining the expressions for 𝒥 and *p*, considering the stationary probability distribution at 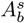, the Kramers escape rate reads

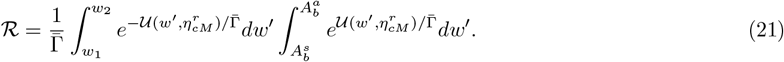

**Figure S13:**
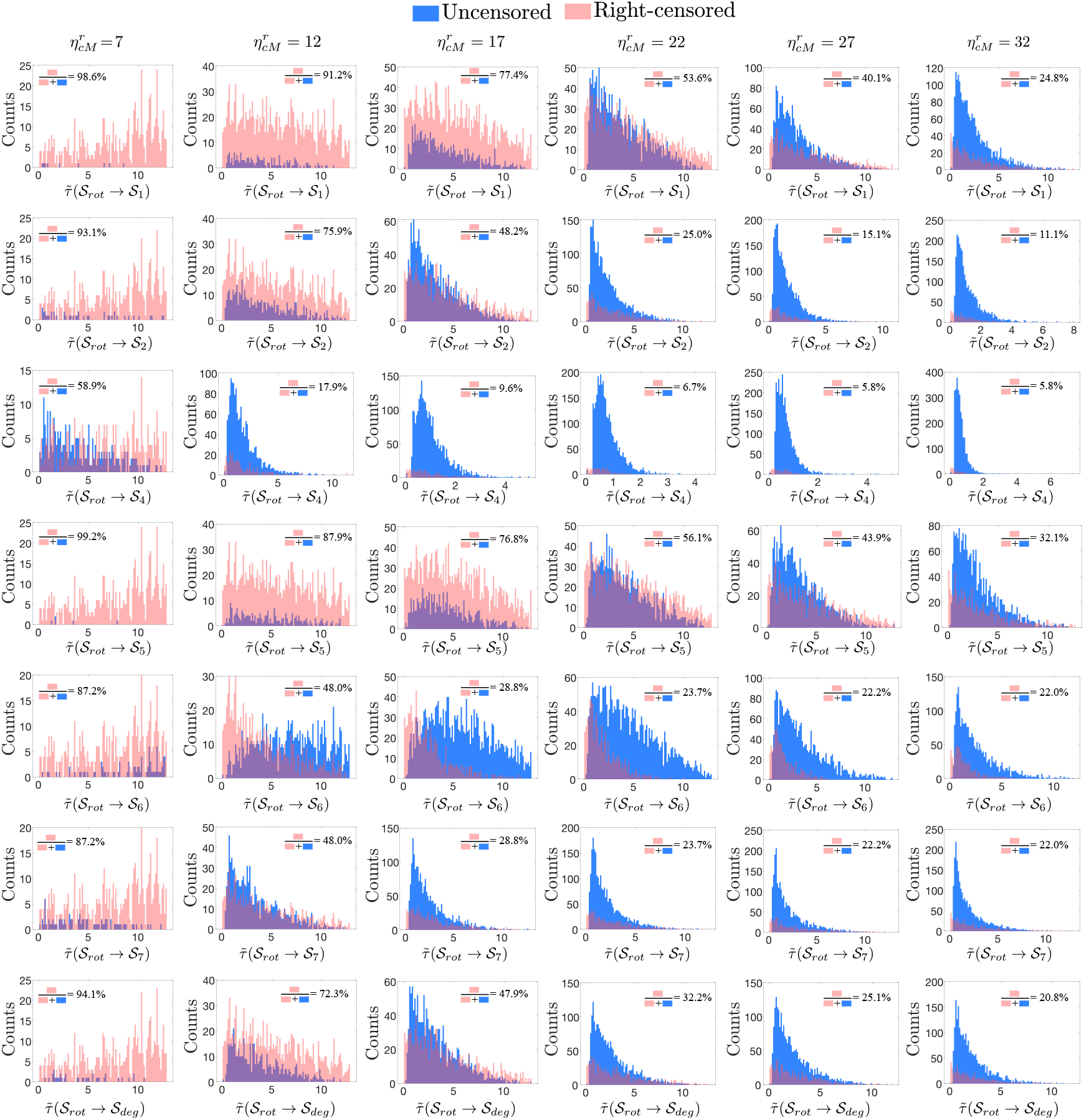
Distribution of first passage times from the rotating state to the rest of states for different confinement values 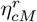.

**Figure S14:**
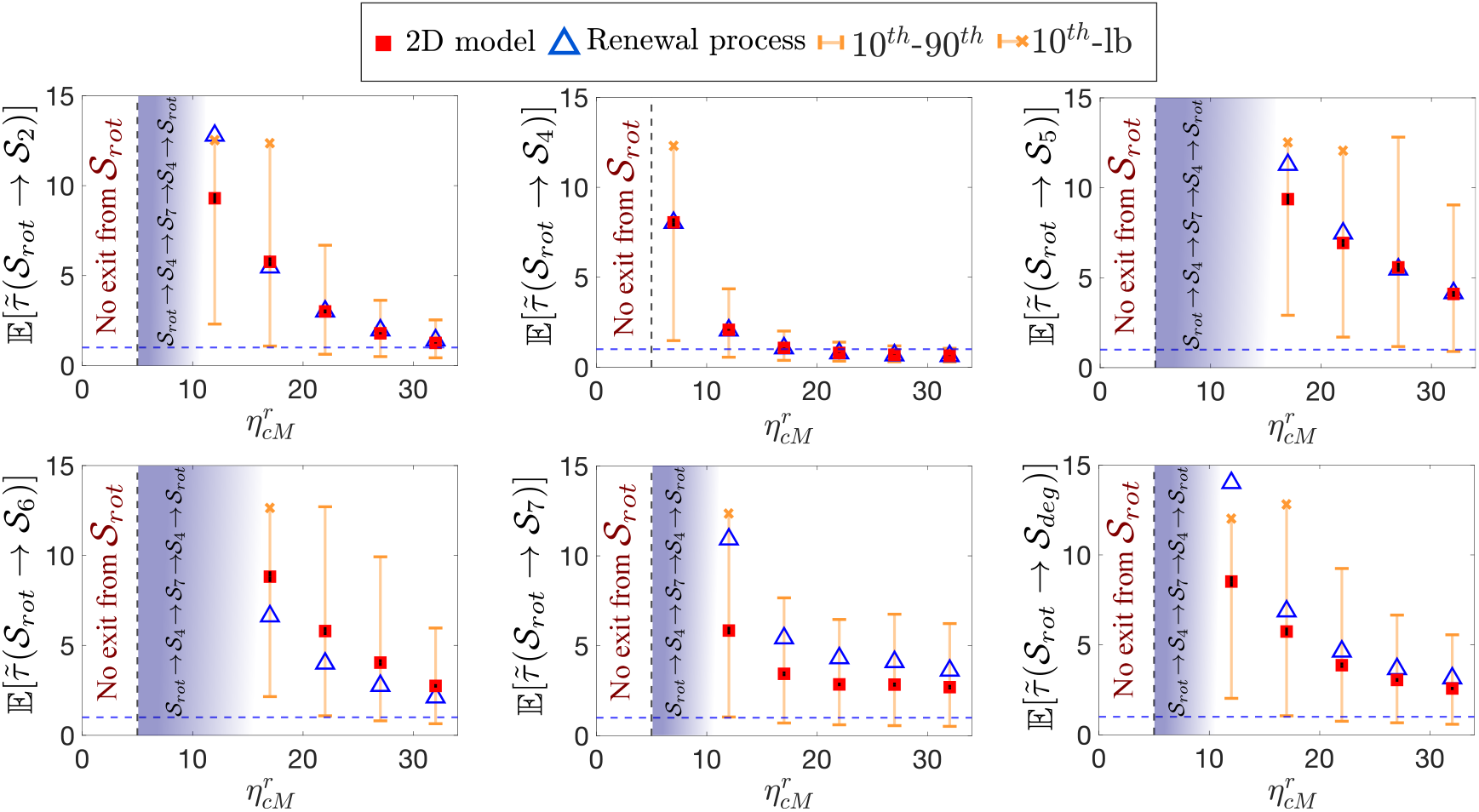
Mean first passage times from the rotating state to the rest of the states except for *𝒮*_1_ (see Main text), computed using the 2D phase field model (squares) and the semi-Markov renewal process (triangles) as a function of 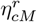. The orange whiskers correspond to the 10^*th*^ and 90^*th*^ percentiles of the first-passage time distributions. The orange (x) symbols indicates the lower bound (maximum first passage time observed) when a 90^*th*^ cannot be estimated. Bootstrap standard errors (500 trajectory resamples) of all the 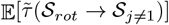 are indicated by red whiskers

**Figure S15:**
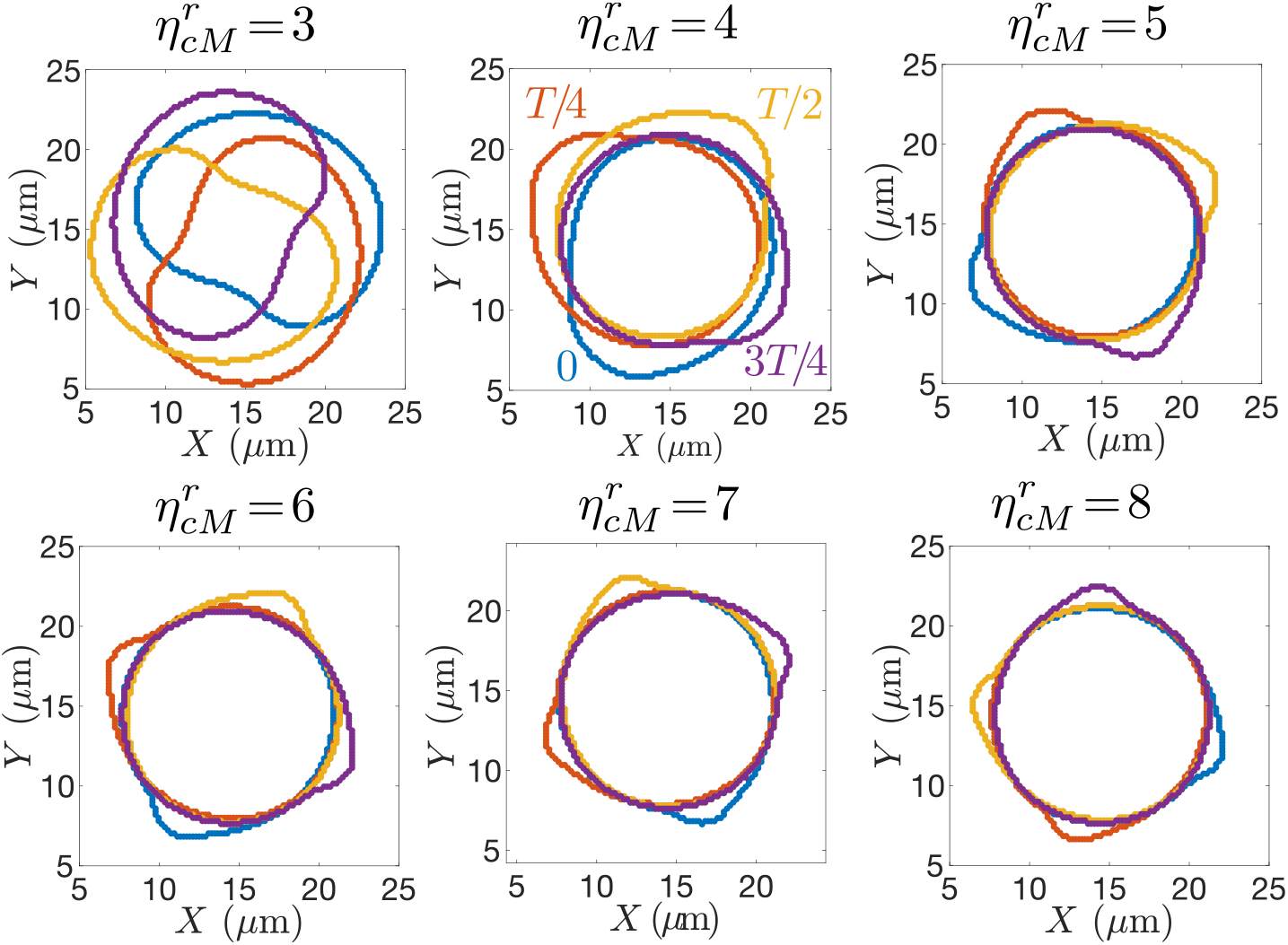
Temporal evolution during one full rotation, shown every *T* /4, of the cell’s interface *φ*_*c*_ = 1/2 near the transition 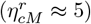 from intermediate to weak confinement regime. Below the critical confinement 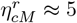, the protrusion becomes large enough to contract the entire cell, resulting in rigid-body motion. These simulations are noiseless (Γ = 0).

**Figure S16:**
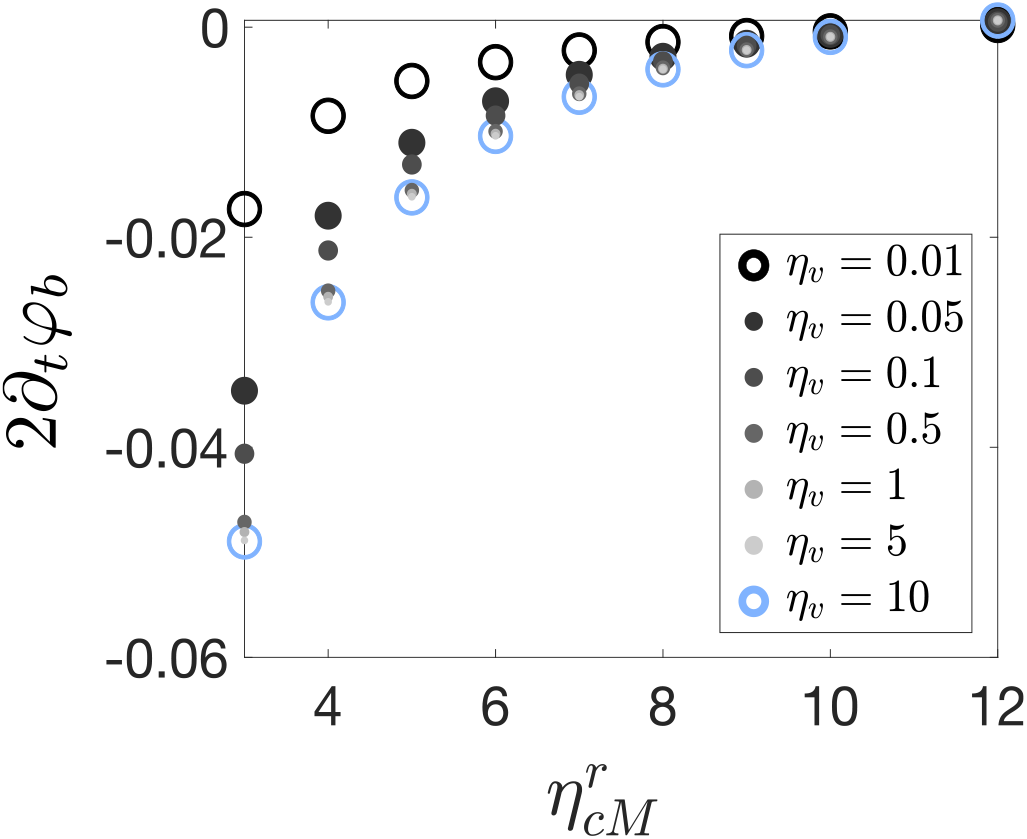
Changes in the contraction effect at the back of the cell, ∂_*t*_*φ*_*b*_/*φ*_*c*_ = 2∂_*t*_*φ*_*b*_, for different values of the strength of the restoring-size force *η*_*v*_ over a range of confinements.

**Figure S17:**
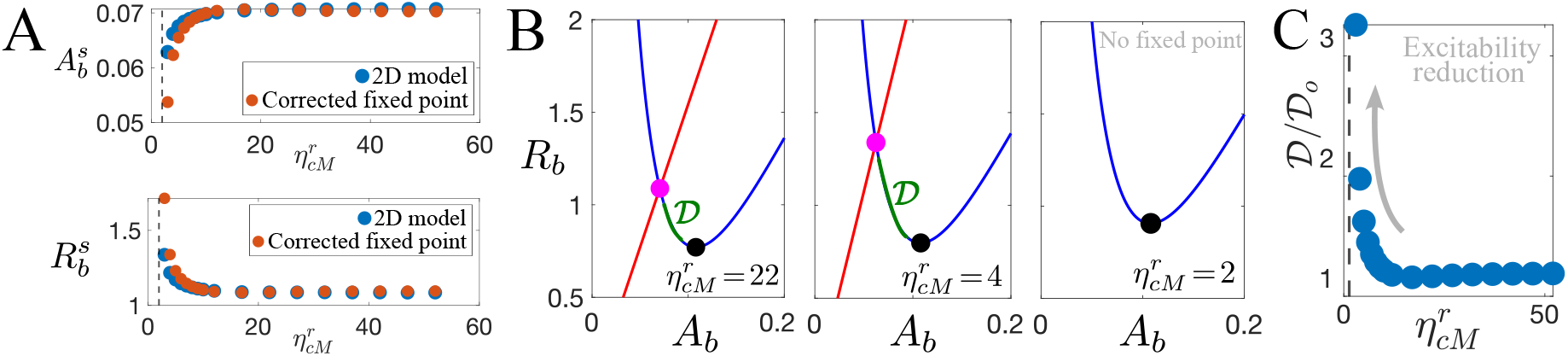
Activator-inhibitor dynamics at the back of the cell. (A) Comparison between the back fixed point 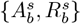 obtained from numerics (2D model) and from Eq. (12) for different values of 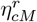. (B) Phase space visualization of the fixed point shift as confinement weakens. The distance 𝒟 between the fixed point (magenta) and the minimum of the cubic nullcline (black) is highlighted in green. (C) Growth of 𝒟 until the fixed point disappears 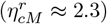. The distance 𝒟 _*o*_ is measured under homogeneous conditions (∂_*t*_*ϕ*_*b*_ = 0).

One can solve both integrals in Eq. (21) using Taylor expansions, up to second order, of the potential well 𝒰around 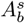 for the first integral, and around 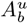 for the second integral. The rate is then given by

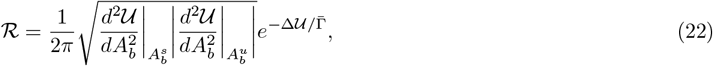

where

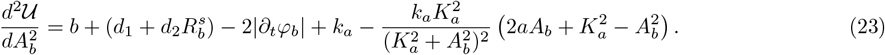

Finally, the inverse of ℛ can be interpreted as the mean escape time 𝒯 from the potential well. Fig. 3D of the main text shows that this escape time increases as confinement weakens, consistent with the reduction of excitability triggered by contraction.

## XI. Experimental setup

MCF10A mammary epithelial cells tranduced with LifeAct-GFP and H2B-mCherry were cultured in growth factor reduced matrigel (Ref: 356231, Phenol red free, Corning). Briefly, MCF10A cells, at a seeding density of 500 cells per *µ*L, were mixed with Matrigel at ratios ranging from 30% to 100% and cast into a 96-well glass-bottom plate (Matek Corporation). The gels were allowed to crosslink at 37^*?*^C for 45 minutes. After 2 h of culture, 35 *µ*m Z-stacks were acquired every 20 min for a total duration of 16 h using a Nikon TiE inverted microscope. The captured timelapses were then post processed using Bitplane Imaris software.

The Z-stacks were manually examined to assess the presence of cell rotation, defined as visible chiral motion persisting for 0.25*T* and accompanied by a distinctive F-actin patch on the cell surface. The identification was independently performed by three users, resulting in perfect agreement for Matrigel concentrations of 50%, 60% and 80%, and slight variable agreement for 70% and 100%. This variability is reported as standard deviations in Fig. 4D of the main text.

## XII. Equivalence between weak confinement and extra free space

In our model, confinement weakening is analogous to increasing the effective cell-ECM distance, allowing protrusions to develop more easily. To illustrate this, we perform a weakly nonlinear analysis around the steady state solution of a one-dimensional representation of the cell (similar to the lower panel in Fig. S1A). We assume that 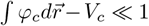, and thus focus only on the effect of confinement and protrusions:

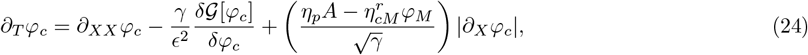

where space and time have been rescaled as 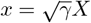 and *t* = *ξT*. Additionally, we assume that the activator dynamics is much faster than the mechanical deformations of the cell membrane such that we can ignore the temporal dependence of *A* in Eq. (24). Because of the symmetry and the simplified size restoring force, we analyze the dynamics locally at one end of the 1D-cell; without loss of generality, we choose the right side of the 1D profile (cf. lower panel in Fig. S1A). In the absence of confinement 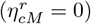 and protrusions (*η*_*p*_ = 0), and at the Maxwell point of the variational system, the 1D phase field is characterized by the homogeneous profile 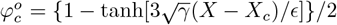, where *X*_c_ is the cell boundary. A similar one-dimensional profile is used for the static ECM: *φ*_*M*_ = {1 + tanh[*q*(*X* − *X*_*M*_)]}, where *X*_*M*_ is the ECM boundary, and for simplicity we choose 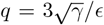. The aim is to track the movement of the cell boundary through the first nonlinear correction *W* introduced by perturbations from a protrusion and confinement: 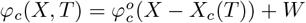. Substituting this ansatz in Eq. (24) yields, at *𝒪* (*W*),

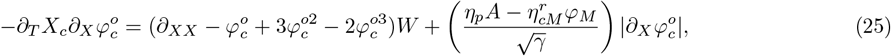

where we assume 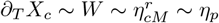 are small. This equation is linear in *W*, ℒ *W* = *b*, with

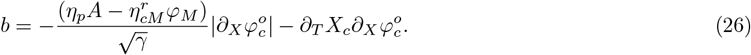

Under the usual inner product ⟨*f, g*⟩ = *fgdX*, the linear operator is self-adjoint, ℒ = ℒ ^*T*^, with kernel 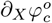. Then, Eq. (25) is solvable if and only if 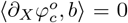. This solvability condition leads to the following expression for the one-dimensional velocity of the cell boundary:

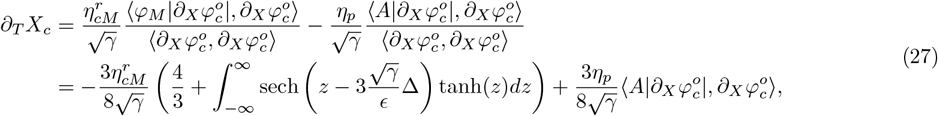

where *z* = *X* − *X*_*c*_ and Δ = *X*_*c*_ − *X*_*M*_. The last term, proportional to *η*_*p*_, is always positive and gives the first order contribution of the protrusive force. Let us focus on the first term in Eq. (27), proportional to 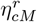, which gives the first order contribution to confinement. The integral term, denoted ℐ, is difficult to evaluate analytically but can be interpreted straightforwardly. When Δ = 0, ℐ is zero by parity. For Δ ≠ 0, the sign of Δ controls the sign of the integral, sgn(ℐ) = sgn(Δ). Therefore, when extra space is available to the cell (Δ *<* 0), the term in parentheses in Eq. (27) decreases, effectively weakening the confinement. Reintroducing the spatial and temporal scales, Eq. (27) becomes

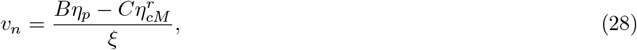

where *B* and *C* are positive constants.

## XIII. The oscillatory regime of the activator-inhibitor dynamics

While the excitable regime is most appropriate for our study, the oscillatory regime of actin wave dynamics has also been reported in the literature [17–19], and previous numerical studies of cell migration within the phase field framework have focused on this regime of the activator-inhibitor dynamics [2, 3]. Therefore, for completeness, we examine the consequences of the oscillatory dynamics of *A* and *R* on the behavior of the rotating states under confinement. To this end, we reduce the value of the activator degradation rate, *d*_1_ (see Supplemental Material), by a factor of 2, which drastically changes the dynamics (Fig. S18). In this regime, the homogeneous steady state of the system is unstable, the intersection of the linear and cubic nullcline occurs at the middle branch of the cubic one, and the activator-inhibitor dynamics is governed by a limit cycle (there is no resting state). We numerically integrate the two-dimensional phase field model across a range of confinement parameter values. We again find that below a critical confinement 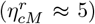, the dynamics is deterministic: once the system reaches the rotating state, it remains there without switching due to the strong back contraction. Above this critical confinement, specifically within the range 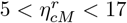, the dynamics is modified and an oscillatory *back and forth* protrusion mode emerges (Video S13). This mode is remarkably stable, and our results show that once the system access it, it never leaves (across 100 realizations of the noise with total time 13*T*). Nevertheless, the system may also spend the full time (13*T*) exhibiting purely stochastic dynamics, similar to what is observed in the excitable regime. We hypothesize that the upper limit 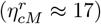 for the existence of the oscillatory mode arises because the system needs to sustain a minimal protrusion size to maintain the *back and forth* motion. The three-dimensional analogue of this mode is a persistent target-wave pattern (Video S14). Together, although the route to persistent chiral motion as confinement weakens depends on whether the activator-inhibitor dynamics is excitable or oscillatory, chiral symmetry breaking remains a robust feature in cells under weak confinement.

## XIV. Sensitivity analysis of the local activator-inhibitor dynamics

The parameters governing the local dynamics in Eq. (2), {*a, b, d*_1_, *d*_2_, *k*_*a*_, *K*_*a*_, *c*_1_, *c*_2_}, determine the fixed point of the nonlinear system (the intersection between the linear and cubic nullclines; Fig. S1C). If this intersection lies too far from the minimum of the cubic nullcline—quantified by the distance 𝒟 (see Section IX—the reduction in excitability can suppress activator nucleations and therefore coherent rotations.

**Figure S18:**
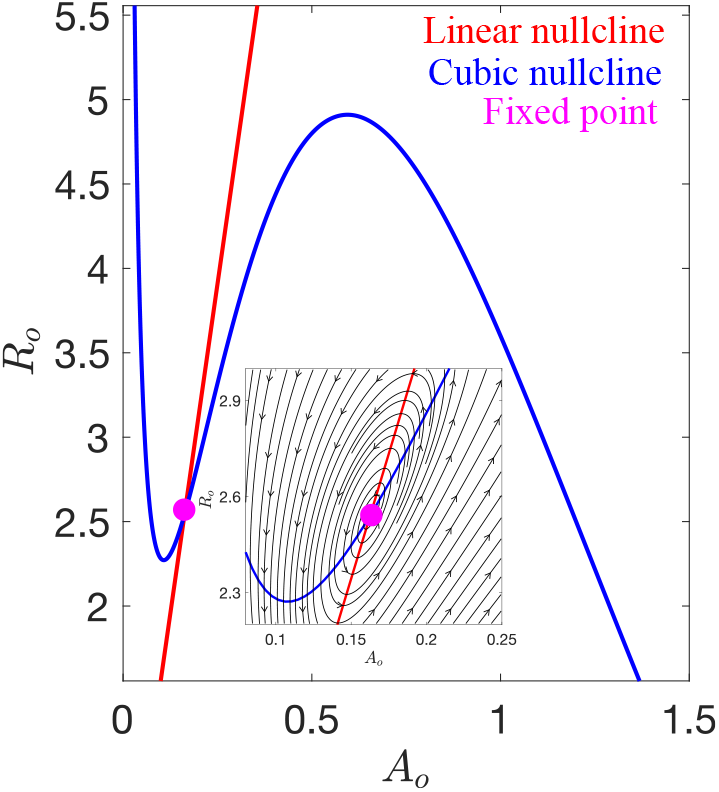
Phase space of the zero-dimensional version of the activator-inhibitor dynamics in Eq. (2), given by {*A*_*o*_, *R*_*o*_}, in the oscillatory regime. The inset shows the structure of the phase flows around the fixed point (repeller).

We vary the parameters {*a, b, d*_1_, *d*_2_, *k*_*a*_, *K*_*a*_, *c*_1_, *c*_2_} one by one within ±50% of their original value (Table S1) and verify that any such variation produces coherent rotations provided that 𝒟 ≲ 0.32 (blue shaded regions in Fig. S19). For larger 𝒟, the system is unable to produce activator pulses (pink shaded regions in Fig. S19). Parameter changes can also induce a bifurcation from the excitable to the oscillatory regime (green shaded regions in Fig. S19). The parameter *τ* in Eq. (2), which sets the time scale of the inhibitor, does not affect the location of the intersection in phase space. However, it cannot be arbitrarily small (i.e., fast time scale), as this can prevent the emergence of activator pulses. We have checked that coherent rotations are still present when varying *τ* within ±25% of its original value.

**Figure S19:**
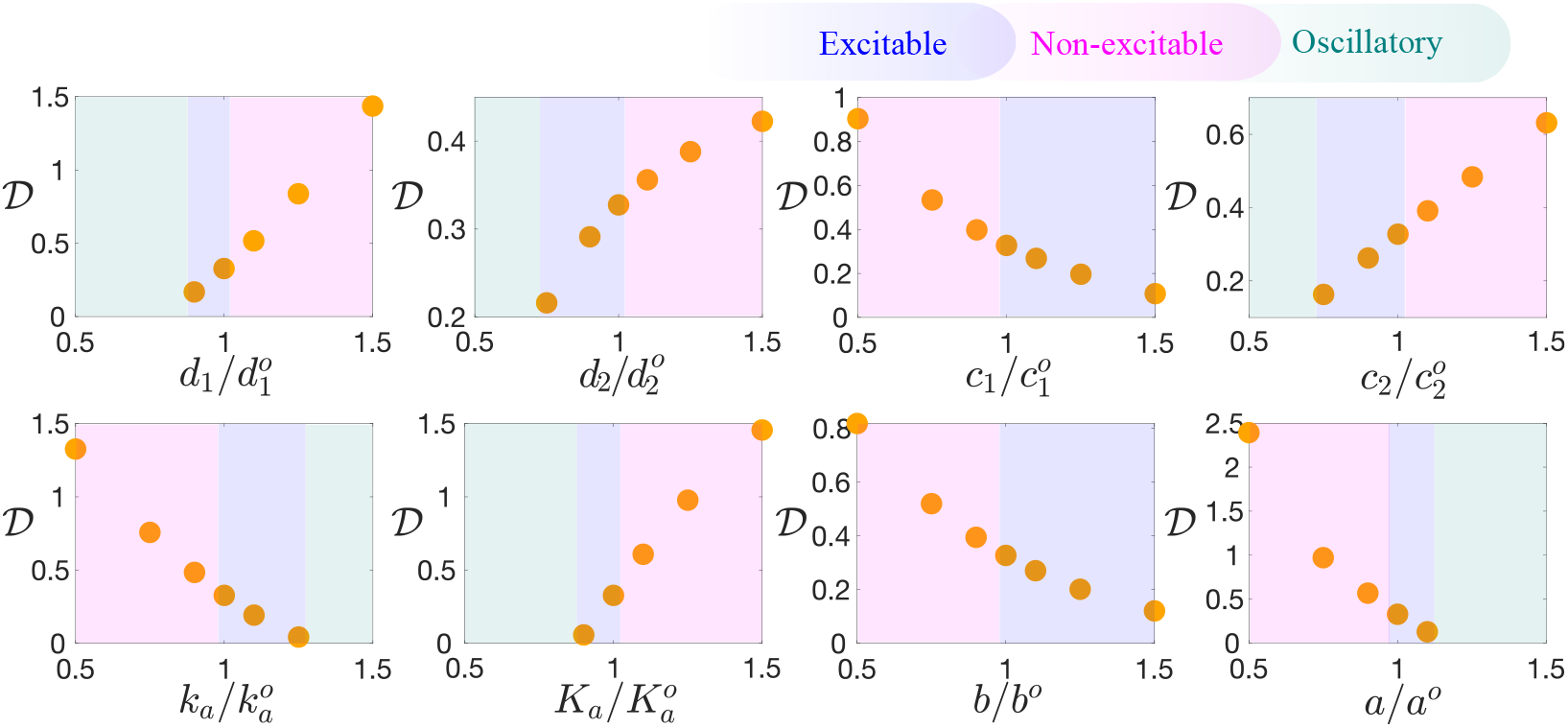
Variations in 𝒟 as a function of eight parameters controlling the wave dynamics in the activator-inhibitor system of Eq. (2), and in the weak confinement regime 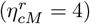. The superscript ° denotes the original parameter value in Table S1.

## XV. Supplemental figures for Section III.D of the main text: Fig. S20 and Fig. S21

Figs. S20 and S21 illustrate how the mean dwell time in the rotating state depends on variations in the diffusion coefficient *D*_*A*_ and the friction coefficient *ξ* across the confinement values analyzed in the main text.

**Figure S20:**
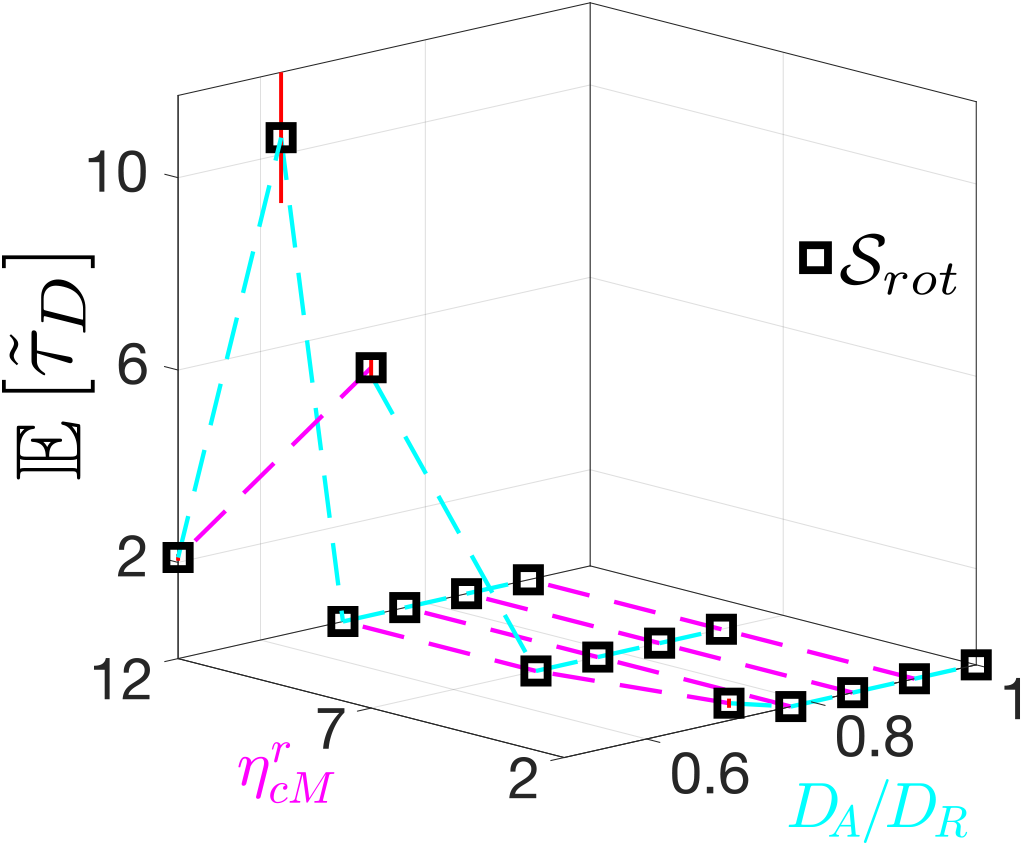
Phase diagram in 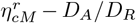 space showing the computable mean dwell times in the rotating state.

**Figure S21:**
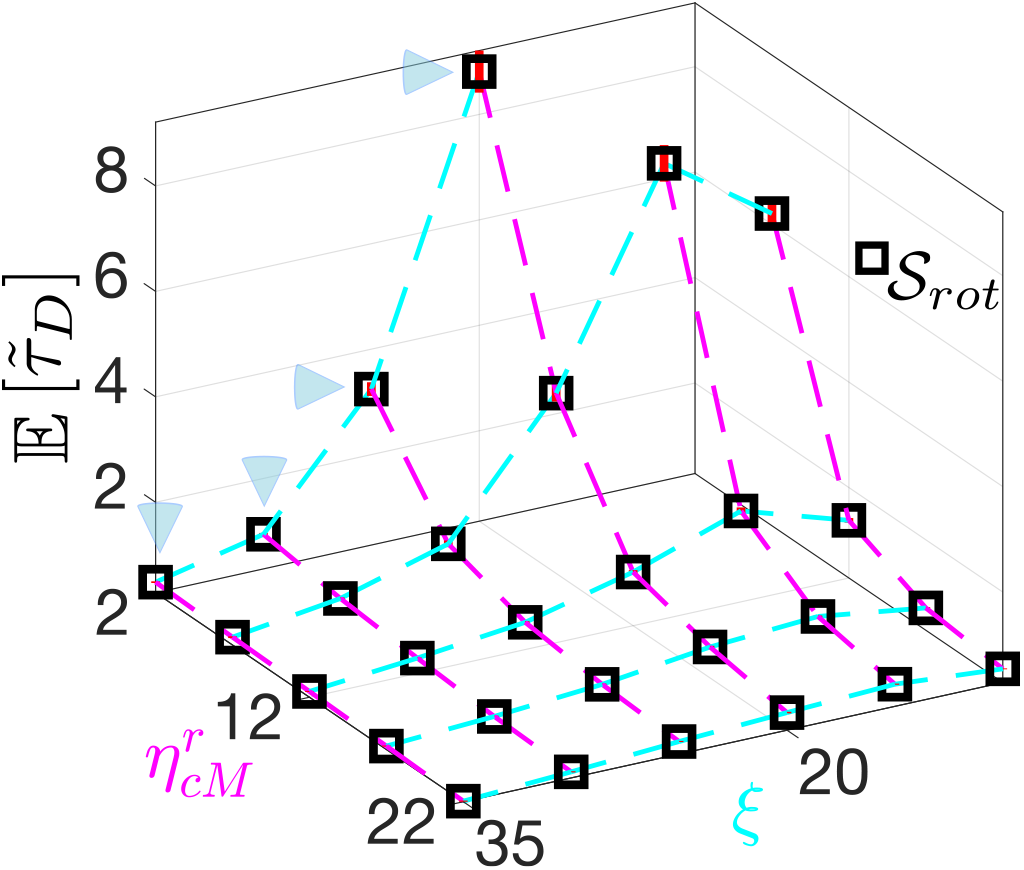
Phase diagram in 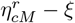 space showing the computable mean dwell times in the rotating state. The blue triangles indicate measurable mean dwell times at 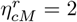 (exit from *𝒮*_*rot*_ is possible).

## XVI. Video captions

**Video S1**: Numerical integration of Eqs. 2 and 4 in 3D with 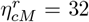. The color code indicates the normalized activator *A*/ max(*A*). The total time of integration is 66*T*.

**Video S2**: Numerical integration of Eqs. 2 and 4 in 3D with 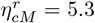. The color code indicates the normalized activator *A*/ max(*A*). The total time of integration is 66*T*.

**Video S3**: Numerical integration of Eqs. 2 and 4 in 3D with 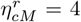. The color code indicates the normalized activator *A*/ max(*A*). The total time of integration is 66*T*.

**Video S4**: Interface dynamics (*φ*_*c*_ = 1/2) shown across an arbitrary plane, extracted from a segment of **Video S1**. The color code indicates the normalized activator *A*/ max(*A*).

**Video S5**: Interface dynamics (*φ*_*c*_ = 1/2) shown across a plane 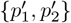, extracted from a segment of **Video S2**. The plane is chosen to coincide with an observed episode of coherent rotation. The color code indicates the normalized activator *A*/ max(*A*).

**Video S6**: Interface dynamics (*φ*_*c*_ = 1/2) shown across a plane {*p*_1_, *p*_2_}, extracted from a segment of **Video S3**. The plane is defined by the cell symmetry axis after reaching for the first time the state of coherent rotation. The color code indicates the normalized activator *A*/ max(*A*).

**Video S7**: Numerical integration of Eqs. 2 and 4 in 2D with 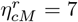 exhibiting the representative *𝒮*_*i*_ states and their transitions. The color code indicates the activator *A* in RGB scale. The total time of integration is 13*T*. The green circles show pulse positions (see Section V).

**Video S8**: F-actin dynamics on MCF10A cell surface in 100% Matrigel showing states *𝒮*_1_, *𝒮*_2_, *𝒮*_*rot*_ and *𝒮*_6_. F-actin is visualized with GFP-tagged LifeAct (green) and the nucleus is visualized with mCherry-tagged H2B (orange).

**Video S9**: Rotating (*𝒮*_*rot*_) MCF10A cell in 100% Matrigel. F-actin is visualized with GFP-tagged LifeAct (green) and the nucleus is visualized with mCherry-tagged H2B (orange).

**Video S10**: F-actin dynamics on MCF10A cell surface in 70% Matrigel showing state *𝒮*_5_. F-actin is visualized with GFP-tagged LifeAct (green) and the nucleus is visualized with mCherry-tagged H2B (orange).

**Video S11**: Translational non-rotating MCF10A cell in 100% Matrigel. F-actin is visualized with GFP-tagged LifeAct (green) and the nucleus is visualized with mCherry-tagged H2B (orange).

**Video S12**: Extract of a numerical integration of Eqs. 2 and 4 in 2D with 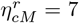 showing a short episode with four pulses. The color code indicates the activator *A*. The green circles show pulse positions (see Section V).

**Video S13**: Extract of a numerical integration of Eqs. 2 and 4 in 2D with 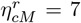 = 7 showing the *back and forth* mode in the oscillatory regime (*d*_1_ = 1.5). The color code indicates the activator *A*.

**Video S14**: Extract of a numerical integration of Eqs. 2 and 4 in 3D with 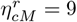 showing the target-wave pattern in the oscillatory regime (*d*_1_ = 1.35). The color code indicates the activator *A*.

## References

[1] C. Huang, F. Ling, and E. Kanso, Proceedings of the National Academy of Sciences 121, e2406293121 (2024).

[2] C. M. Topaz and A. L. Bertozzi, SIAM Journal on Applied Mathematics 65, 152 (2004).

[3] W.-J. Rappel, A. Nicol, A. Sarkissian, H. Levine, and W. F. Loomis, Physical review letters 83, 1247 (1999).

[4] T. H. Tan, A. Mietke, J. Li, Y. Chen, H. Higinbotham, P. J. Foster, S. Gokhale, J. Dunkel, and N. Fakhri, Nature 607, 287 (2022).

[5] T. H. Tan, A. Amiri, I. Seijo-Barandiarán, M. F. Staddon, A. Materne, S. Tomas, C. Duclut, M. Popović, A. Grapin-Botton, and F. Jülicher, PRX Life 2, 033006 (2024).

[6] J. L. Silverberg, M. Bierbaum, J. P. Sethna, and I. Cohen, Physical review letters 110, 228701 (2013).

[7] K. Tanner, H. Mori, R. Mroue, A. Bruni-Cardoso, and M. J. Bissell, Proceedings of the National Academy of Sciences 109, 1973 (2012).

[8] P. A. Fernández, B. Buchmann, A. Goychuk, L. K. Engelbrecht, M. K. Raich, C. H. Scheel, E. Frey, and A. R. Bausch, Nature physics 17, 1130 (2021).

[9] S. K. Ranamukhaarachchi, A. Walker, M.-H. Tang, W. D. Leineweber, S. Lam, W.-J. Rappel, and S. I. Fraley, Developmental Cell (2024).

[10] K. M. Yamada and M. Sixt, Nature Reviews molecular cell biology 20, 738 (2019).

[11] S. Huang, C. Brangwynne, K. Parker, and D. E. Ingber, Cell motility and the cytoskeleton 61, 201 (2005).

[12] B. A. Camley, Y. Zhang, Y. Zhao, B. Li, E. Ben-Jacob, H. Levine, and W.-J. Rappel, Proceedings of the National Academy of Sciences 111, 14770 (2014).

[13] F. J. Segerer, F. Thüroff, A. Piera Alberola, E. Frey, and J. O. Rädler, Physical Review Letters 114, 228102 (2015).

[14] A. M. Turing, Bulletin of mathematical biology 52, 153 (1990).

[15] A. Erzberger, A. Jacobo, A. Dasgupta, and A. Hudspeth, Nature physics 16, 949 (2020).

[16] S. Chen, D. S. Seara, A. Michaud, S. Kim, W. M. Bement, and M. P. Murrell, Nature Physics 20, 1824 (2024).

[17] J. Drgonova, T. Drgon, K. Tanaka, R. Kollar, G.-C. Chen, R. A. Ford, C. S. Chan, Y. Takai, and E. Cabib, Science 272, 277 (1996).

[18] D. B. Brückner, M. Schmitt, A. Fink, G. Ladurner, J. Flommersfeld, N. Arlt, E. Hannezo, J. O. Radler, and C. P. Broedersz, Physical Review X 12, 031041 (2022).

[19] R. Gorelik and A. Gautreau, Cytoskeleton 72, 362 (2015).

[20] Y. Cao, E. Ghabache, Y. Miao, C. Niman, H. Hakozaki, S. L. Reck-Peterson, P. N. Devreotes, and W.-J. Rappel, Journal of the Royal Society Interface 16, 20190619 (2019).

[21] P. Lu and Y. Lu, Frontiers in Cell and Developmental Biology 9, 704939 (2021).

[22] J. Allard and A. Mogilner, Current opinion in cell biology 25, 107 (2013).

[23] W. M. Bement, M. Leda, A. M. Moe, A. M. Kita, M. E. Larson, A. E. Golding, C. Pfeuti, K.-C. Su, A. L. Miller, A. B. Goryachev, et al., Nature cell biology 17, 1471 (2015).

[24] P. N. Devreotes, S. Bhattacharya, M. Edwards, P. A. Iglesias, T. Lampert, and Y. Miao, Annual review of cell and developmental biology 33, 103 (2017).

[25] Y. Miao, S. Bhattacharya, M. Edwards, H. Cai, T. Inoue, P. A. Iglesias, and P. N. Devreotes, Nature cell biology 19, 329 (2017).

[26] Y. Miao, S. Bhattacharya, T. Banerjee, B. Abubaker-Sharif, Y. Long, T. Inoue, P. A. Iglesias, and P. N. Devreotes, Molecular systems biology 15, e8585 (2019).

[27] S. Bhattacharya, T. Banerjee, Y. Miao, H. Zhan, P. N. Devreotes, and P. A. Iglesias, Science Advances 6, eaay7682 (2020).

[28] Y. Mori, A. Jilkine, and L. Edelstein-Keshet, Biophysical journal 94, 3684 (2008).

[29] N. Verschueren and A. Champneys, SIAM Journal on Applied Dynamical Systems 16, 1797 (2017).

[30] S. S. Lou, A. Diz-Muñoz, O. D. Weiner, D. A. Fletcher, and J. A. Theriot, Journal of Cell Biology 209, 275 (2015).

[31] F. Raynaud, M. E. Ambühl, C. Gabella, A. Bornert, I. F. Sbalzarini, J.-J. Meister, and A. B. Verkhovsky, Nature Physics 12, 367 (2016).

[32] H. Wang, S. Lacoche, L. Huang, B. Xue, and S. K. Muthuswamy, Proceedings of the National Academy of Sciences 110, 163 (2013).

[33] L. Lu, T. Guyomar, Q. Vagne, R. Berthoz, A. Torres-Sánchez, M. Lieb, C. Martin-Lemaitre, K. van Unen, A. Honigmann, O. Pertz, et al., Nature Physics, 1 (2024).

[34] D. Shao, W.-J. Rappel, and H. Levine, Physical review letters 105, 108104 (2010).

[35] Y. Cao, R. Karmakar, E. Ghabache, E. Gutierrez, Y. Zhao, A. Groisman, H. Levine, B. A. Camley, and W.-J. Rappel, Soft Matter 15, 2043 (2019).

[36] Y. Cao, E. Ghabache, and W.-J. Rappel, Elife 8, e48478 (2019).

[37] J. Löber, F. Ziebert, and I. S. Aranson, Scientific reports 5, 9172 (2015).

[38] See Supplemental Material at [URL] for details about the phase field model, activator-inhibitor dynamics, numerical methods, coarse-graining into 𝒮 space, statistics in 𝒮 space, semi-Markov renewal process, zero-dimensional back representation, Kramers escape time, experimental methods, robustness of the rotational rotational motion and weakly nonlinear analysis. The Supplemental Material cites Refs. [68–76].

[39] B. Lindner, J. Garcia-Ojalvo, A. Neiman, and L. Schimansky-Geier, Physics reports 392, 321 (2004).

[40] S. Alonso, M. Stange, and C. Beta, PloS one 13, e0201977 (2018).

[41] S. Monfared, A. Ardaševa, and A. Doostmohammadi, arXiv preprint arXiv:2503.05053 (2025).

[42] N. G. Van Kampen, Stochastic processes in physics and chemistry, Vol. 1 (Elsevier, 1992).

[43] O. Ibe, Markov processes for stochastic modeling (Newnes, 2013).

[44] H. Risken, in The Fokker-Planck equation: methods of solution and applications (Springer, 1989).

[45] H. A. Kramers, physica 7, 284 (1940).

[46] A. S. Chin, K. E. Worley, P. Ray, G. Kaur, J. Fan, and L. Q. Wan, Proceedings of the National Academy of Sciences 115, 12188 (2018).

[47] J. Hadidjojo and D. K. Lubensky, arXiv preprint arXiv:1708.08560 (2017).

[48] M. C. Cross and P. C. Hohenberg, Reviews of modern physics 65, 851 (1993).

[49] M. Cross and H. Greenside, Pattern formation and dynamics in nonequilibrium systems (Cambridge University Press, 2009).

[50] M. Ehrbar, A. Sala, P. Lienemann, A. Ranga, K. Mosiewicz, A. Bittermann, S. C. Rizzi, F. E. Weber, and M. P. Lutolf, Biophysical Journal 100, 284 (2011).

[51] Y. Abbas, A. Carnicer-Lombarte, L. Gardner, J. Thomas, J. J. Brosens, A. Moffett, A. M. Sharkey, K. Franze, G. J. Burton, and M. L. Oyen, Human Reproduction 34, 1999 (2019).

[52] M. H. Zaman, L. M. Trapani, A. L. Sieminski, D. MacKellar, H. Gong, R. D. Kamm, A. Wells, D. A. Lauffenburger, and P. Matsudaira, Proceedings of the National Academy of Sciences 103, 10889 (2006).

[53] J. Imran Alsous, N. Romeo, J. A. Jackson, F. M. Mason, J. Dunkel, and A. C. Martin, Proceedings of the National Academy of Sciences 118, e2019749118 (2021).

[54] O. Chaudhuri, L. Gu, D. Klumpers, M. Darnell, S. A. Bencherif, J. C. Weaver, N. Huebsch, H.-p. Lee, E. Lippens, G. N. Duda, et al., Nature materials 15, 326 (2016).

[55] R. M. Adar and J.-F. Joanny, Physical Review Letters 133, 118402 (2024).

[56] B. N. Narasimhan and S. I. Fraley, Proceedings of the National Academy of Sciences 122, e2416771122 (2025).

[57] M. P. Stewart, J. Helenius, Y. Toyoda, S. P. Ramanathan, D. J. Muller, and A. A. Hyman, Nature 469, 226 (2011).

[58] H. Jiang and S. X. Sun, Biophysical journal 105, 609 (2013).

[59] C. Roffay, G. Molinard, K. Kim, M. Urbanska, V. Andrade, V. Barbarasa, P. Nowak, V. Mercier, J. García-Calvo, S. Matile, et al., Proceedings of the National Academy of Sciences 118, e2103228118 (2021).

[60] M. Murrell, P. W. Oakes, M. Lenz, and M. L. Gardel, Nature reviews Molecular cell biology 16, 486 (2015).

[61] G. Badih, A. Schaeffer, B. Vianay, P. Smilovici, L. Blanchoin, M. Théry, and L. Kurzawa, Proceedings of the National Academy of Sciences 122, e2415028122 (2025).

[62] D. Vidmar and W.-J. Rappel, Physical Review E 99, 012407 (2019).

[63] D. Taniguchi, S. Ishihara, T. Oonuki, M. Honda-Kitahara, K. Kaneko, and S. Sawai, Proceedings of the National Academy of Sciences 110, 5016 (2013).

[64] Q. Yang, Y. Miao, L. J. Campanello, M. J. Hourwitz, B. Abubaker-Sharif, A. L. Bull, P. N. Devreotes, J. T. Fourkas, and W. Losert, Elife 11, e73198 (2022).

[65] Y. Yang and M. Wu, Philosophical Transactions of the Royal Society B: Biological Sciences 373, 20170116 (2018).

[66] C. Beta, L. Edelstein-Keshet, N. Gov, and A. Yochelis, Elife 12, e87181 (2023).

[67] S. Echeverría-Alar, B. N. Narasimhan, S. I. Fraley, and W.-J. Rappel, 10.5281/zenodo.18038143 (2025).

[68] A. Karma and W.-J. Rappel, Physical review E 57, 4323 (1998).

[69] T. Wakatsuki, R. B. Wysolmerski, and E. L. Elson, Journal of cell science 116, 1617 (2003).

[70] E. Perez Ipiña, J. d’Alessandro, B. Ladoux, and B. A. Camley, Proceedings of the National Academy of Sciences 121, e2318248121 (2024).

[71] M. S. Fabien, Spectral methods for partial differential equations that model shallow water wave phenomena, Ph.D. thesis (2014).

[72] A. Kaboudian, E. M. Cherry, and F. H. Fenton, Science advances 5, eaav6019 (2019).

[73] S. Echeverría-Alar and W.-J. Rappel, Phys. Rev. Lett. 135, 267201 (2025).

[74] J. P. Klein and M. L. Moeschberger, Survival Analysis: Techniques for Censored and Truncated Data, 2nd ed. (Springer, New York, 2003).

[75] J. B. Collins and H. Levine, Physical Review B 31, 6119 (1985).

[76] J. Landino, M. Leda, A. Michaud, Z. T. Swider, M. Prom, C. M. Field, W. M. Bement, A. G. Vecchiarelli, A. B. Goryachev, and A. L. Miller, Current Biology 31, 5613 (2021).

## References

[1] D. Shao, W.-J. Rappel, and H. Levine, Physical review letters 105, 108104 (2010).

[2] Y. Cao, E. Ghabache, and W.-J. Rappel, Elife 8, e48478 (2019).

[3] Y. Cao, E. Ghabache, Y. Miao, C. Niman, H. Hakozaki, S. L. Reck-Peterson, P. N. Devreotes, and W.-J. Rappel, Journal of the Royal Society Interface 16, 20190619 (2019).

[4] B. A. Camley, Y. Zhang, Y. Zhao, B. Li, E. Ben-Jacob, H. Levine, and W.-J. Rappel, Proceedings of the National Academy of Sciences 111, 14770 (2014).

[5] Y. Miao, S. Bhattacharya, M. Edwards, H. Cai, T. Inoue, P. A. Iglesias, and P. N. Devreotes, Nature cell biology 19, 329 (2017).

[6] A. Karma and W.-J. Rappel, Physical review E 57, 4323 (1998).

[7] T. Wakatsuki, R. B. Wysolmerski, and E. L. Elson, Journal of cell science 116, 1617 (2003).

[8] E. Perez Ipiña, J. d’Alessandro, B. Ladoux, and B. A. Camley, Proceedings of the National Academy of Sciences 121, e2318248121 (2024).

[9] M. S. Fabien, Spectral methods for partial differential equations that model shallow water wave phenomena, Ph.D. thesis (2014).

[10] A. Kaboudian, E. M. Cherry, and F. H. Fenton, Science advances 5, eaav6019 (2019).

[11] S. Echeverría-Alar, B. N. Narasimhan, S. I. Fraley, and W.-J. Rappel, 10.5281/zenodo.18038143 (2025).

[12] S. Echeverría-Alar and W.-J. Rappel, Phys. Rev. Lett. 135, 267201 (2025).

[13] J. P. Klein and M. L. Moeschberger, Survival Analysis: Techniques for Censored and Truncated Data, 2nd ed. (Springer, New York, 2003).

[14] O. Ibe, Markov processes for stochastic modeling (Newnes, 2013).

[15] J. B. Collins and H. Levine, Physical Review B 31, 6119 (1985).

[16] H. Risken, in The Fokker-Planck equation: methods of solution and applications (Springer, 1989) pp. 63–95.

[17] Y. Yang and M. Wu, Philosophical Transactions of the Royal Society B: Biological Sciences 373, 20170116 (2018).

[18] J. Landino, M. Leda, A. Michaud, Z. T. Swider, M. Prom, C. M. Field, W. M. Bement, A. G. Vecchiarelli, A. B. Goryachev, and A. L. Miller, Current Biology 31, 5613 (2021).

[19] C. Beta, L. Edelstein-Keshet, N. Gov, and A. Yochelis, Elife 12, e87181 (2023).

