## Supplemental Material for "Single-cell chiral symmetry breaking under confinement"

Sebastián Echeverría-Alar[1], Badri Narayanan Narasimhan[2], Stephanie I Fraley[2], and Wouter-Jan Rappel[1]
[1] *Department of Physics, University of California, San Diego, California 92093, USA and*
[2] *Department of Bioengineering, University of California, San Diego, California 92093, USA*

**Contents**

I. Phase field model 2

II. Numerical methods 3

III. Numerical implementation of the 2D phase field model in WebGL 5

IV. Supplemental figures for Fig. 1: Figs. S2, S3, S4 and S5 5

V. Coarse-graining into $\mathcal{S}$-space 5

VI. Statistics in $\mathcal{S}$-space 7

VII. Semi-Markov renewal process 10

VIII. Supplemental figures for Fig. 3: Fig.S15 and Fig.S16 10

IX. Zero-dimensional representation of the back of the cell 11

X. Quasi-steady approximation and Kramers escape 13

XI. Experimental setup 17

XII. Equivalence between weak confinement and extra free space 17

XIII. The oscillatory regime of the activator-inhibitor dynamics 18

XIV. Sensitivity analysis of the local activator-inhibitor dynamics 18

XV. Supplemental figures for Section III.D of the main text: Fig. S20 and Fig. S21 20

XVI. Video captions 20

### I. Phase field model

We model confined single-cell dynamics using a phase field approach. Let us consider a scalar field $\varphi_c(\mathbf{r},t)$ where $\mathbf{r}=(x,y,z)$, which provides a smooth and continuous representation of the cell position. Specifically, $\varphi_c=1$ inside the cell and $\varphi_c=0$ outside the cell, with the two states connected through an interface of width $\epsilon$ (Fig. S1A). The spatiotemporal evolution of the cell interface is governed by an advection equation, $\partial_t\varphi_c=-\mathbf{v}\cdot\vec{\nabla}\varphi_c$, where $\mathbf{v}$ is the velocity of the cell membrane in the outward normal direction, $\hat{\mathbf{n}}=-\nabla\varphi_c/|\nabla\varphi_c|$. We assume that cell motion occurs in a highly viscous extracellular matrix (ECM); thus, the friction at the cell-ECM interface can be modeled as a simple linear drag, $\mathbf{F}_f=-\xi\varphi_M\mathbf{v}$, where $\xi$ is the friction coefficient and $\varphi_M$ is the position of the ECM (see below). Then, a force balance at the cell membrane determines the velocity (Fig. S1B):

$$-\xi\mathbf{v}+\mathbf{F}_v+\mathbf{F}_t+\mathbf{F}_p+\mathbf{F}_r=0. \tag{1}$$

The second term in Eq. (1) represents an isotropic size-restoring force acting in the normal direction, which enforces a prescribed cell size $V_c$: $-\eta_v(\int\varphi_c d\mathbf{r}-V_c)\widehat{\mathbf{n}}$, where $\eta_v$ is a parameter controlling the strength of this force. The third term models the membrane tension of the cell. In equilibrium, this tension is associated with a Ginzburg-Landau type of energy $\mathcal{F}[\varphi_c,\nabla\varphi_c]=\gamma\int(\epsilon|\nabla\varphi_c|^2/2+G(\varphi_c)/\epsilon)d\mathbf{r}$, and the corresponding force can be expressed as $\mathbf{F}_t=(\delta\mathcal{F}[\varphi_c,\nabla\varphi_c]/\delta\varphi_c)\nabla\varphi_c/\tilde{\epsilon}$ [1], where $\tilde{\epsilon}=\epsilon|\nabla\varphi_c|^2$ and $\gamma$ is the surface tension coefficient. The function $G(\varphi_c)=18\varphi_c^2(1-\varphi_c)^2$ is a double well potential with minima at $\varphi_c=0$ and $\varphi_c=1$.

The fourth term is a non-equilibrium force in the outward normal direction that drives cell motion through protrusions, governed by activator-inhibitor ($A$-$R$) dynamics: $\mathbf{F}_p=\eta_p f(A)\widehat{\mathbf{n}}$. The parameter $\eta_p$ is chosen such that, in the absence of the ECM, the cell exhibits a persistent unidirectional motion, which can be interpreted as migration on a substrate [2, 3]. The activator $A$ represents a mean-field description of a signaling protein that models cell polarization by promoting actin polymerization. Following previous studies, we adopt the simple linear case $f(A)=A$ [1, 4]. However, we have verified that the more complex Hill-type function $f(A)=A^3/(A^3+A_o^3)$, used in Ref. [2], can also generate single-cell rotation. The activator-inhibitor dynamics is restricted to the cell interior through a function $\Omega_c(\varphi_c)$, which implicitly imposes non-flux boundary conditions, and is described by the coupled reaction-diffusion equation:

$$\begin{aligned}\partial_t(A\Omega_c)&=\Omega_c\left(ba-d_1A+k_a\frac{A^2}{K_a^2+A^2}a-d_2AR+\sqrt{\Gamma}\gamma_1(\mathbf{r},t)\right)+\nabla(D_A\Omega_c\nabla A)\\ \partial_t(R\Omega_c)&=\Omega_c\left(\frac{c_2A-c_1R}{\tau}\right)+\nabla(D_R\Omega_c\nabla R).\end{aligned} \tag{2}$$

This equation is equivalent to Eq. (1) of the main text with

$$\begin{aligned}\mathcal{A}(A,R)&=ba-d_1A+k_a\frac{A^2}{K_a^2+A^2}a-d_2AR\\ \mathcal{R}(A,R)&=\frac{c_2A-c_1R}{\tau}.\end{aligned} \tag{3}$$

The reaction terms have a similar functional form to those used in previous studies [5], and Eq. (2) has been used in previous models of cell plasticity in both two and three dimensions [2, 3]. The term $\gamma_1(\mathbf{r},t)$ denotes delta-correlated Gaussian noise in space and time, with $\langle\gamma_1(\mathbf{r},t)\rangle=0$ and $\langle\gamma_1(\mathbf{r},t)\rangle\langle\gamma_1(\mathbf{r}',t')\rangle=\delta(\mathbf{r}-\mathbf{r}')\delta(t-t')$, modeling the inherent fluctuations of microscopic protein dynamics. We have verified that including noise in both the activator and inhibitor equations does not affect our results in a qualitative way. The parameters in Eq. (2) are chosen such that the activator-inhibitor system exhibits excitable dynamics (Table S1 and Fig. S1C) and, as discussed above, induces unidirectional migration in the absence of confinement. Up to this point, the modeling framework has considered a three-dimensional cell (Fig. S1A). In this case, $\Omega_c(\varphi_c)=2G(\varphi_c)/\epsilon$, restricting the activator-inhibitor evolution near the cell surface to minimize numerical costs. Nevertheless, we also employ the same approach to study cell dynamics in a two-dimensional representation, where we define $\Omega_c(\varphi_c)=\varphi_c(x,y,t)$. Choosing the later for the 3D case does not affect qualitatively our results. Importantly, irrespective of the smooth function $\Omega_c$ used to indicate the cell position in space, the true free boundary problem for the activator-inhibitor system in a moving domain is recovered in the sharp interface limit ($\epsilon\to0$) [6].

The last term in Eq. (1) accounts for the confining effect of the ECM. To model this interaction, we introduce a second phase field, $\varphi_M(\mathbf{r})$, which specifies the position of ECM (Fig. S1A). The ECM is static, and its geometry is generated by carving a spherical (circular) hole in a cube (square) of size $L^3$ ($L^2$), and subsequently allowing the interface to relax diffusely until a prescribed volume $V_M$ is reached (see Section II). The force exerted by the ECM is given by $\mathbf{F}_r=-\eta_{cM}^r\varphi_M\widehat{\mathbf{n}}$, where $\eta_{cM}^r$ is the parameter controlling the confinement strength. We have further verified that modeling confinement as a repulsive potential interaction between the cell and the ECM, using

$\mathcal{F}^{rep}_{ECM}[\varphi_c, \varphi_M] = \eta^r_{cM} \int \varphi_c^2 \varphi_M^2 d\mathbf{r}$, does not alter our conclusions. Finally, substituting the velocity from Eq. (1) into the advection equation for $\varphi_c$ yields:

$$\xi \partial_t \varphi_c = \gamma \left( \nabla^2 \varphi_c - \frac{1}{\epsilon^2} \frac{\delta \mathcal{G}[\varphi_c]}{\delta \varphi_c} \right) - \eta_v \left( \int \varphi_c d\vec{r} - V_c \right) |\nabla \varphi_c| + \eta_p A |\nabla \varphi_c| - \eta^r_{cM} \varphi_M |\nabla \varphi_c|, \tag{4}$$

which is valid only when the cell is in contact with the ECM. Note that Eq. (4) implicitly incorporates cell contraction. For example, immediately after a protrusion locally extends the interface of a cell at equilibrium, the cell size increases: $\int \varphi_c d\vec{r} \approx V_c + |\delta V|$. This results in a contraction away from the protrusion with a strength proportional to the size increment $|\delta V|$. We have verified that explicit incorporation of local cell contraction, for example driven by myosin activity [7], as implemented in previous studies [4, 8], does not qualitatively alter our results. All parameters in Eq. (4) are summarized in Table S1.

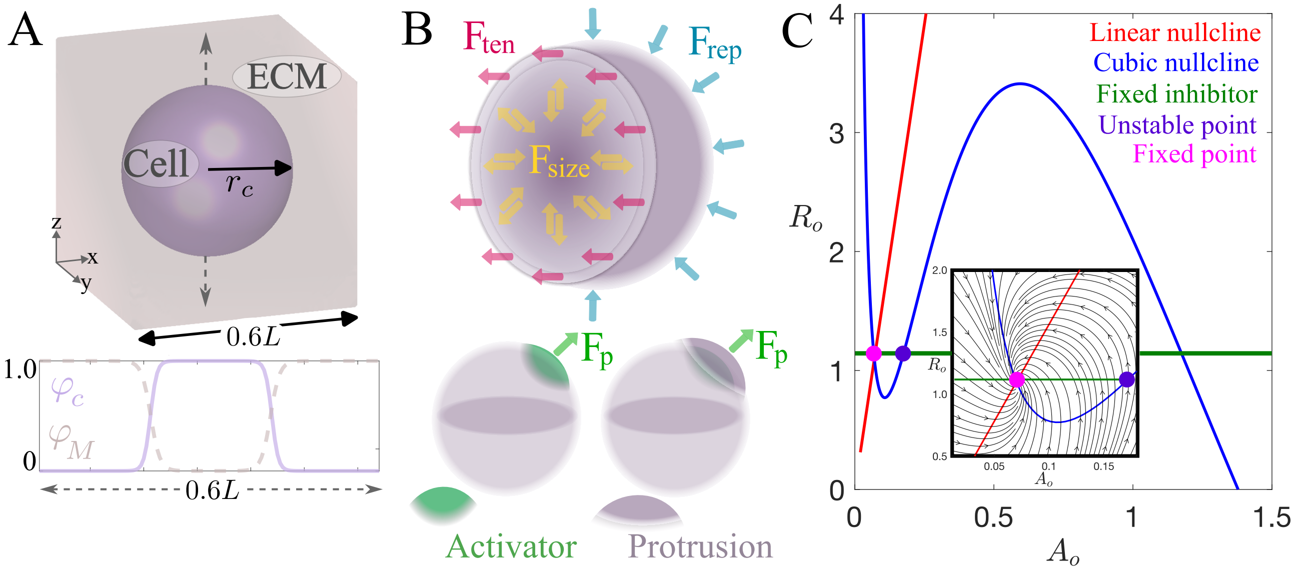


**Figure S1:** Model set-up. (A) Top panel: Initial three dimensional phase fields. For the cell, the phase field is shown at $\varphi_c = 1/2$, while the ECM phase field is shown for $\varphi_M > 1/2$. Bottom panel: One-dimensional cuts of $\varphi_c$ and $\varphi_M$ along the dashed line in the top panel. $r_c$ denotes the initial radius of the spherical cell. (B) Schematic representation of the forces acting on the cell surface, excluding ECM-cell friction. (C) Phase space of the zero-dimensional version of the activator-inhibitor dynamics in Eq. (2), given by $\{A_o, R_o\}$, in the excitable regime. The inset shows the structure of the phase flows around the fixed point (attractor). The unstable point, crossing between the fixed inhibitor and the cubic nullcline, is introduced further in this Supplemental Material.

### II. Numerical methods

The initial condition for the ECM, in both 2D and 3D simulations, is given by the steady state of an equation similar to Eq. (4), but dimensionless:

$$\partial_t \varphi_M = a_1 \left( \nabla^2 \varphi_M - \frac{1}{2} \frac{\delta \mathcal{F}[\varphi_M]}{\delta \varphi_M} \right) - a_2 \left( \int \varphi_M d\vec{r} - V_M \right) |\nabla \varphi_M|, \tag{5}$$

where $a_1 = 2/150$ and $a_2 = 100/150$ sets the strength of the diffusive process and the strength of prescribing the ECM size, respectively. This reaction-diffusion equation is discretized using finite differences with anisotropic stencils in the $\{x, y, z\}$ directions in 3D ($\{x, y\}$ directions in 2D) with non-flux boundary conditions and integrated in time with a forward Euler scheme. The integral is discretized using the trapezoidal method. The initial condition for the cell, $\varphi_c$ , is designed to emulate the experimental scenario of a cell seeded into an ECM. Particularly, the phase field $\varphi_c$ is initiated in the cavity of $\varphi_M$ as $1 - \varphi_M$, and is evolved for a short period with $\eta_v = \eta_p = 0$ and $\eta^r_{cM} = 52$ until reaching an equilibrium size determined by the balance between curvature and confinement (Fig. S1A). This equilibrium size is then used as the prescribed cell size $V_c$.

The three-dimensional phase field simulations were implemented in Matlab using GPU acceleration. The equation for $\varphi_c$ is solved via a pseudospectral integrating factor method with a RK4 time-stepping [9]. In this numerical scheme, the linear part is directly approximated in Fourier space, while the nonlinear terms are discretized using fast Fourier transforms (FFT) and transformed back to real space via inverse FFT. We use periodic boundary conditions for the phase field equations. Once $\varphi_c$ is obtained at times $t$ and $t+\Delta t$, the activator and inhibitor equations are solved using finite differences with anisotropic stencils (two neighbors per spatial direction). For example, the weighted diffusive term in the x-direction (for fixed $y$ and $z$) is discretized as

$$\partial_x(\Omega_c\partial_x A)|_{y,z} = \frac{1}{2\Delta x^2}[\Omega_c(x+\Delta x)+\Omega_c(x)][A(x+\Delta x)-A(x)] - \frac{1}{2\Delta x^2}[\Omega_c(x)+\Omega_c(x-\Delta x)][A(x)-A(x-\Delta x)]. \quad (6)$$

The corresponding terms in $y$ and $z$ directions of $A$ as well as the diffusive terms for $R$ are computed analogously. The weighting by the phase field automatically introduces non-flux boundary conditions. The activator and inhibitor are marched in time only inside the cell, $\Omega_c > \varphi_c^{th}$, using a forward Euler scheme. For example, the activator is updated as (similarly for the inhibitor):

$$\partial_t(\Omega_c A)|_{x,y,z} = \frac{1}{\Delta t}\left\{[\Omega_c(t+\Delta t)-\Omega_c(t)]\,A(t) + [A(t+\Delta t)-A(t)\Omega_c(t)]\right\}. \quad (7)$$

The noise terms $\gamma_1(\mathbf{r},t)$ and $\gamma_2(\mathbf{r},t)$ are generated using a Box-Mueller transform applied to two uniform random samples, and their discretization follows the Ito representation.

The two-dimensional simulations are performed either using WebGL, with custom Java and GLSL codes based on the *Abubu.js* library [10] (see Section III), or using Matlab. Numerically, the main difference from the 3D simulations is that the entire discretization of the phase field equation is carried out in real space using finite differences. Additional details can be found in the repository for this paper [11]. The numerical parameters are summarized in Table S1.

**Table S1:** Model and numerical parameters used for all the 2D and 3D computational simulations in this study. Note that $\Gamma$ and $\eta_v$ are fixed except in Figs. S11 and S16, respectively.

| Parameter | Description | Value |
|---|---|---|
| $\xi$ | Friction coefficient | 15 pN min $\mu$m$^{-2}$ |
| $\gamma$ | Membrane tension coefficient | 2 pN |
| $\epsilon$ | Width of the phase fields $\varphi_c$ and $\varphi_M$ | $\sqrt{2}$ $\mu$m |
| $\eta_v$ | Restoring-size strength in 2D (3D) simulations | 10 pN $\mu$m$^{-3}$ (6 pN $\mu$m$^{-4}$) |
| $\eta_p$ | Protrusion strength | 7 pN $\mu$m$^{-1}$ |
| $\eta_{cM}^r$ | Confinement strength | $[2-52]$ pN $\mu$m$^{-1}$ |
| $V_c$ | Cell volume (area) | 376 $\mu$m$^3$ (83 $\mu$m$^2$) |
| $V_M$ | ECM volume (area) | 15250 $\mu$m$^3$ (2370 $\mu$m$^2$) |
| $k_a$ | Rate of activation | 10 min$^{-1}$ |
| $K_a$ | Threshold of activation | 1 $\mu$M |
| $b$ | Rate of basal activation | 0.1 min$^{-1}$ |
| $a$ | Total activator concentration in the cell | 2 $\mu$M |
| $d_1$ | Basal degradation rate of the activator in 2D (3D) simulations | 3 (2.7) s$^{-1}$ |
| $d_2$ | Inhibitor-induced degradation of the activator | 1 $\mu$M$^{-1}$ min$^{-1}$ |
| $D_A$ | Diffusion coefficient of the activator in 2D (3D) simulations | 0.1 (0.08) $\mu$m$^2$ min$^{-1}$ |
| $\tau$ | Slow time scale of the inhibitor | 10 min |
| $c_2$ | Activation coefficient of the inhibitor | 15 |
| $c_1$ | Degradation coefficient of the inhibitor | 0.96 |
| $\Gamma$ | Amplitude of the noise in 2D (3D) simulations | 0.03$^2$ (0.09$^2$) $\mu$M$^2$ min$^{-2}$ |
| $D_R$ | Diffusion coefficient of the inhibitor in 2D (3D) simulations | 0.2 (0.4) $\mu$m$^2$ min$^{-1}$ |
| $\Delta t$ | Time step in 2D (3D) simulations | 0.004 (0.006) min |
| $L$ | Domain size in 2D (3D) simulations | 50 (25) $\mu$m |
| $N$ | Number of discretization points per direction in 2D (3D) simulations | 256 (128) |
| $\varphi_c^{th}$ | Phase field threshold | 0.0025 |

#### III. Numerical implementation of the 2D phase field model in WebGL

We have developed a GPU-accelerated WebGL platform to solve the two-dimensional forms of Eqs (2) and (4) [11], using a specialized library for integrating partial differential equations via fragment shaders [10]. The simulations run directly in a web browser (e.g., Google Chrome, Safari, Firefox), enabling real-time interaction with the dynamics. For example, a click on top of the cell can perturb its activator field $A$, allowing the manual generation of protrusions. Additionally, the code is capable of storing data—simply by taking snapshots of the canvas on the webpage—while bypassing the usual GPU-CPU memory-transfer bottleneck [12]. We have leveraged this storage approach to efficiently explore multiple parameters in parallel (see [11] for details).

#### IV. Supplemental figures for Fig. 1: Figs. S2, S3, S4 and S5

The activator-inhibitor dynamics in the weak and intermediate regimes that lead to rotational motion involve the break-up of target waves along the cell surface. This breakage can be triggered either by stochastic fluctuations (Fig. S2), or by heterogeneities in the refractoriness of the excitable system (Fig. S3), which are dictated by the presence of multiple waves interacting along the cell surface.

The rotational motion in the weak confinement regime ($\eta^r_{CM} \lesssim 5$) is characterized by a frustrated "figure of eight" activator pattern (Fig. S4), and by a rotational period $T$. This frustration arises because the characteristic wavelength of the excitable waves—imposed by the diffusion coefficients $D_A$ and $D_R$—is relatively large compared with the finite size of the cell. Another signature of the coherent motion is the increase in the amplitude of the centroid trajectory, which reflects protrusion enlargement as confinement weakens (Fig. S5).

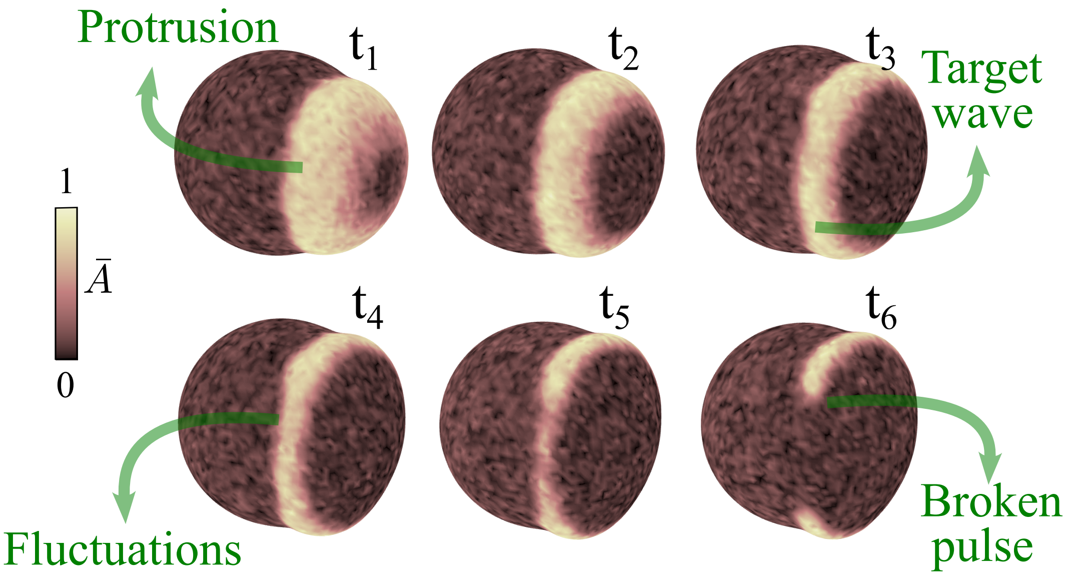


**Figure S2:** Temporal snapshots ($t_1 < t_2 < t_3 < t_4 < t_5 < t_6$) of the normalized activator field $\bar{A} = A/\max(A)$, illustrating the break-up of a target wave induced by stochastic fluctuations in the intermediate regime ($\eta^r_{cM} = 5.3$).

#### V. Coarse-graining into $\mathcal{S}$-space

To coarse-grain the 2D phase-field data into the discrete space $\mathcal{S}$, we begin by processing the numerical results in MATLAB, using output from 2D simulations of the phase-field and activator–inhibitor dynamics using WebGL. We use the activator field $A$ to detect pulses (case $\eta^r_{cM} = 6.5$ in Fig. S6A). First, the raw RGB image of the activator is smoothed using a Gaussian blur with a $9 \times 9$ kernel to homogenize the noisy background. Next, a binary image (mask) is created to indicate where the activator exceeds 90% of its maximum value. To avoid holes in the mask, a dilation process with a $9 \times 9$ kernel is applied. Finally, we use the function *regionprops()* of Matlab to identify connected regions in the binary image and extract their centroids. The number of centroids serves as a proxy for the number of pulses.

Errors in the detection algorithm are expected due to noisy transient pulses that may become frequent in the large dataset under analysis. These transients exhibit a high value of $A$ and are large enough to produce a measurable centroid, but they persist only for a short time. Figure S6B shows a raw trajectory of the number of pulses, where many of such short episodes are detected. It is important to remove them because they bias the computation of transition probabilities and first passage times. To address this, we implement a custom code to filter out spurious events defined as having a duration shorter than $T/45$. Briefly, the code detects segments of the trajectory, e.g., 11110001111, and replaces the sandwiched zeros with ones if their duration is smaller than $T/45$. Notice that events containing four pulses are always removed because they last less than the cutoff. The origin of this configuration is

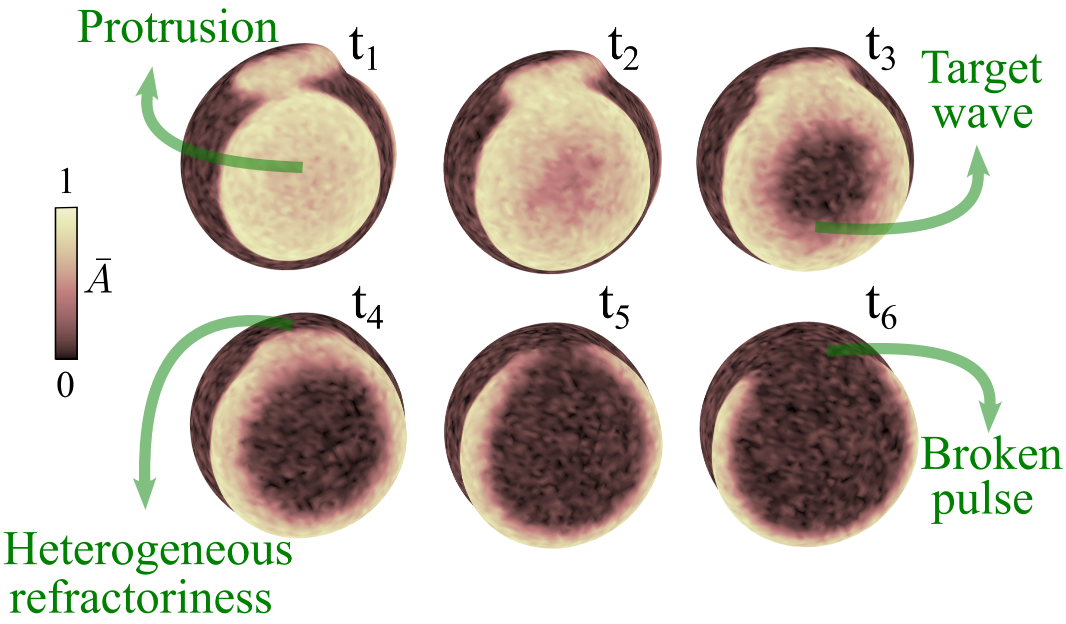


**Figure S3:** Temporal snapshots ($t_1 < t_2 < t_3 < t_4 < t_5 < t_6$) of the normalized activator field $\bar{A} = A/\max(A)$, illustrating the break-up of a target wave induced by heterogeneities in refractoriness in the intermediate regime ($\eta^r_{cM} = 5.3$).

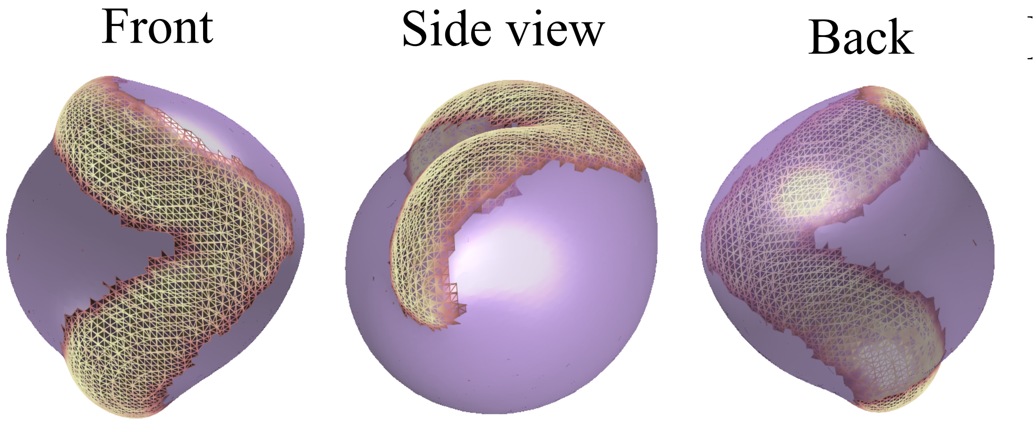


**Figure S4:** Coherent rotational wave. Different three-dimensional views of the activator spatial structure in the rotating state at $\eta^r_{cM} = 4$. Only the portion $A > 0.8$ of the activator field is shown.

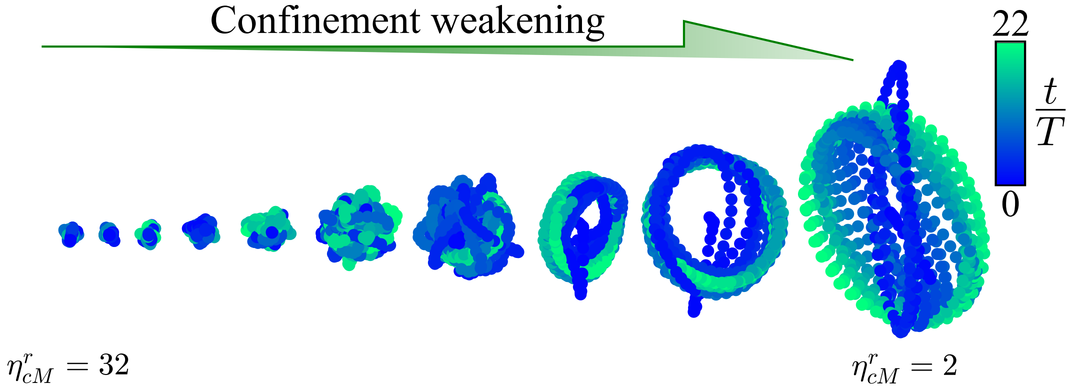


**Figure S5:** Three dimensional trajectories of the centroid of the cell for the range $\eta^r_{cM} = [2, 32]$ over $22T$. The trajectories start at the initial condition.

the fragmentation of two protrusions (Video S12), and their short duration is governed by the size-restoring force preventing the presence of more than three pulses. Figure S6C illustrates the temporal evolution of the number of pulses after applying the cutoff $T/45$.

Detecting the number of pulses alone is not sufficient to distinguish between the possible discrete states of the system, as discussed in the main text. Particularly, both a single chiral pulse and a non-chiral pulse yield the same pulse count of one, but their difference is fundamental. To tackle this, we include in our data analysis a routine to determine whether or not pulses rotate. For this, we compute the angle $\theta$ of the orbit of each activator pulse with respect to the cell center (rightmost panel; Figure S6A). Next, this angle is unwrapped and a linear model is fitted over the duration of the single pulse visit. For each visit, we calculate the coefficient

$$R^2 = 1 - \frac{\sum_i (\theta_i - \hat{\theta}_i)^2}{\sum_i (\theta_i - \langle\theta\rangle_{visit})^2}, \tag{8}$$

where $\hat{\theta}$ are the angle values predicted by the linear model, and $\langle\theta\rangle_{visit}$ is the average angle within the visit. If $R^2 > R^2_{th}$, the single pulse event is classified as a rotating pulse. Short duration events may produce false positives by flagging non-chiral pulses as chiral pulses. We correct these cases by considering the quasi-deterministic transitions on the discrete system. For example, the transition out of the resting state typically proceeds through the creation of a non-chiral pulse, which then divides into two chiral pulses (Video S7). If that one-pulse state, which has a short duration ($< 0.25T$), is flagged as a chiral pulse, we reclassify it as non-chiral.

The same procedure is applied to differentiate between chiral and non-chiral structures when the pulse count is equal to two, by defining two angles. However, when the pulse count is equal to three, the differentiation by angle is more cumbersome due to the short duration of almost all the episodes with three pulses. As mentioned in the main text, we distinguish only one of the four possible three-pulse configurations. This analysis leads to the coarse-grained state space $\mathcal{S} = \{\mathcal{S}_1, \mathcal{S}_2, \mathcal{S}_3, \mathcal{S}_4, \mathcal{S}_5, \mathcal{S}_6, \mathcal{S}_7, \mathcal{S}_8\}$ (Fig. S6D), where the chiral single pulse corresponds to $\mathcal{S}_3$, which is equivalent to $\mathcal{S}_{rot}$ (see the main text for the other definitions). More trajectories, for different confinement values, are shown in Fig. S7.

We chose $R^2_{th} = 0.9$ to define coherent rotations. While this value affects the classification quantitatively, the qualitative trends are preserved across different confinement regimes. As a consistency check, Fig. S8 shows how the average time spent at the rotating state $\mathcal{S}_3$ before jumping to another state (average dwell time; see next Section VI) depends on $R_{th}$ for different values of $\eta^r_{cM}$.

### VI. Statistics in $\mathcal{S}$-space

In the framework of the discrete dynamics in $\mathcal{S}$, we characterize the system in terms of dwell times $\tilde{\tau}_D$ in each $\mathcal{S}_i$ and the transition probabilities between states $P(\mathcal{S}_j|\mathcal{S}_i)$. Dwell times are recorded from 400 trajectories for each confinement parameter. In each trajectory, every visit to state $\mathcal{S}_i$ contributes to the dwell time count, even if the visit is right-censored. We record which dwell times are uncensored versus right-censored and store them as separate distributions (see Fig. S9). From this statistical information, the survival probability $Pr\{\tilde{\tau}_D > \tilde{t}\}$ can be obtained with the built-in function of Matlab *ecdf()*, which computes the probability distribution $Pr\{\tilde{\tau}_D \leq \tilde{t}\}$ including right-censored data via the Kaplan-Meier estimator (Fig S10). Moreover, mean dwell times $\mathbb{E}[\tilde{\tau}_D]$ per state can also be extracted from the distributions in Fig. S9. In order not to underestimate this averaged quantity, we approximate the expectation by integrating the corresponding survival probabilities [13]:

$$\mathbb{E}[\tilde{\tau}_D] = \int_0^{\tilde{t}_{max}} Pr\{\tilde{\tau}_D > \tilde{t}\} d\tilde{t}, \tag{9}$$

where $\tilde{t}_{max}$ is the largest time at which one can estimate $Pr\{\tilde{\tau}_D > \tilde{t}\}$. $\mathbb{E}[\tilde{\tau}_D]$ is a compact measurement of persistence in each state of the discrete system, which is particularly useful to study confinement effects on the maintenance of cell rotation (see Main text). In additional simulations, we have checked the influence of noise amplitude ($\Gamma$) on mean dwell times at $\mathcal{S}_{rot}$ (Fig. S11).

From the observed uncensored visits per state, the transition probabilities are approximated by

$$P(\mathcal{S}_j|\mathcal{S}_i) = \frac{\mathcal{N}_{i\to j}}{\sum_{k\neq i} \mathcal{N}_{i\to k}}, \tag{10}$$

where $\mathcal{N}_{i\to j}$ is the number of times a transition is observed from $\mathcal{S}_i$ to $\mathcal{S}_j$ and $\sum_{k\neq i} \mathcal{N}_{i\to k}$ is the total number of exits from $\mathcal{S}_i$. Transition probabilities smaller than 2% are set to zero, and the respective state probabilities are renormalized. The dependence of $P(\mathcal{S}_j|\mathcal{S}_i)$ on the confinement parameter is illustrated in Fig S12.

Additionally, we estimate first passage times $\tilde{\tau}(\mathcal{S}_i \to \mathcal{S}_j)$ from the rotating state ($\mathcal{S}_{rot}$) to all the other possible states (Fig. S13). The first passage time is numerically defined as the time spent in state $\mathcal{S}_i$ before transitioning

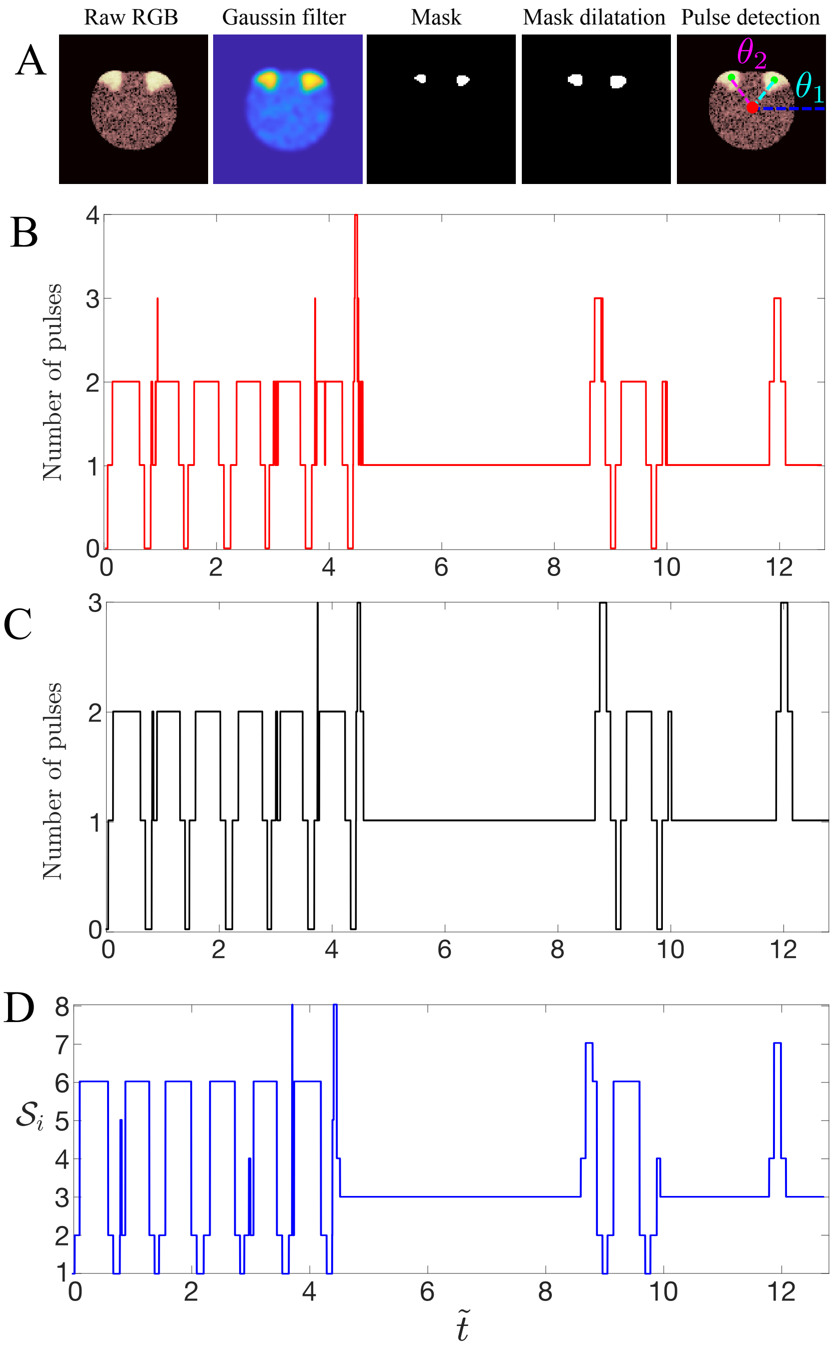


**Figure S6:** Detection of pulses. (A) Image process to extract the number of pulses (green circles) from the whole activator field. The red circle indicate the centroid of the cell. (B) Raw temporal trajectory of the number of pulses after image processing shown in (A). (C) Temporal trajectory of the number of pulses after removing spurious pulse counts. (D) Trajectory in $\mathcal{S}$-space. The confinement value for this particular trajectory is $\eta^r_{cM} = 6.5$. The tilde superscript represents normalization by the rotational period $T$ of the rotating state, measured at $\eta^r_{cM} = 3$.

for the first time to another state $\mathcal{S}_j$. Mean first passage times are calculated by building a survival probability for empirical first passage times $Pr\{\tilde{\tau}(\mathcal{S}_i \to \mathcal{S}_j) > \tilde{t}\}$, considering uncensored and censored (not reaching the target $\mathcal{S}_j$ over the observation window given the start $\mathcal{S}_i$) trips. Then, the expectation (mean) value is determined by the integration of this survival probability. In addition to the mean, we also report the $10^{\text{th}}$ and $90^{\text{th}}$ percentiles of the first-passage times. These are estimated from the probability distribution $Pr\{\tilde{\tau}(\mathcal{S}_i \to \mathcal{S}_j) \leq \tilde{t}\}$, as the smallest $\tilde{t}$ such

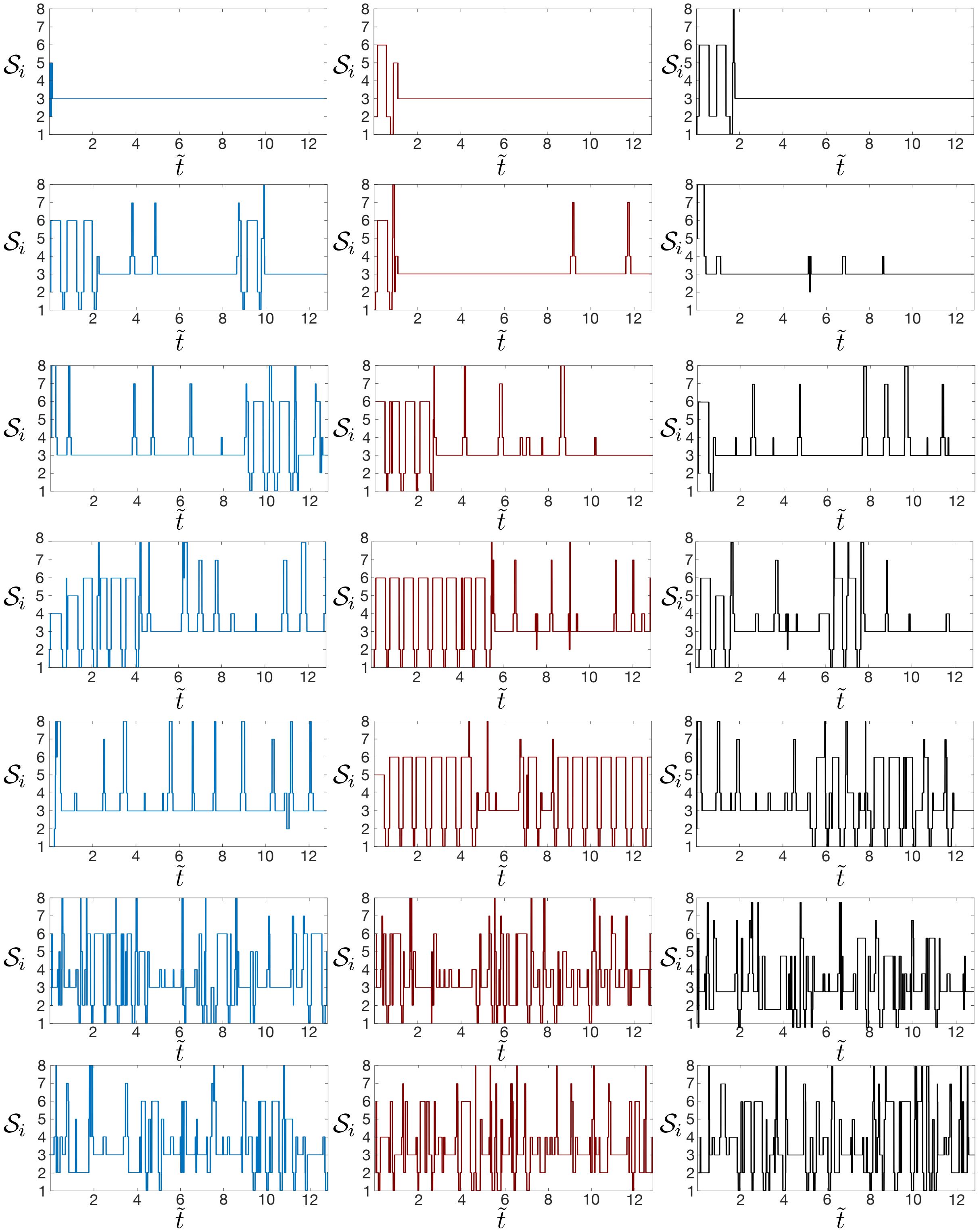


**Figure S7:** Three exemplary trajectories (columns) for each confinement $\eta^r_{cM}$ in the range $[2, 32]$. The tilde superscript represents normalization by the rotational period $T$ of the rotating state, measured at $\eta^r_{cM} = 3$

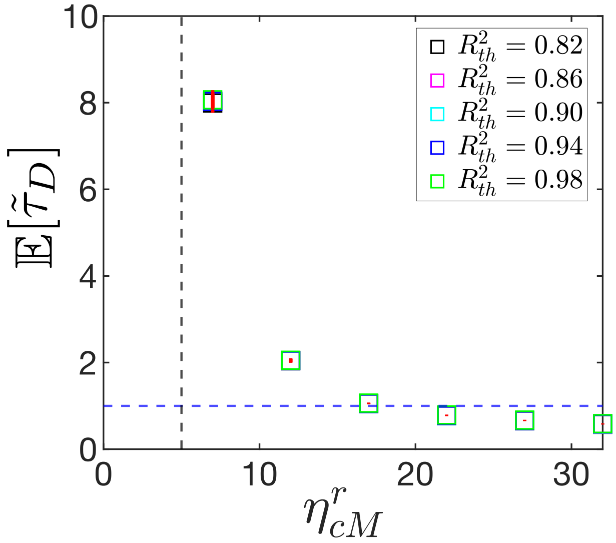


**Figure S8:** Effect of $R^2_{th}$ on the average dwell time at state $\mathcal{S}_3$. The tilde represents normalization by the rotational period $T$. The solid red lines indicate the bootstrap standard error (SE) of the average dwell time, estimated from 500 resamples. The maximum SE is less than $T/5$. The blue dashed line labels one rotational period and the black dashed line accounts for the critical confinement $\eta^r_{cM} \approx 5$.

that $Pr\{\tilde{\tau}(\mathcal{S}_i \to \mathcal{S}_j) \leq \tilde{t}\} \geq p$, with $p$ equal to 0.1 and 0.9, respectively. We do this to show the variability of the data, but also to stress the effects of heavy right-censoring, which sometimes forbids computing the 90$^{\text{th}}$ percentile. The latter is a signature that expectation values are being underestimated. We also estimate the median (50$^{\text{th}}$) to determine whether computing mean first passage times is meaningful.

### VII. Semi-Markov renewal process

Based on the dwell time statistics, we decide to introduce a semi-Markov renewal process as a data-driven model for our 2D system. We assume that an embedded Markov chain governs the transitions between states, i.e., the process is Markovian just before and after the transition occurs. However, the waiting times in each state before jumping to any other state are not exponentially distributed across confinements and across the whole observation time. Therefore, the whole stochastic dynamics is better represented by a semi-Markovian process. The renewal part comes from the assumption that upon arrival at a new state, the system resets and it is governed by the waiting time distributions and transition probabilities of that new state only. In this framework, one can write a simple linear equation for the mean first passage time between states $\mathcal{S}_i$ and $\mathcal{S}_j$, which compactly combines average dwell times and transition probabilities [14]:

$$\mathbb{E}[\tilde{\tau}(\mathcal{S}_i \to \mathcal{S}_j)] = \mathbb{E}[\tilde{\tau}_D]_i + \sum_{k \neq j} P(\mathcal{S}_k|\mathcal{S}_i)\, \mathbb{E}[\tilde{\tau}(\mathcal{S}_k \to \mathcal{S}_j)], \tag{11}$$

where the first passage times are $\tilde{\tau}(\mathcal{S}_i \to \mathcal{S}_j) = \min(\tilde{t} > 0 : \mathcal{S}(\tilde{t}) = \mathcal{S}_j|\mathcal{S}(0) = \mathcal{S}_i)$. The first term accounts for the average waiting time in state $\mathcal{S}_i$, while the second term considers all the possible transient trips $\mathcal{S}_i \to \mathcal{S}_k$ before absorption at $\mathcal{S}_j$. The linear system (11) can be solved by fixing the target state $\mathcal{S}_j$, extracting the matrix $\mathcal{Q} = \sum_{k \neq j} P(\mathcal{S}_k|\mathcal{S}_i)$, and inverting the matrix $I - \mathcal{Q}$, where $I$ is the identity matrix.

To test the predictive power of the renewal model, we compare the empirical and modeled mean first passage times. The semi-Markov renewal description reproduces fairly well $\mathbb{E}[\tilde{\tau}(\mathcal{S}_{rot} \to \mathcal{S}_j)]$ sufficiently far from $\eta^r_{cM} \approx 5$ (Fig. S14). The differences are less than two rotational periods except at $\eta^r_{cM} = 7$. The significant mismatch at $\eta^r_{cM} = 7$ comes from a large number of censored trajectories at $\mathcal{S}_{rot}$ (Fig. S13), which greatly underestimates the empirical mean first passage times. It also reflects that the renewal theory assigns a small probability of transitioning to $\mathcal{S}_1$, and a high probability of remaining in the loop $\mathcal{S}_{rot} \leftrightarrow \mathcal{S}_4$ (Fig. S12).

### VIII. Supplemental figures for Fig. 3: Fig.S15 and Fig.S16

Upon confinement weakening, the localized protrusion that drives the rotational motion becomes larger, and eventually ($\eta^r_{cM} \lesssim 5$), is able to translate the entire cell (Fig. S15). This morphological and dynamical change is associated with an enhancement of cell contraction far from the protrusion, which is sensible to variations in the strength of the size-restoring force (Fig. S16).

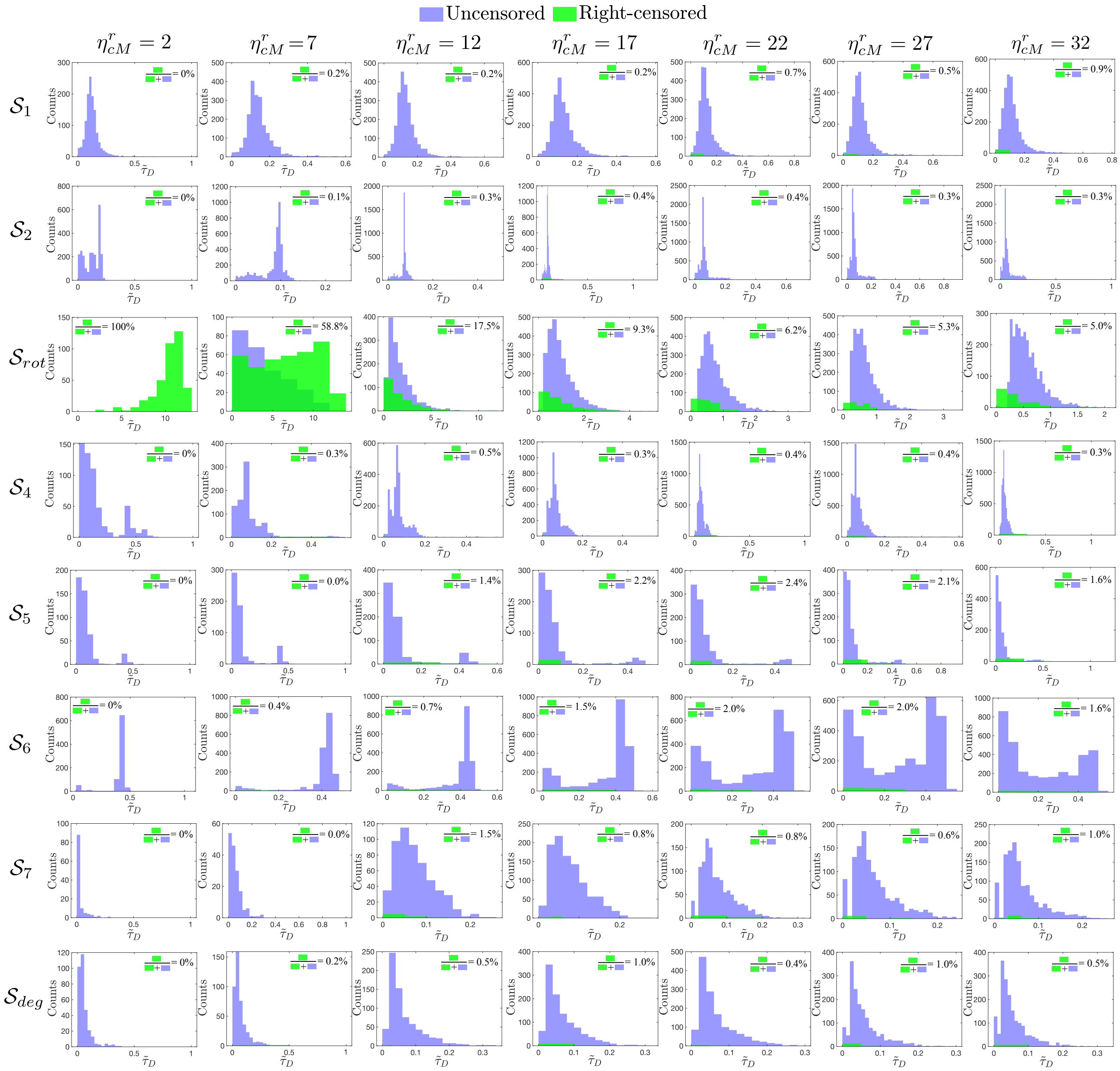


**Figure S9:** Distribution of dwell times $p(\tilde{\tau}_D)$ of the eight states of the discrete for different confinement values $\eta^r_{cM}$.

#### IX. Zero-dimensional representation of the back of the cell

In the main text, we investigate the role of the rotational protrusion in the dynamics of the rest of the cell. Particularly, due to trapping in the rotating state under weak confinement, a key question is how the excitable dynamics of the system is affected such that new activations (trips out of $\mathcal{S}_3$) are not observed over finite times. This question is complex to address at the whole cell level, so we adopt a simplified approach. Excitations cannot be triggered at the location of the first pulse, but only at a finite distance from it. Then, a convenient and simple region to analyze the effects of activation suppression is at the back of the cell; farthest from the protrusion.

Activation suppression in the excitable system arises from cell mechanics, as reducing confinement modifies the force balance in Eq. (1). At the back, the cell experiences compression triggered by membrane extension at the protrusion site. This mechanical effect feeds back into the activator-inhibitor dynamics via the phase-field coupling in Eq. (2). In our simplified approach, we collapse Eq. (2) to a single point at the back by fixing $\varphi_c = 1/2$, defining local back variables $\{A_b, R_b\}$ and neglecting spatial variations:

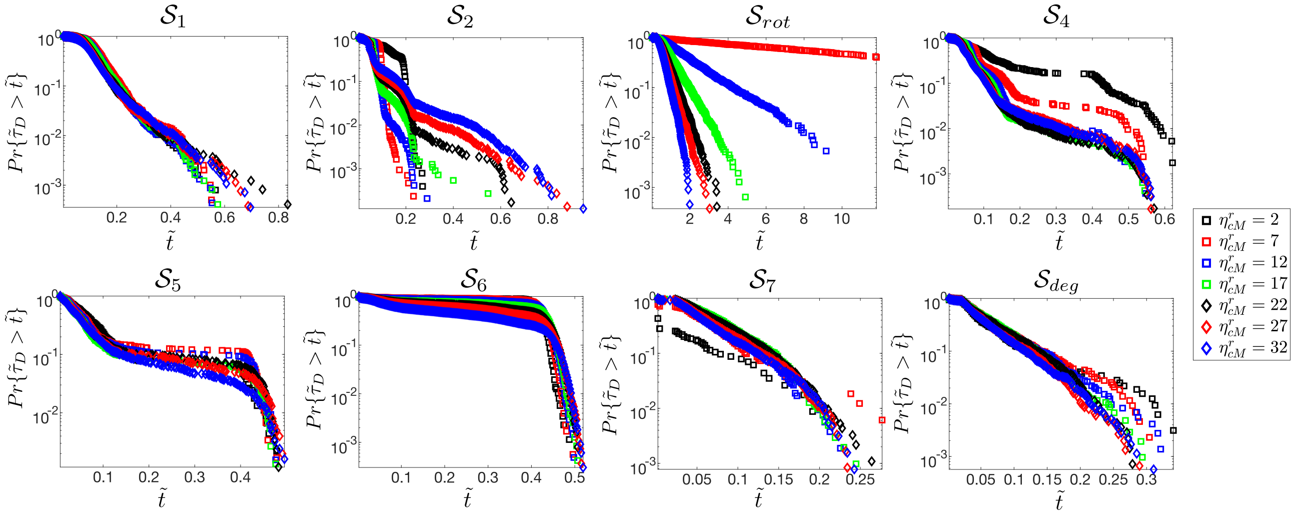


**Figure S10:** Survival curves $Pr(\tilde{\tau}_D > \tilde{t})$ of the eight states of the discrete system for different confinements.

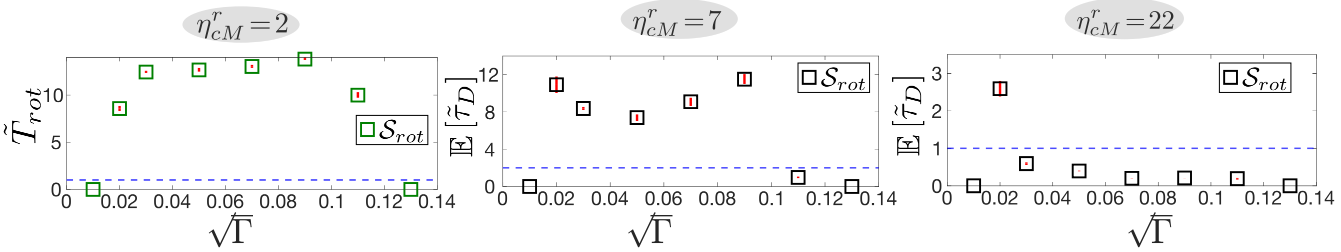


**Figure S11:** Characterization of noise amplitude variations in 2D simulations. The sweep of $\sqrt{\Gamma}$ is is in steps of 0.02. There is a minimum noise necessary to trigger transitions out of $\mathcal{S}_1$ ($\sqrt{\Gamma} = 0.02$) and a maximum amount of noise that does not allow the necessary coherence to create protrusions ($\sqrt{\Gamma} = 0.13$). $\tilde{T}_{rot}$ is the average time spent in the rotating state at $\eta^r_{cM} = 2$, where all the trajectories are right-censored. The large $\tilde{T}_{rot}$ is robust over a range of noise amplitudes. Interestingly, the behavior of $\mathbb{E}[\tilde{\tau}_D]$ at $\eta^r_{cM} = 7$ as a function of $\sqrt{\Gamma}$ is bimodal. The second peak for relatively high noises is correlated with a high chance of reaching $\mathcal{S}_{rot}$ from $\mathcal{S}_1$, but a low chance of going out. All the results in the main text are reported for $\sqrt{\Gamma} = 0.03$.

$$\begin{aligned}
\partial_t A_b &\approx ba - d_1 A_b + k_a \frac{A_b^2}{K_a^2 + A_b^2} a - d_2 A_b R_b + \sqrt{\Gamma}\gamma_1(\mathbf{r}, t)\delta(\mathbf{r} - \mathbf{r}_b) - 2\partial_t \varphi_b A_b \\
\partial_t R_b &\approx \left( \frac{c_2 A_b - c_1 R_b}{\tau} \right) - 2\partial_t \varphi_b R_b.
\end{aligned} \tag{12}$$

The corrections arising from the phase field coupling, proportional to $\partial_t \varphi_b / \varphi_c = 2\partial_t \varphi_b$, incorporate boundary deformation effects on the excitable dynamics in the form of back contractions: $\partial_t \varphi_b \approx v_n |\partial_n \varphi_b|$ (using Eq. (2) of the main text), where the subscript $n$ indicates the normal direction at the interface and $v_n < 0$. Additionally, the term $\partial_t \varphi_b$ serves as a proxy for the advection correction localized at the boundary in the sharp interface limit [15]. The reported value of $\partial_t \phi_b$ is a spatiotemporal average over $2T$ rotations and over an arc—centered at the interface point farthest from max$(A)$ and spanning approximately one quarter of the cell perimeter ($\sqrt{\pi V_c}/2$)—obtained from 2D numerical integrations of Eqs. (2) and (4) in the noiseless case ($\Gamma = 0$), using a stable rotating state as the initial condition (Fig. 3; main text). The modification of the underlying excitable activator-inhibitor dynamics can be quantified by computing the corrected fixed point of Eq. (12), and a comparison between the corrected and numerically measured fixed points at the back ($\{A_b^s, R_b^s\}$) reveals the same decreasing trend with decreasing confinement (Fig. S17A). In phase space, the change upon confinement (or contraction) can be simply characterized by the distance $\mathcal{D}$ from the fixed point to the minimum of the cubic nullcline (Fig. S17B). The trend in $\mathcal{D}$ as a function of $\eta^r_{cM}$ indicates that, when the cell is less confined (and more contracted), the activator-inhibitor dynamics becomes less excitable.

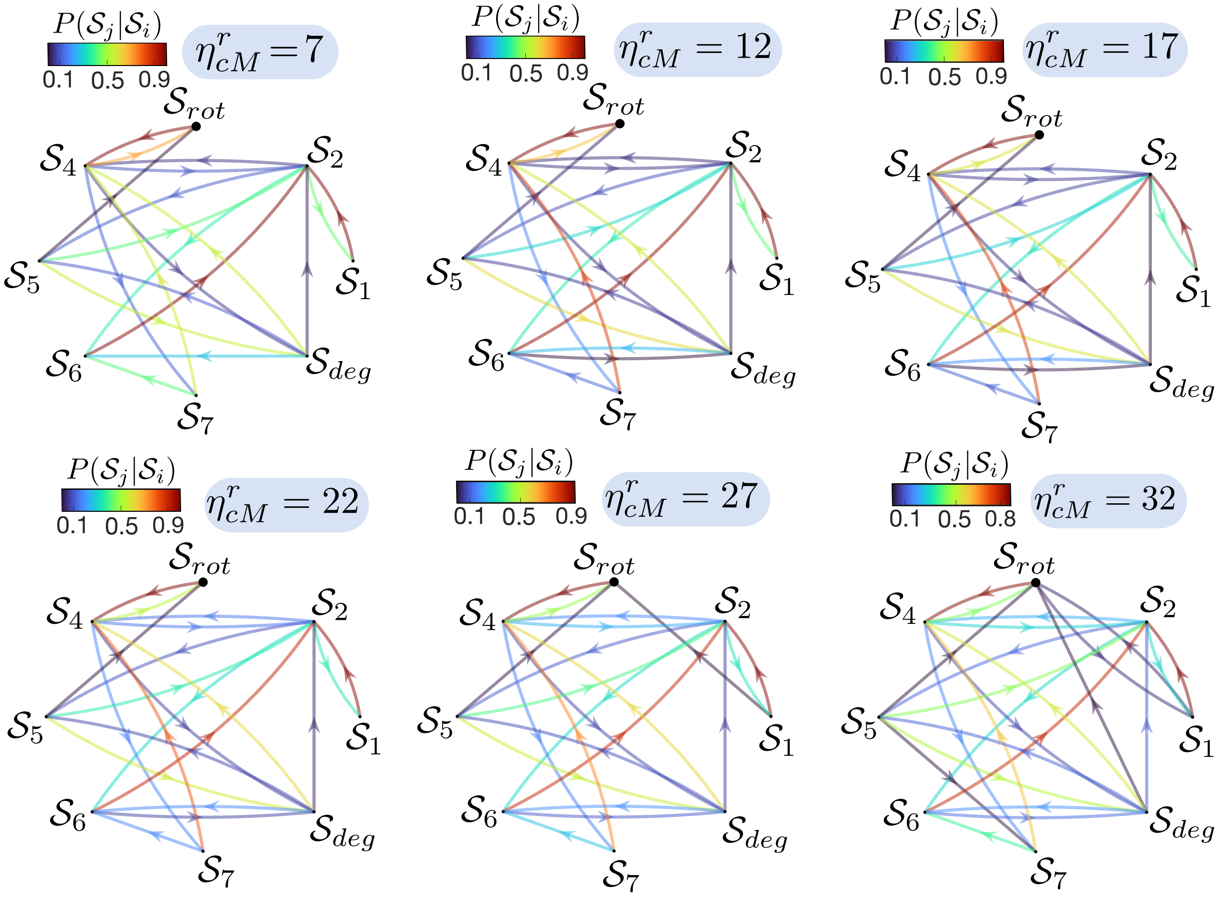


**Figure S12:** Transition probabilities in the discrete space $\mathcal{S}$ for different confinement values $\eta^r_{cM}$.

### X. Quasi-steady approximation and Kramers escape

To connect the statistical analysis performed in this study with the simplified back of the cell approach, we assume a quasi-steady approximation for the activator-inhibitor dynamics and freeze the slow inhibitor dynamics in time, setting $R_b = R_b^s$, i.e., $\tau \gg 1$ in the dimensionless version of Eq. (12). The mechanical correction in the dynamics of the back inhibitor does not introduce additional time scales ($c_2 \sim \tau|\partial_t\varphi_b|$). Under this approximation, the back activator-inhibitor dynamics can be recast into a compact Langevin equation in the Ito representation

$$\begin{aligned} dw &= \left(ba - d_1 w + k_a \frac{w^2}{K_a^2 + w^2} a - d_2 w R_b^s - 2\partial_t\varphi_b w\right) dt + \sqrt{\Gamma} dW_t \\ &= -\frac{d\mathcal{U}(w, \eta^r_{cM})}{dw} dt + \sqrt{\Gamma} dW_t, \end{aligned} \tag{13}$$

where $W_t$ is a Wiener process [16], and for simplicity $w = A_b$. Moreover, this Langevin equation is variational with potential:

$$\mathcal{U}(w, \eta^r_{cM}) = -abw + \frac{1}{2}\left(b + (d_1 + d_2 R_b^s) - 2|\partial_t\varphi_b|\right) w^2 - \frac{k_a}{2}\left(K_a^2 \log(K_a^2 + w^2) - 2aK_a \operatorname{atan}\left(\frac{w}{K_a}\right) + w(2a - w)\right), \tag{14}$$

where the confinement dependence comes via $\partial_t\varphi_b$. The problem is thus reduced to a particle in a potential well that may escape over the potential barrier due to stochastic fluctuations. This is the well-known Kramers escape problem, in which the escape rate $\mathcal{R}$ from the potential well—defined as the conditional probability per unit time that a particle escapes, given that it is in the potential well minima—can be computed [16]. For this, we introduce the corresponding Fokker-Planck equation for the probability density $P(w, t)$:

$$\partial_t P(w, t) = \partial_w[\partial_w \mathcal{U}(w, \eta^r_{cM}) P(w, t)] + \bar{\Gamma}\partial_w^2 P(w, t), \tag{15}$$

with $\bar{\Gamma} = \Gamma/2$. This drift-diffusion equation for the probability density can be written in flux form:

$$\partial_t P(w,t) = -\partial_w \mathcal{J}, \tag{16}$$

where

$$\mathcal{J} = -\partial_w \mathcal{U}(w, \eta^r_{cM}) P(w,t) - \bar{\Gamma} \partial_w P(w,t) = -\bar{\Gamma} e^{-\mathcal{U}(w,\eta^r_{cM})/\bar{\Gamma}} \partial_w \left( P(w,t) e^{\mathcal{U}(w,\eta^r_{cM})/\bar{\Gamma}} \right) \tag{17}$$

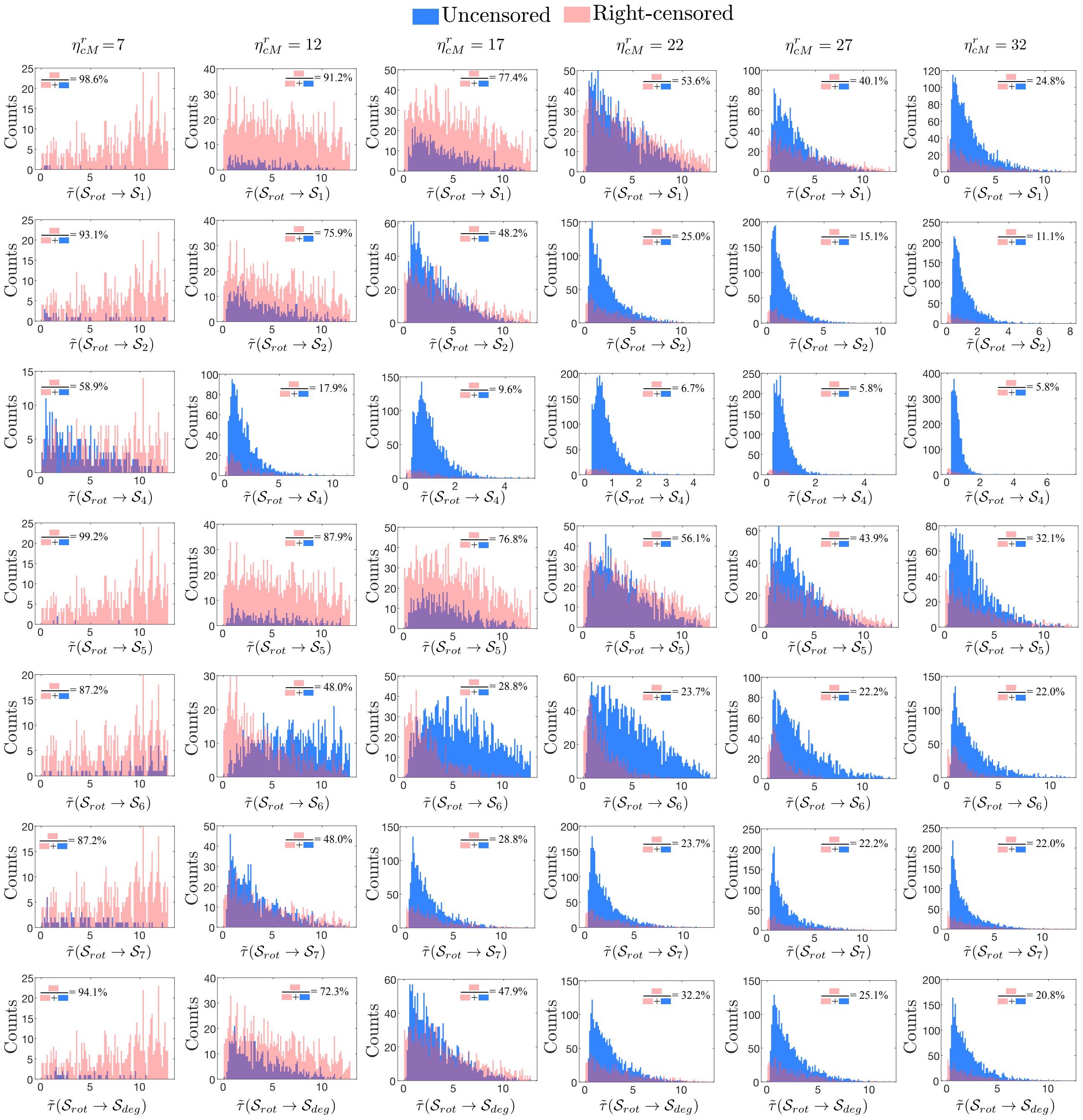


**Figure S13:** Distribution of first passage times from the rotating state to the rest of states for different confinement values $\eta^r_{cM}$.

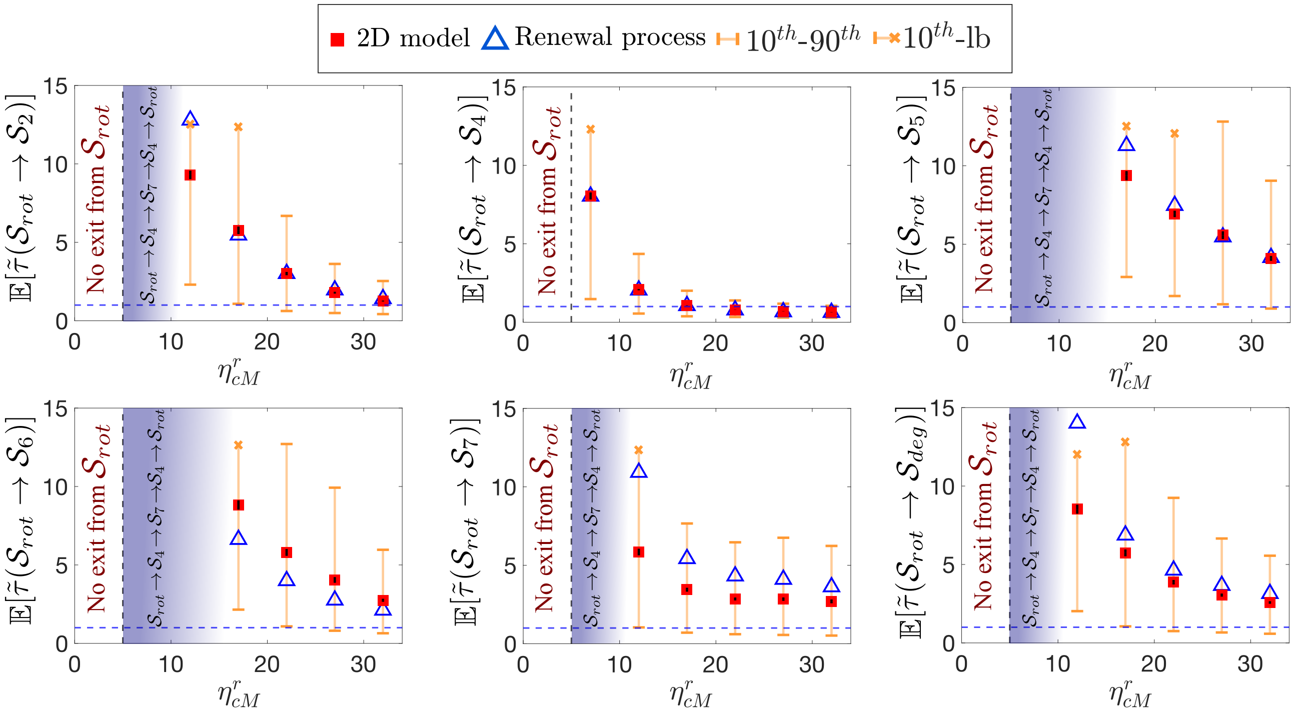


**Figure S14:** Mean first passage times from the rotating state to the rest of the states except for $\mathcal{S}_1$ (see Main text), computed using the 2D phase field model (squares) and the semi-Markov renewal process (triangles) as a function of $\eta^r_{cM}$. The orange whiskers correspond to the $10^{th}$ and $90^{th}$ percentiles of the first-passage time distributions. The orange (x) symbols indicates the lower bound (maximum first passage time observed) when a $90^{th}$ cannot be estimated. Bootstrap standard errors (500 trajectory resamples) of all the $\mathbb{E}[\tilde{\tau}(\mathcal{S}_{rot} \to \mathcal{S}_{j\neq 1})]$ are indicated by red whiskers

.

can be interpreted as a probability flux over the potential barrier. Therefore, this flux is related to $\mathcal{R}$ by the simple relationship $\mathcal{R}p = \mathcal{J}$, where $p$ is the probability of being near the potential minimum ($A^s_b$):

$$p = \int_{w_1}^{w_2} P(w', t) dw', \tag{18}$$

where $w_1$ and $w_2$ are arbitrary points inside the potential. If the barrier height, $\Delta\mathcal{U} = \mathcal{U}(A^u_b) - \mathcal{U}(A^s_b)$, is much larger than $\bar{\Gamma}$, which holds for sufficiently weak confinements and the noise amplitudes considered in our study (Fig. 3C and Table S1), the probability distribution near $A^s_b$ can be approximated by the stationary distribution:

$$P(w, t) = P(A^s_b, t) e^{-[\mathcal{U}(w, \eta^r_{cM}) - \mathcal{U}(A^s_b, \eta^r_{cM})]/\bar{\Gamma}}, \tag{19}$$

which is obtained from Eq. (17) with $\mathcal{J} = 0$ and normalizing by the probability distribution at $A^s_b$. To obtain an expression for $\mathcal{J}$ and estimate $\mathcal{R}$, we assume that the system is near steady state, and thus, one can consider the limit $\partial_t P(w, t) \approx 0$, which implies that the flux $\mathcal{J}$ is independent of $w$, and integrate Eq. (17) from $A^s_b$ to $A^a_b$, to obtain

$$\mathcal{J} = \frac{\bar{\Gamma} P(A^s_b) e^{-\mathcal{U}(A^s_b, \eta^r_{cM})/\bar{\Gamma}}}{\int_{A^s_b}^{A^a_b} e^{\mathcal{U}(w', \eta^r_{cM})/\bar{\Gamma}} dw'}, \tag{20}$$

where $A^a_b$ is an arbitrary point outside the potential in which $P(A^a_b) \approx 0$ and $A^a_b > A^u_b$. Combining the expressions for $\mathcal{J}$ and $p$, considering the stationary probability distribution at $A^s_b$, the Kramers escape rate reads

$$\mathcal{R} = \frac{1}{\bar{\Gamma}} \int_{w_1}^{w_2} e^{-\mathcal{U}(w', \eta^r_{cM})/\bar{\Gamma}} dw' \int_{A^s_b}^{A^a_b} e^{\mathcal{U}(w', \eta^r_{cM})/\bar{\Gamma}} dw'. \tag{21}$$

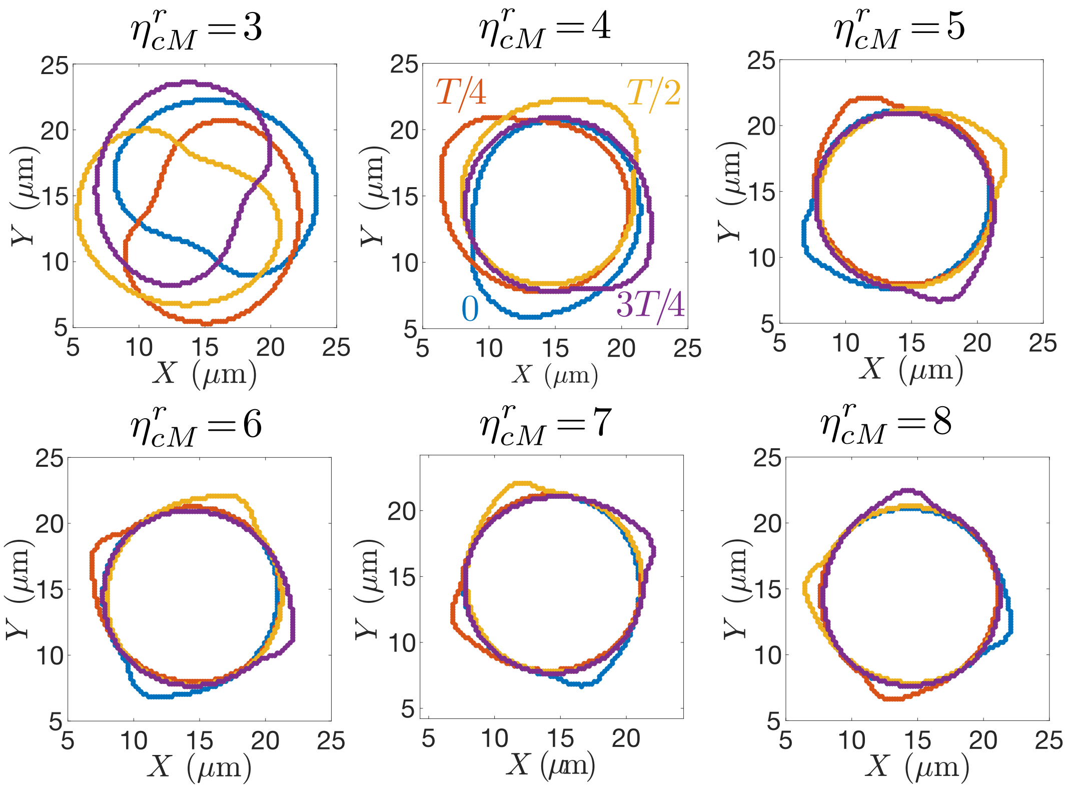


**Figure S15:** Temporal evolution during one full rotation, shown every $T/4$, of the cell's interface $\varphi_c = 1/2$ near the transition ($\eta^r_{cM} \approx 5$) from intermediate to weak confinement regime. Below the critical confinement $\eta^r_{cM} \approx 5$, the protrusion becomes large enough to contract the entire cell, resulting in rigid-body motion. These simulations are noiseless ($\Gamma = 0$).

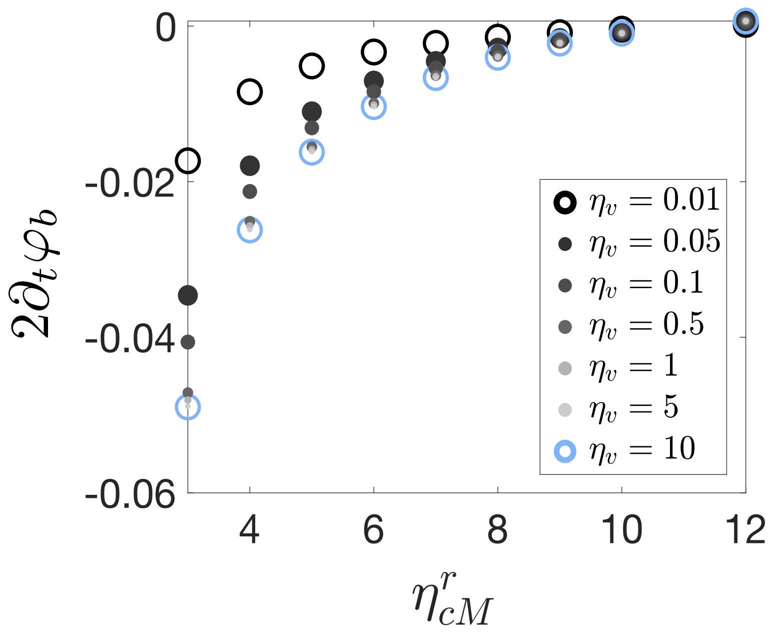


**Figure S16:** Changes in the contraction effect at the back of the cell, $\partial_t\varphi_b/\varphi_c = 2\partial_t\varphi_b$, for different values of the strength of the restoring-size force $\eta_v$ over a range of confinements.

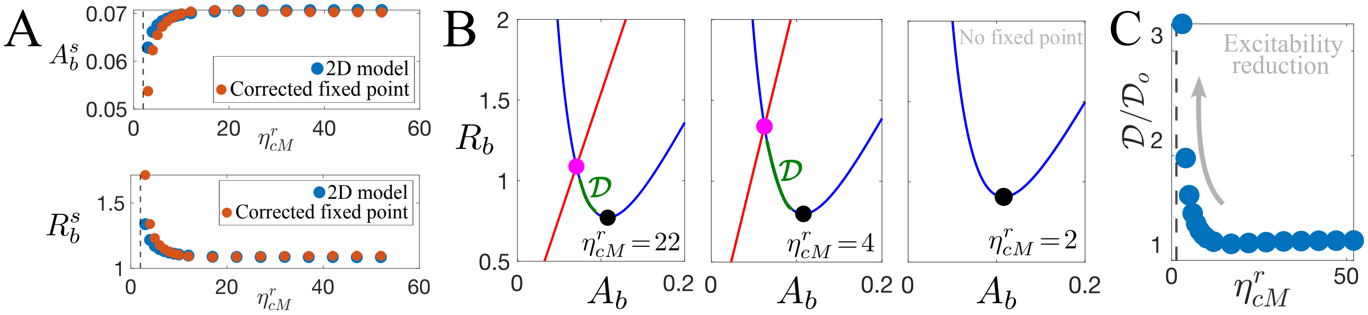


**Figure S17:** Activator-inhibitor dynamics at the back of the cell. (A) Comparison between the back fixed point $\{A_b^s, R_b^s\}$ obtained from numerics (2D model) and from Eq. (12) for different values of $\eta_{cM}^r$. (B) Phase space visualization of the fixed point shift as confinement weakens. The distance $\mathcal{D}$ between the fixed point (magenta) and the minimum of the cubic nullcline (black) is highlighted in green. (C) Growth of $\mathcal{D}$ until the fixed point disappears ($\eta_{cM}^r \approx 2.3$). The distance $\mathcal{D}_o$ is measured under homogeneous conditions ($\partial_t \phi_b = 0$).

One can solve both integrals in Eq. (21) using Taylor expansions, up to second order, of the potential well $\mathcal{U}$ around $A_b^s$ for the first integral, and around $A_b^u$ for the second integral. The rate is then given by

$$\mathcal{R} = \frac{1}{2\pi}\sqrt{\left.\frac{d^2\mathcal{U}}{dA_b^2}\right|_{A_b^s}\left|\left.\frac{d^2\mathcal{U}}{dA_b^2}\right|_{A_b^u}\right|}\, e^{-\Delta\mathcal{U}/\bar{\Gamma}}, \tag{22}$$

where

$$\frac{d^2\mathcal{U}}{dA_b^2} = b + (d_1 + d_2 R_b^s) - 2|\partial_t \varphi_b| + k_a - \frac{k_a K_a^2}{(K_a^2 + A_b^2)^2}\left(2aA_b + K_a^2 - A_b^2\right). \tag{23}$$

Finally, the inverse of $\mathcal{R}$ can be interpreted as the mean escape time $\mathcal{T}$ from the potential well. Fig. 3D of the main text shows that this escape time increases as confinement weakens, consistent with the reduction of excitability triggered by contraction.

### XI. Experimental setup

MCF10A mammary epithelial cells tranduced with LifeAct-GFP and H2B-mCherry were cultured in growth factor reduced matrigel (Ref: 356231, Phenol red free, Corning). Briefly, MCF10A cells, at a seeding density of 500 cells per $\mu$L, were mixed with Matrigel at ratios ranging from 30% to 100% and cast into a 96-well glass-bottom plate (Matek Corporation). The gels were allowed to crosslink at 37°C for 45 minutes. After 2 h of culture, 35 $\mu$m Z-stacks were acquired every 20 min for a total duration of 16 h using a Nikon TiE inverted microscope. The captured timelapses were then post processed using Bitplane Imaris software.

The Z-stacks were manually examined to assess the presence of cell rotation, defined as visible chiral motion persisting for $0.25T$ and accompanied by a distinctive F-actin patch on the cell surface. The identification was independently performed by three users, resulting in perfect agreement for Matrigel concentrations of 50%, 60% and 80%, and slight variable agreement for 70% and 100%. This variability is reported as standard deviations in Fig. 4D of the main text.

### XII. Equivalence between weak confinement and extra free space

In our model, confinement weakening is analogous to increasing the effective cell-ECM distance, allowing protrusions to develop more easily. To illustrate this, we perform a weakly nonlinear analysis around the steady state solution of a one-dimensional representation of the cell (similar to the lower panel in Fig. S1A). We assume that $\int \varphi_c d\vec{r} - V_c \ll 1$, and thus focus only on the effect of confinement and protrusions:

$$\partial_T \varphi_c = \partial_{XX}\varphi_c - \frac{\gamma}{\epsilon^2}\frac{\delta \mathcal{G}[\varphi_c]}{\delta \varphi_c} + \left(\frac{\eta_p A - \eta_{cM}^r \varphi_M}{\sqrt{\gamma}}\right)|\partial_X \varphi_c|, \tag{24}$$

where space and time have been rescaled as $x = \sqrt{\gamma}X$ and $t = \xi T$. Additionally, we assume that the activator dynamics is much faster than the mechanical deformations of the cell membrane such that we can ignore the temporal dependence of $A$ in Eq. (24). Because of the symmetry and the simplified size restoring force, we analyze the dynamics locally at one end of the 1D-cell; without loss of generality, we choose the right side of the 1D profile (cf. lower panel in Fig. S1A). In the absence of confinement ($\eta_{cM}^r = 0$) and protrusions ($\eta_p = 0$), and at the Maxwell point of the variational system, the 1D phase field is characterized by the homogeneous profile $\varphi_c^o = \{1 - \tanh[3\sqrt{\gamma}(X - X_c)/\epsilon]\}/2$, where $X_c$ is the cell boundary. A similar one-dimensional profile is used for the static ECM: $\varphi_M = \{1 + \tanh[q(X - X_M)]\}$, where $X_M$ is the ECM boundary, and for simplicity we choose $q = 3\sqrt{\gamma}/\epsilon$. The aim is to track the movement of the cell boundary through the first nonlinear correction $W$ introduced by perturbations from a protrusion and confinement: $\varphi_c(X,T) = \varphi_c^o(X - X_c(T)) + W$. Substituting this ansatz in Eq. (24) yields, at $\mathcal{O}(W)$,

$$-\partial_T X_c \partial_X \varphi_c^o = (\partial_{XX} - \varphi_c^o + 3\varphi_c^{o2} - 2\varphi_c^{o3})W + \left(\frac{\eta_p A - \eta_{cM}^r \varphi_M}{\sqrt{\gamma}}\right)|\partial_X \varphi_c^o|, \tag{25}$$

where we assume $\partial_T X_c \sim W \sim \eta_{cM}^r \sim \eta_p$ are small. This equation is linear in $W$, $\mathcal{L}W = b$, with

$$b = -\frac{(\eta_p A - \eta_{cM}^r \varphi_M)}{\sqrt{\gamma}}|\partial_X \varphi_c^o| - \partial_T X_c \partial_X \varphi_c^o. \tag{26}$$

Under the usual inner product $\langle f, g\rangle = \int fg dX$, the linear operator is self-adjoint, $\mathcal{L} = \mathcal{L}^T$, with kernel $\partial_X \varphi_c^o$. Then, Eq. (25) is solvable if and only if $\langle \partial_X \varphi_c^o, b\rangle = 0$. This solvability condition leads to the following expression for the one-dimensional velocity of the cell boundary:

$$\begin{aligned}\partial_T X_c &= \frac{\eta_{cM}^r}{\sqrt{\gamma}} \frac{\langle \varphi_M |\partial_X \varphi_c^o|, \partial_X \varphi_c^o\rangle}{\langle \partial_X \varphi_c^o, \partial_X \varphi_c^o\rangle} - \frac{\eta_p}{\sqrt{\gamma}} \frac{\langle A|\partial_X \varphi_c^o|, \partial_X \varphi_c^o\rangle}{\langle \partial_X \varphi_c^o, \partial_X \varphi_c^o\rangle} \\ &= -\frac{3\eta_{cM}^r}{8\sqrt{\gamma}}\left(\frac{4}{3} + \int_{-\infty}^{\infty} \mathrm{sech}\left(z - 3\frac{\sqrt{\gamma}}{\epsilon}\Delta\right)\tanh(z) dz\right) + \frac{3\eta_p}{8\sqrt{\gamma}}\langle A|\partial_X \varphi_c^o|, \partial_X \varphi_c^o\rangle,\end{aligned} \tag{27}$$

where $z = X - X_c$ and $\Delta = X_c - X_M$. The last term, proportional to $\eta_p$, is always positive and gives the first order contribution of the protrusive force. Let us focus on the first term in Eq. (27), proportional to $\eta_{cM}^r$, which gives the first order contribution to confinement. The integral term, denoted $\mathcal{I}$, is difficult to evaluate analytically but can be interpreted straightforwardly. When $\Delta = 0$, $\mathcal{I}$ is zero by parity. For $\Delta \neq 0$, the sign of $\Delta$ controls the sign of the integral, $\mathrm{sgn}(\mathcal{I}) = \mathrm{sgn}(\Delta)$. Therefore, when extra space is available to the cell ($\Delta < 0$), the term in parentheses in Eq. (27) decreases, effectively weakening the confinement. Reintroducing the spatial and temporal scales, Eq. (27) becomes

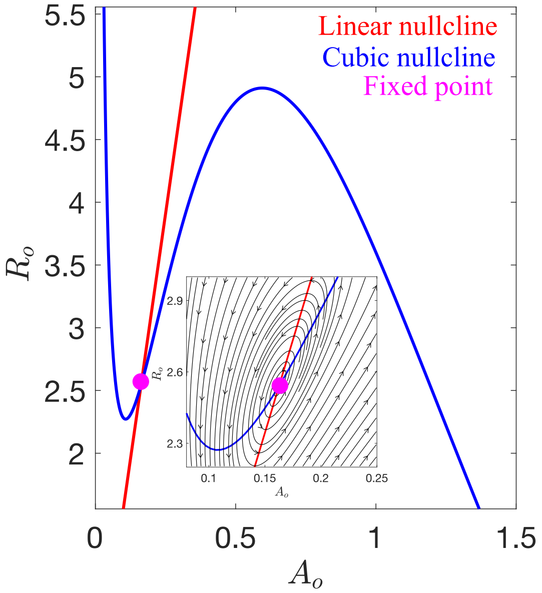


**Figure S18:** Phase space of the zero-dimensional version of the activator-inhibitor dynamics in Eq. (2), given by $\{A_o, R_o\}$, in the oscillatory regime. The inset shows the structure of the phase flows around the fixed point (repeller).

from the minimum of the cubic nullcline—quantified by the distance $\mathcal{D}$ (see Section IX—the reduction in excitability can suppress activator nucleations and therefore coherent rotations.

We vary the parameters $\{a, b, d_1, d_2, k_a, K_a, c_1, c_2\}$ one by one within $\pm 50\%$ of their original value (Table S1) and verify that any such variation produces coherent rotations provided that $\mathcal{D} \lesssim 0.32$ (blue shaded regions in Fig. S19). For larger $\mathcal{D}$, the system is unable to produce activator pulses (pink shaded regions in Fig. S19). Parameter changes can also induce a bifurcation from the excitable to the oscillatory regime (green shaded regions in Fig. S19). The parameter $\tau$ in Eq. (2), which sets the time scale of the inhibitor, does not affect the location of the intersection in phase space. However, it cannot be arbitrarily small (i.e., fast time scale), as this can prevent the emergence of activator pulses. We have checked that coherent rotations are still present when varying $\tau$ within $\pm 25\%$ of its original value.

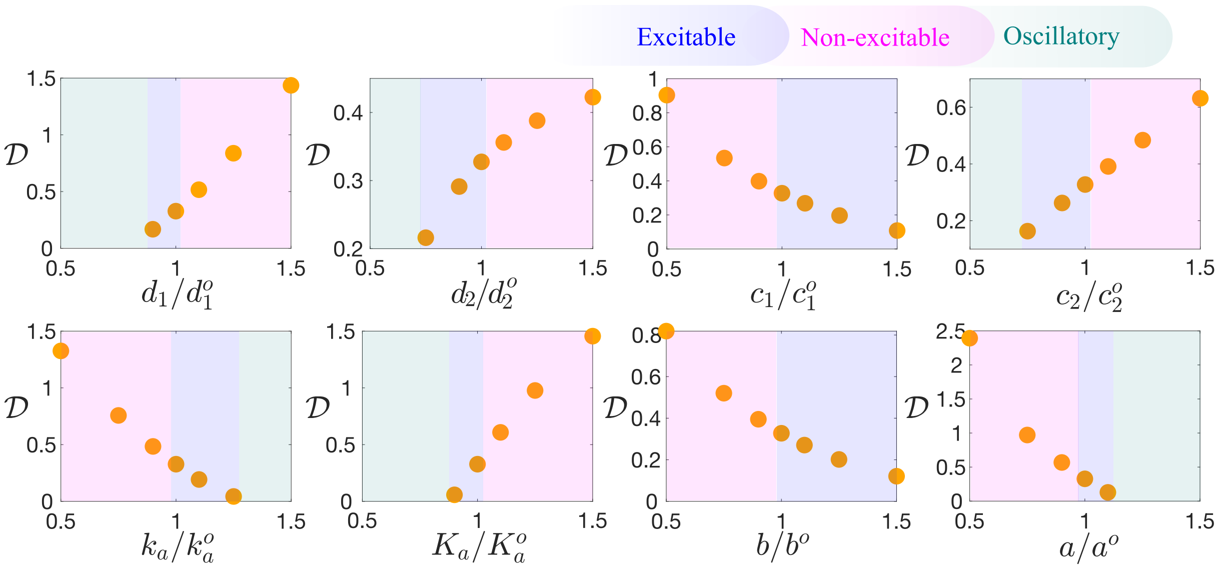


**Figure S19:** Variations in $\mathcal{D}$ as a function of eight parameters controlling the wave dynamics in the activator-inhibitor system of Eq. (2), and in the weak confinement regime ($\eta^r_{cM} = 4$). The superscript $^\circ$ denotes the original parameter value in Table S1.

**XV. Supplemental figures for Section III.D of the main text: Fig. S20 and Fig. S21**

Figs. S20 and S21 illustrate how the mean dwell time in the rotating state depends on variations in the diffusion coefficient $D_A$ and the friction coefficient $\xi$ across the confinement values analyzed in the main text.

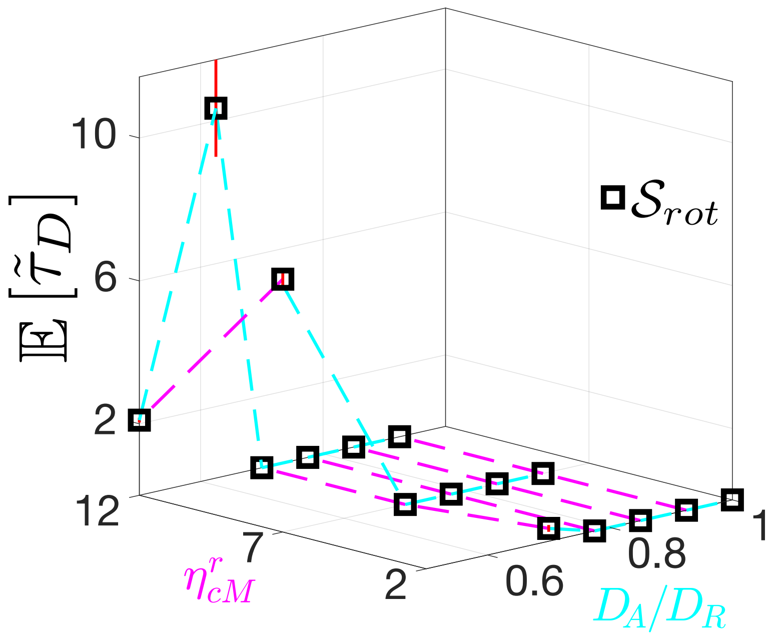


**Figure S20:** Phase diagram in $\eta^r_{cM} - D_A/D_R$ space showing the computable mean dwell times in the rotating state.

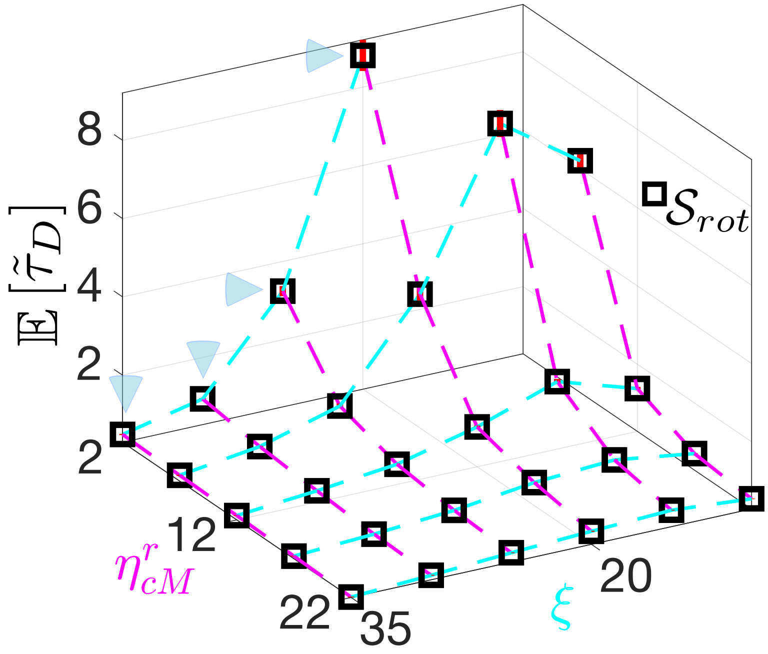


**Figure S21:** Phase diagram in $\eta^r_{cM} - \xi$ space showing the computable mean dwell times in the rotating state. The blue triangles indicate measurable mean dwell times at $\eta^r_{cM} = 2$ (exit from $\mathcal{S}_{rot}$ is possible).

**XVI. Video captions**

**Video S1**: Numerical integration of Eqs. 2 and 4 in 3D with $\eta^r_{cM} = 32$. The color code indicates the normalized activator $A/\max(A)$. The total time of integration is $66T$.

**Video S2**: Numerical integration of Eqs. 2 and 4 in 3D with $\eta^r_{cM} = 5.3$. The color code indicates the normalized activator $A/\max(A)$. The total time of integration is $66T$.

**Video S3**: Numerical integration of Eqs. 2 and 4 in 3D with $\eta^r_{cM} = 4$. The color code indicates the normalized activator $A/\max(A)$. The total time of integration is $66T$.

**Video S4**: Interface dynamics ($\varphi_c = 1/2$) shown across an arbitrary plane, extracted from a segment of **Video S1**. The color code indicates the normalized activator $A/\max(A)$.

**Video S5**: Interface dynamics ($\varphi_c = 1/2$) shown across a plane $\{p'_1, p'_2\}$, extracted from a segment of **Video S2**. The plane is chosen to coincide with an observed episode of coherent rotation. The color code indicates the normalized activator $A/\max(A)$.

**Video S6**: Interface dynamics ($\varphi_c = 1/2$) shown across a plane $\{p_1, p_2\}$, extracted from a segment of **Video S3**. The plane is defined by the cell symmetry axis after reaching for the first time the state of coherent rotation. The color code indicates the normalized activator $A/\max(A)$.

**Video S7**: Numerical integration of Eqs. 2 and 4 in 2D with $\eta^r_{cM} = 7$ exhibiting the representative $\mathcal{S}_i$ states and their transitions. The color code indicates the activator $A$ in RGB scale. The total time of integration is $13T$. The green circles show pulse positions (see Section V).

**Video S8**: F-actin dynamics on MCF10A cell surface in 100% Matrigel showing states $\mathcal{S}_1$, $\mathcal{S}_2$, $\mathcal{S}_{rot}$ and $\mathcal{S}_6$. F-actin is visualized with GFP-tagged LifeAct (green) and the nucleus is visualized with mCherry-tagged H2B (orange).

**Video S9**: Rotating ($\mathcal{S}_{rot}$) MCF10A cell in 100% Matrigel. F-actin is visualized with GFP-tagged LifeAct (green) and the nucleus is visualized with mCherry-tagged H2B (orange).

**Video S10**: F-actin dynamics on MCF10A cell surface in 70% Matrigel showing state $\mathcal{S}_5$. F-actin is visualized with GFP-tagged LifeAct (green) and the nucleus is visualized with mCherry-tagged H2B (orange).

**Video S11**: Translational non-rotating MCF10A cell in 100% Matrigel. F-actin is visualized with GFP-tagged LifeAct (green) and the nucleus is visualized with mCherry-tagged H2B (orange).

**Video S12**: Extract of a numerical integration of Eqs. 2 and 4 in 2D with $\eta^r_{cM} = 7$ showing a short episode with four pulses. The color code indicates the activator $A$. The green circles show pulse positions (see Section V).

**Video S13**: Extract of a numerical integration of Eqs. 2 and 4 in 2D with $\eta^r_{cM} = 7$ showing the *back and forth* mode in the oscillatory regime ($d_1 = 1.5$). The color code indicates the activator $A$.

**Video S14**: Extract of a numerical integration of Eqs. 2 and 4 in 3D with $\eta^r_{cM} = 9$ showing the target-wave pattern in the oscillatory regime ($d_1 = 1.35$). The color code indicates the activator $A$.
